# The critical role of natural history museums in advancing eDNA for biodiversity studies: a case study with Amazonian fishes

**DOI:** 10.1101/2021.04.18.440157

**Authors:** C. David de Santana, Lynne R. Parenti, Casey B. Dillman, Jonathan A. Coddington, D. A. Bastos, Carole C. Baldwin, Jansen Zuanon, Gislene Torrente-Vilara, Raphaël Covain, Naércio A. Menezes, Aléssio Datovo, T. Sado, M. Miya

## Abstract

Ichthyological surveys have traditionally been conducted using whole-specimen, capture-based sampling with varied, but conventional fishing gear. Recently, environmental DNA (eDNA) metabarcoding has emerged as a complementary, and possible alternative, approach to whole-specimen methodologies. In the tropics, where much of the diversity remains undescribed, vast reaches continue unexplored, and anthropogenic activities are constant threats; there have been few eDNA attempts for ichthyological inventories. We tested the discriminatory power of eDNA using MiFish primers with existing public reference libraries and compared this with capture-based methods in two distinct ecosystems in the megadiverse Amazon basin. In our study, eDNA provided an accurate snapshot of the fishes at higher taxonomic levels and corroborated its effectiveness to detect specialized fish assemblages. Some flaws in fish metabarcoding studies are routine issues addressed in natural history museums. Thus, by expanding their archives to include eDNA and adopting a series of initiatives linking collection-based research, training and outreach, natural history museums can enable the effective use of eDNA to survey Earth’s hotspots of biodiversity before taxa go extinct. Our project surveying poorly explored rivers and using DNA vouchered archives to build metabarcoding libraries for Neotropical fishes can serve as a model of this protocol.

## Introduction

Historical ichthyological surveys in freshwater ecosystems globally were conducted with whole-specimen, capture-based sampling using conventional fishing methods such as gill nets, cast nets, hook and line, dipnets, seines, and rotenone – the last a chemical ichthyocide. Although the use of ichthyocides is considered advantageous in the tropics where unknown quantities of diversity remain to be described (e.g.,^1^), collecting fishes with rotenone has been banned as a sampling method in many regions due to its extraordinary power to kill fishes and associated fauna (e.g.,^2^). Capture-based methods other than rotenone are less powerful, especially for collecting small cryptobenthic species, and all capture-based methods may result in low capture rates in hard-to-sample environments, such as rapids, waterfalls, and deep-water reaches. Yet, by undersampling we increase the likelihood of overlooking, and in the sense of the biodiversity crisis globally missing, heretofore unaccounted-for species diversity.

Recently, DNA barcodes from environmental samples (eDNA), a non-invasive and quickly developing methodology that captures genetic material of multiple organisms, has emerged as a complementary, and a possible alternative approach, to repeated whole-specimen capture methods. Universal eDNA metabarcoding primers based on short variable DNA regions (typically ribosomal RNA – 12S rRNA, e.g.,^3^) were developed to detect multiple species of fishes through next-generation sequencing (NGS) of free DNA molecules that exist in nature (e.g., lost scales, excrement, and mucosal secretions in the water^4,5,6^).

The Amazon rainforest and basin maintain the most diverse riverine ichthyofauna on Earth, with more than 2,700 species classified in 18 orders and 60 families^7,8^. Such numbers are underestimates as many undescribed taxa await discovery and formal description^9^. The evolution of this fauna, one-fifth of the world’s freshwater fishes^10^, dates to at least the upper Cretaceous and lower Cenozoic, ~ 120-150 million years before present (mybp^11^). The Characiphysae — catfishes (Siluriformes), piranhas and allies (Characiformes) and electric fishes (Gymnotiformes)— represent more than 75% of the fishes in Amazonian aquatic ecosystems^7^. That overwhelming fish diversity is also represented by, among other taxa, cichlids (Cichliformes), killifishes (Cyprinodontiformes), river stingrays (Myliobatiformes), pufferfishes (Tetraodontiformes), and silver croakers (Perciformes)^7,12,13^. Over time these fishes have diversified under a wildly varied set of environmental conditions to inhabit myriad aquatic systems^14^. In contrast to the proposed ancient age of those lineages, most species-level diversification is hypothesized to have occurred relatively recently, less than 10 mybp (e.g.,^13,15^).

Accurate and thorough sampling is the critical first step towards a more complete knowledge of biodiversity, a path that also requires the proper identification of collected samples. Specimens of Amazonian fishes have been identified almost exclusively based on morphology, but given that molecular evolutionary rates can far outpace divergence in phenotypes, recent studies that integrate molecular and morphological data have greatly improved our understanding of species diversity, including that of fishes^15,16^. DNA barcoding – which typically uses the mitochondrial COI (Cytochrome Oxidase subunit I) gene to identify candidate species^17^ – is now a common molecular method used in taxonomic studies of fishes and has been valuable in revealing cryptic species diversity and in helping to resolve complex taxonomic issues^15,18,19^.

Accordingly, the demand for samples appropriate for DNA barcoding, i.e., properly preserved and vouchered in ichthyological collections by the scientific community, has increased significantly. This demand is correlated directly with efforts to collect DNA-worthy samples during biodiversity surveys along with museum vouchers and has become a common practice among scientists worldwide^20,21,22^. Concomitant with sampling and curating efforts, new public platforms have been created to help close gaps in shared sample information (e.g., Global Genome Biodiversity Network^23^) and facilitate access to the sequences (e.g., BOLD). In contrast to this trend, historically few efforts to collect a substantial number of tissue samples during ichthyological surveys – possibly because of the lack of infrastructure to maintain such a collection – results in a lack of robust reference libraries for Amazonian fishes (e.g.,^24^). In addition, although Genbank often is a reliable resource^25^, several samples of Amazonian fishes are poorly identified in GenBank, and somelack properly preserved voucher specimens – a problem that extends to other fishes as well (e.g.,^26,27^).

Most metabarcoding inventories of freshwater fishes have been conducted in temperate habitats with well-characterized species diversity^28^. There have been only a few attempts to use eDNA metabarcoding in ichthyological surveys in the Neotropical region^29,30,31,32^, an area where understanding species-level diversity is more complex. For example, ^30^ built 12S eDNA metabarcoding primers based on a reference library for over 130 species known to occur in the rivers and streams of the French Guiana, and the eDNA results were compared with capture-based sampling methodologies. They recovered a similar number of species, with a partial match to species identification, using both capture-based and eDNA approaches. Conversely, ^32^ used MiFish primers^3^ in three localities in the central Amazon and suggested that a new approach would be necessary to evaluate the Neotropical fish fauna using eDNA metabarcoding.

Despite the problems inherent in the development of new methodologies, eDNA technology and bioinformatics is evolving at accelerated rates and will soon play a central role in the inventory of fish diversity^6,33,28,34^. Freshwater aquatic ecosystems, many of which are poorly explored, are under severe and fast-pacing threat due to anthropogenic activities^35^. Thus, the next decade or so will be pivotal to survey these habitats to secure vouchers, DNA, and eDNA samples to build reference libraries and archive the samples as well as to engage society in protection and preservation as these environments reach their tipping point. Natural history museums are the sound common ground where key flaws and gaps in those two inversely proportional trends can be addressed and filled. Here, we tested the discriminatory power of the MiFish primers using the existing public reference libraries by surveying two distinct ecosystems, river and stream, during a scientific expedition to the heretofore largely unexplored Javari River basin in Brazil-Peru-Colombia border. The results of eDNA analysis were compared with the capture-based methodology and are discussed in the context of the critical role of natural history museums in the development of eDNA metabarcoding as a tool for biodiversity studies.

## Results

### 1. Ichthyological survey – Capture-based sampling (CBS)

In total, 443 species classified in 236 genera, 49 families, and 15 orders were collected using traditional methods from 46 stations in multiple environments during the Javari River expedition (Table S1). Among these collections are over 60 species that are new to science.

More specifically, in the three stations sampled using traditional and eDNA methodologies (Figure 1), we collected the following: 145 species, 101 genera, 32 families and nine orders in the main Javari River (station 1); 56 species, 38 genera, 21 families, and six orders in a stream (station 2); and 67 species, 58 genera, 27 families, and seven orders in the Quixito River (station 3; Table S1).

**Figure 1.**
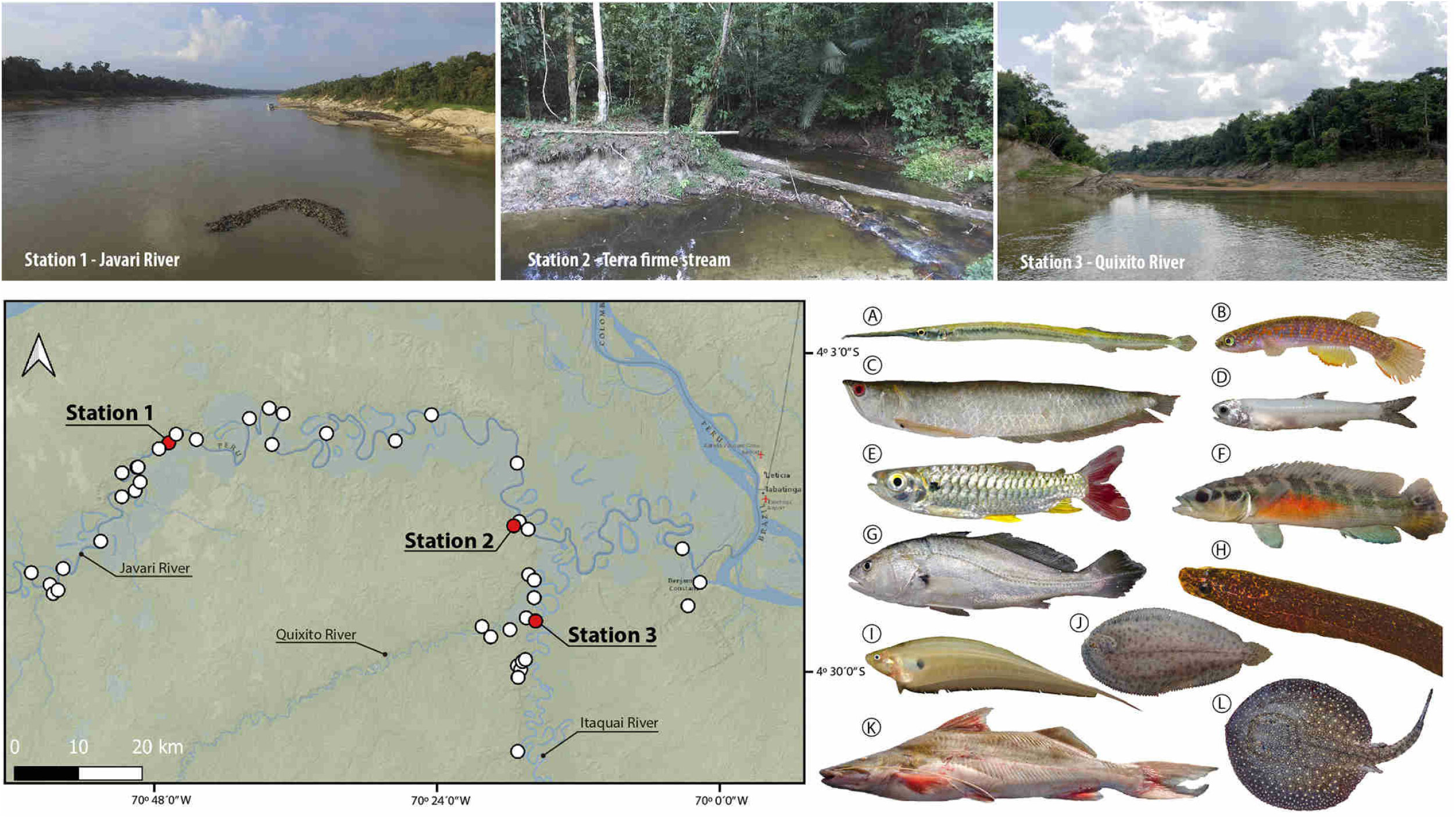
Map of Javari River basin showing 46 sampling stations (white and red dots). Red dots represent stations in two distinct ecosystems (River: stations 1 – Javari, 3 – Quixito; Stream: station 2 – Terra firme stream) sampled by capture and molecular based methodologies. In the map one dot can represent more than one station. Illustration of 12 orders detected by capture-based sampling (CBS) and molecular based sampling (MBS) in the three stations: (A) Beloniformes – *Potamorrhaphis guianensis*, INPA-ICT 055254, station 2(CBS); (B) Cyprinodontiformes – *Laimosemion* sp., INPA-ICT 056039, station 2(CBS); (C) Osteoglossiformes – *Osteoglossum bicirrhosum*, INPA-ICT 056354; (D) Clupeiformes – *Anchoviella jamesi*, INPA-ICT 055391, stations 1&3(CBS); (E) Characiformes – *Chalceus erythrurus*, INPA-ICT 055360, stations 1(CBS, MBS), 2(MBS); (F) Cichliformes – *Crenicichla reticulata*, INPA-ICT 055413, station 1(CBS); (G) Perciformes – *Plagioscion squamosissimus*, INPA-ICT 055328, stations 1(CBS, MBS), 3(CBS); (H) Synbranchiformes – *Synbranchus* sp., INPA-ICT 055815, station 2(MBS); (I) Gymnotiformes – *Eigenmannia limbata*, INPA-ICT 055420, stations 1 & 3(CBS, MBS), 2(MBS); (J) Pleuronectiformes – *Apionichthys nattereri*, INPA-ICT 055487, stations 1(CBS, MBS), 3(MBS); (K) Siluriformes –*Brachyplatystoma vaillantii*, INPA-ICT 056703, station 1(MBS); (L) Myliobatiformes – *Potmotrygon scobina*, INPA-ICT 055553.

#### eDNA data analyses and assessment of taxonomic resolution of public reference database (Molecular-based Sampling -- MBS)

A total of 1,903,160 reads was assigned to the 11 libraries (station 1 = 5 libraries; station 2 = 5 libraries; station 3 = 1 library), and the number of raw reads for each library ranged from 135,818 to 213,952 with an average of 173,015 reads (Table S2). The final reference database with 1,671,871 fish reads (99.7% of the denoised reads) yielded 222 MOTUs assigned to 104 genera, 41 families, and 9 orders of fishes (Figure 2 and Tables S3, S4).

**Figure 2.**
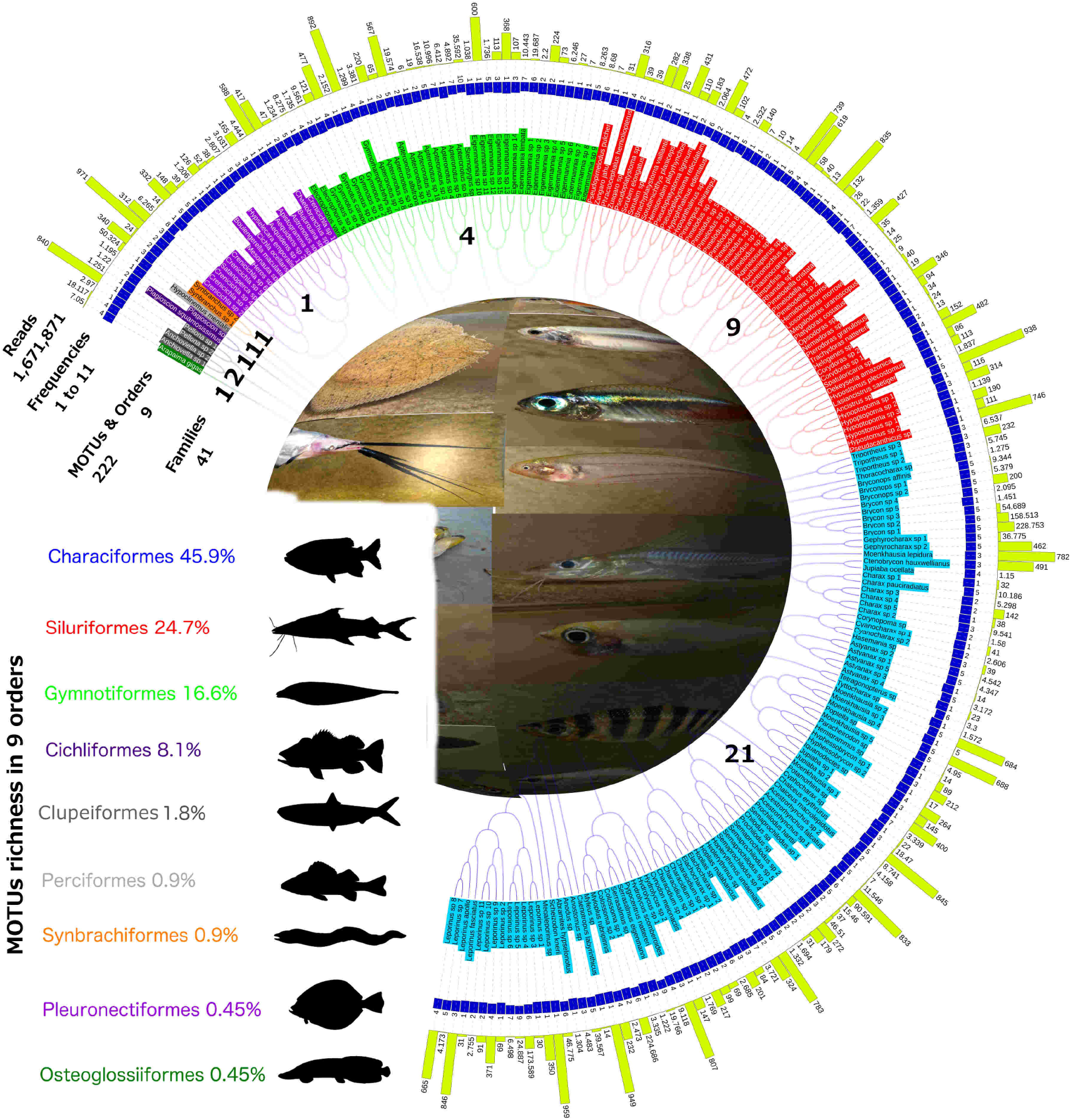
Illustrative cladogram with reads and frequencies for each of 222 metabarcoding operational taxonomic units (MOTUs) and reference sequences included in nine orders and 41 families detected by 11 eDNA samples in the Javari River basin. Color highlighting MOTUs names corresponds to each of the nine orders. In the left side, species richness, key color, and general bauplan silhouettes for each order. At the center, spherical view of species diversity detected by eDNA.

Matching sequences identity of >98.5% for 58 species (26%) of 222 species detected by eDNA were found in the reference library database (Table 1). It represented 36.3% of Siluriformes; 27.1% of Characiformes; 11.7% of Cichliformes; and 10.8% of Gymnotiformes in the most species-rich orders detected by eDNA. From these, six species (10.3%) were identified as “sp.” in reference libraries. Only 17species (7.6%) were also identified in the CBS (Table 1).

### 2. Species composition among eDNA samples: distinguishing between river versus stream-dwelling communities

Six eDNA samples were collected in the river (stations 1 and 3) and five in the stream (station 2), and a clear split is seen between these two fish communities (Figure 3). The number of species detected per sample ranged from 33 to 87 (for details see Supp information) with an abrupt differentiation between species composition in the stream (samples 1-5) and river samples (samples 6-11), as detected by the Pearson correlation coefficients (Figure 3A). That is, stream and river-dwelling communities are clearly distinct on the species composition Habitat axis. Pearson coefficients are varying from 0.5 to 1.0 in stream versus 0.0 to −0.5 in river. Thus, species composition is more similar within each community, except for a clear distinction between the river assemblages at Javari (samples 6-10) and Quixito Rivers (sample 11).

**Figure 3.**
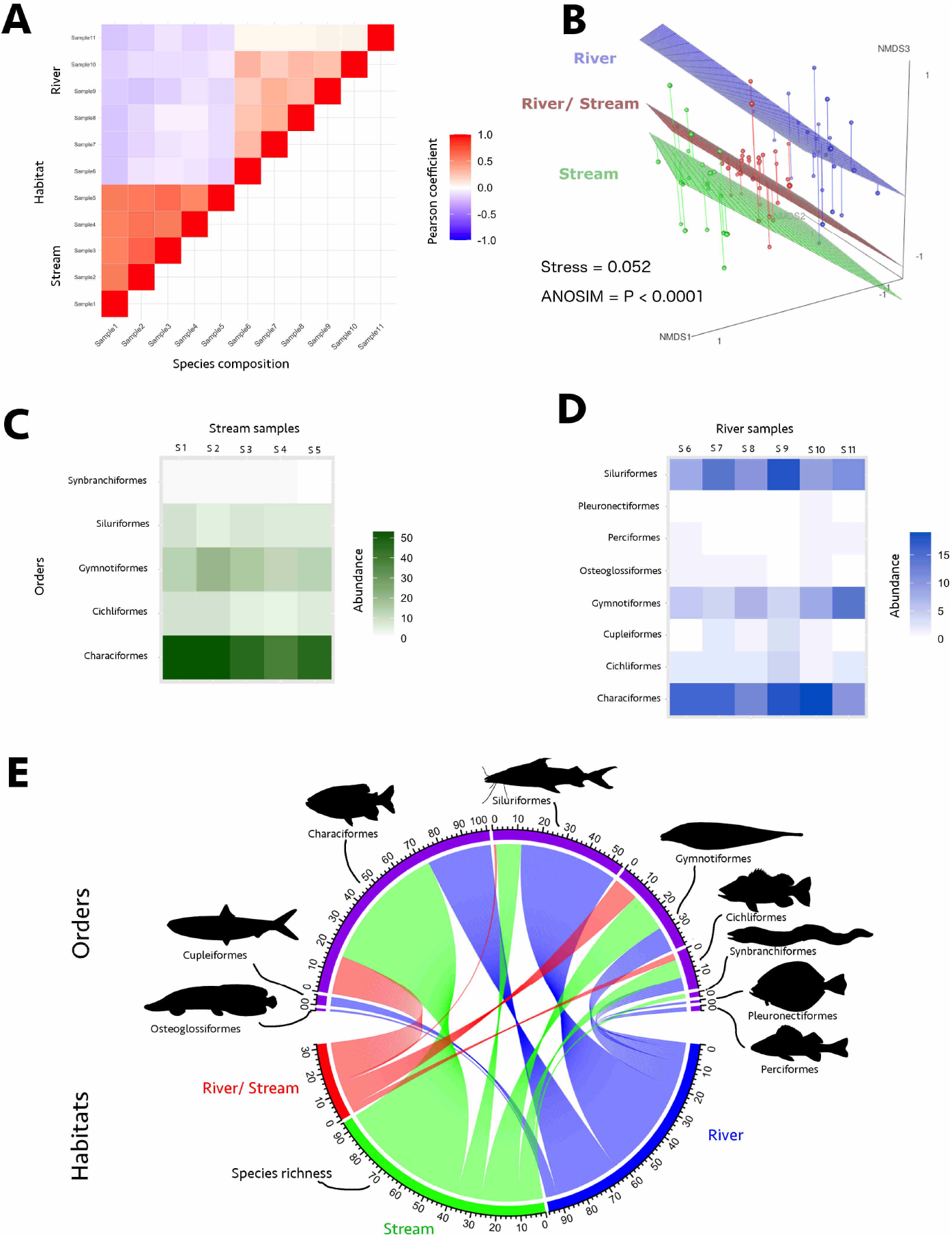
Ichthyofauna segregation into river and stream at Javari Basin as detected by 11 samples of eDNA in the three stations: stream (station 2): samples 1 to 6; Javari river (station 1): Samples 6 to 10; and Quixito River (station 3): sample 11. (A) Heatmap based on the Pearson correlation coefficients between species composition and habitat. Note the difference in the species composition along the river, i.e., Javari versus Quixito rivers (B) Non-metric multidimensional scaling (NMDS) based on Jaccard’s dissimilarities coefficients discriminating habitat (streams vs. rivers). The Stress of the NMDS plot 0.052 indicates that its first three axes provided an appropriate three dimensional representation of the habitats according to their species composition. Each dot represents a species and the relative distance between two points represents the dissimilarity. The ANOSIM p < 0.0001 suggests that NMDS significantly distinguished between the river and stream communities; (C) Heatmap for species abundance for each of the five orders detected in the stream; (D) Heatmap showing species abundance for each of the eight orders detected in the Javari river and Quixito River. Note the alteration in species abundance between samples 5 to 10 (Javari) and 11 (Quixito). In samples 5 to 10 Characiformes and Siluriformes are more abundant. Conversely, Gymnotiformes and Siluriformes are more species rich in the sample 6. (E) Chord diagram showing the directional relationship between habitat and species richness distributed into the nine detected orders.

To assess whether the difference in species composition between stream and river communities observed in the Pearson correlation coefficients were significant, we calculated Jaccard’s dissimilarities indices through a NMDS analysis. The original position of the 222 detected species in river, stream, and in both habitats were represented in a three-dimensional NMDS space (Figure 3B). The Stress = 0.0524 of the NMDS plot indicated that its first three axes provided an appropriate three-dimensional representation of the habitats according to their species composition^36^, and NMDS significantly distinguished between the river and stream communities (ANOSIM R = 0.4327; p < 0.0001; Figure 3B).

Based on the species frequency detected per order we determined the composition of the stream and river habitats (Figures 3B and C). Of note is the difference in the species composition between the five samples from Javari River (Samples 1 to 5) and the single sample (Sample 6) collected in the Quixito River (Figures 3A and D). The interrelationships between habitat and species diversity and composition per order are represented in the chord diagram in Figure 3E.

### 3. Comparing Capture-based sampling (CBS) and Molecular-based sampling (MBS) – eDNA metabarcoding species richness

#### 4.1. Javari River (station 1)

CBS captured a total of 145 species, 101 genera, 32 families and nine orders in the main Javari River. Conversely, MBS found 107 species, 28 genera, 20 families, and seven orders (Figure 4A, B; Tables S5, S6). Thirteen species were detected by both CBS and MBS (Table 1). The rarefaction sampling curve illustrating the accumulation of unique species with the number of individuals collected by CBS does not reach an asymptote (Figure 4C), indicating that several species remain to be detected. This is also corroborated by the Chao II species richness bias-corrected estimator for MBS, which predicated 216 species (95% confidence interval: 163–318).

**Figure 4.**
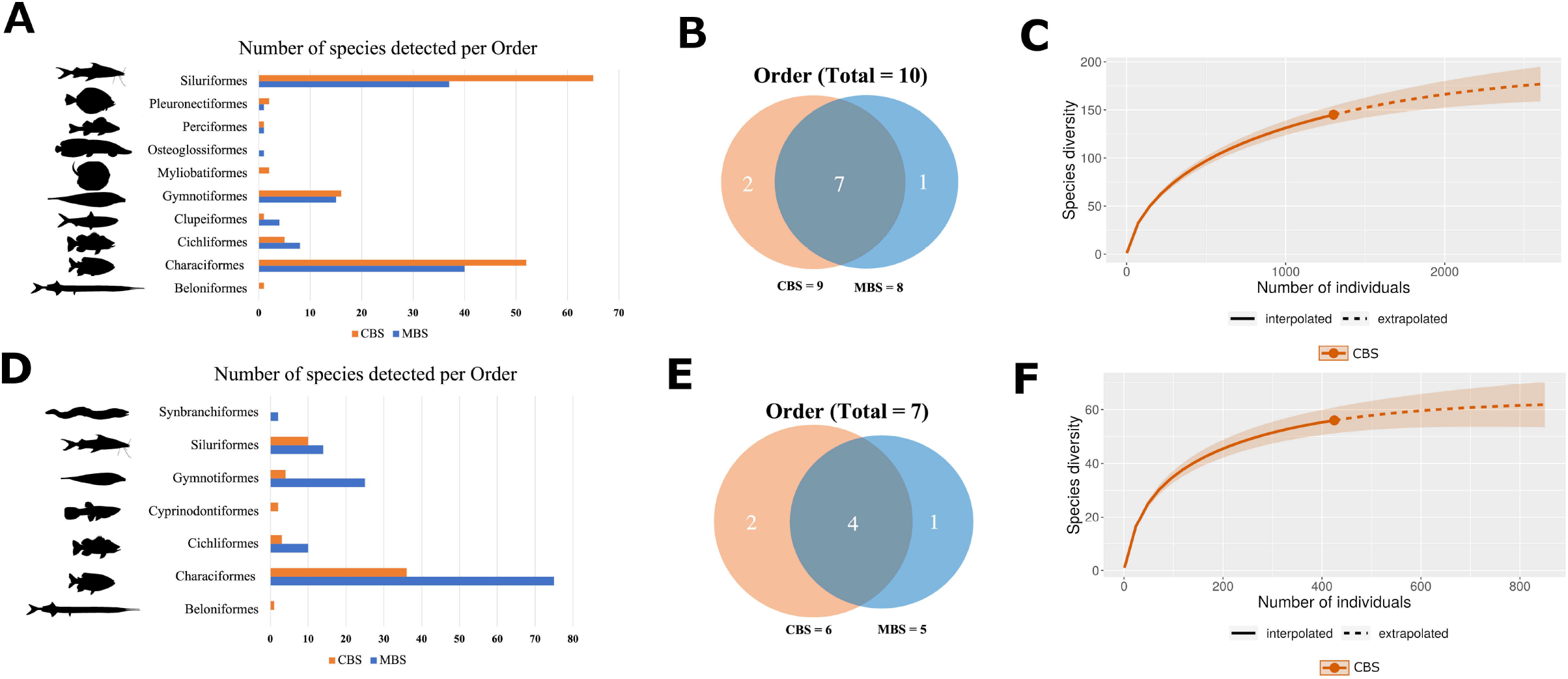
Comparison between Capture-Based Sampling (CBS) in blue and Molecular-Based Sampling (MBS) in red in two sampled localities. Javari River: (A) Histogram comparing the number of species per order detected by CBS and MBS; (B) Venn diagram of the number of orders detected by CBS and MBS; (C) Rarefaction species accumulation curve for CBS with 95% confidence interval and extrapolation for twice the number of individuals sampled. Terra firme stream: (D) Histogram comparing the number of species per order detected by CBS and MBS; (E) Venn diagram of the number of orders detected by CBS and MBS; (F) Rarefaction species accumulation curve for CBS with 95% confidence interval and extrapolation for twice the number of individuals sampled.

#### 4.2. Stream (station 2)

CBS caught 56 species, 38 genera, 21 families, and 6 orders. In contrast, in the stream, MBS detected 126 species, 22 genera, 17 families and 4 orders (Figure 4D, E; Tables S7, S8). Six species were detected by both methodologies (Table 1). The rarefaction curve for CBS extrapolates to slightly over 60 species the diversity in the stream (Figure 4F).

Conversely, MBS Chao II bias- corrected estimator calculated 145 species in the stream (95% confidence interval: 134–172).

## Discussion

### 1. Can eDNA provide an accurate snapshot of the Amazonian megadiverse freshwater ichthyofauna considering current public reference libraries for 12s rRNA?

The Javari River basin contains a considerable fraction of Amazonian fish diversity, ca. 15% of species, 37% of genera, 60% of families, and 83%. It is, therefore, an excellent testing ground for eDNA metabarcoding effectiveness for the Amazonian fish fauna. Based on the current public reference libraries, i.e., Genbank and MiFish DB, MBS provided an accurate snapshot of the Amazonian megadiverse freshwater ichthyofauna at the Javari River basin when we consider higher taxonomic levels, i.e., order.

The detection of 222 species in 11 samples from three stations confirms that eDNA is highly sensitive. However, the low number (28%) of matching sequences with identity of >98.5% in the public reference libraries suggests severe gaps in the library for Amazonian fishes. It corroborates a recent global gap analysis of reference databases^24^, which revealed that 13% of the over 33,000 known teleostean fish species are sequenced for 12S, representing 38% of genera, 80% of families and 98.5% of orders. For freshwater fishes, among all continents, South America and Africa had by far the lowest coverage. Not surprisingly, we found the lowest eDNA identification match at the species level.

Conversely, studies that built reference libraries for highly diverse fish communities considerably improved the match ratio to species identification between capture-based and eDNA approaches. ^30^, for example, identified 65% of 203 species of Guianese fishes. Likewise, ^37^ detected and correctly assigned all 67 species with 12S previously designed primers and reference library in the São Francisco River, Brazil. In contrast, ^32^ assigned only 4 of 84 MOTUs to species, demonstrating problems on the taxonomic resolution in the target gene and general threshold used for species assignment.

The DNA barcoding and eDNA metabarcoding both rely on short, variable, standardized DNA regions, which can be amplified by PCR, sequenced, and analyzed to identify taxa. The eDNA approach for vertebrates does not efficiently employ the COI gene because interspecific genetic variation prevents the use of universal primers^38^ and can result in non-specific amplifications (^39^; but see, ^40^). Instead, rRNA genes used in DNA metabarcoding, such as 12S rRNA (e.g.,^3^), have the acceptable resolution at the species level and an elevated copy number per cell due to the number of mitochondria per cell. Similarly, rRNA genes are preferable over single-copy nuclear DNA, which is less likely to be detected in the environment. Yet, the low substitution rate of rRNA genes will compromise the identification of rapidly evolved and complex fish assemblages such as those in the Neotropical region. Thus, it is likely that, in the near future, DNA barcode and eDNA metabarcode methods will converge to use large portions of the mitochondrial genome. Regardless of the fragment or the threshold used to delimit species (e.g.,^32^), it is essential that studies involving eDNA for assessing fish diversity move towards building robust mitochondrial DNA reference libraries based on vouchered specimens.

### 2. eDNA species detection across heterogeneous aquatic environments

Amazonian aquatic environments are characterized by specialized fish communities segregated across a variety of habitats, such as streams, rivers, and their microhabitats^14,41^. In streams, diverse microhabitats are home to leaf-dwelling, sand-dwelling, and pool-dwelling fish communities^42,43,44^. Similarly, rivers have specialized fish groups living in high-energy or deep water (>5 meters) environments. It is critical that fishes inhabiting all aquatic environments are sampled in biodiversity inventories. Perhaps unsurprisingly, it is incredibly difficult to sample and therefore assess some microhabitats by CBS. For example, some species are buried deep in the roots of plants in the riparian zone (e.g.,^45^), leaf litter, or in the sand of streams that are extremely difficult to collect with traditional sampling gear. These life history strategies naturally underestimate the number of fish species living in these areas due to microhabitat partitioning and undescribed diversity. We corroborate the potential effectiveness of MBS to detect specialized fish assemblages across heterogeneous aquatic environments. More intensive sampling efforts might be required to detect low-occurrence taxa as well as to appropriately sample microhabitats, e.g., filtering a higher amount of water or collecting water from temporary pools and the river bottom.

#### 2.1. River

In the Javari River (station 1), seven orders were detected by CBS and MBS (Characiformes, Cichliformes, Clupeiformes, Gymnotiformes, Perciformes, Pleuronectiformes, Siluriformes). In addition, CBS detected Beloniformes and Myliobatiformes; and MBS found Osteoglossiformes. The absence of Beloniformes and Myliobatiformes in the MBS could be due to the poor reference library for comparisons. In contrast, Osteoglossiformes (*Arapaima gigas*) is well known to occur in the region and specimens were found in the local market. Thus, the absence of *Arapaima* in the CBS was circumstantial.

The difference in species composition between the two methodologies that was detected possibly is due to sampling bias in MBS. Water samples for MBS were only collected at the river surface, detecting mostly free DNA of fish assemblages occurring at midwater and near the surface, where species-diverse Characiformes are the dominant assemblage. Despite that, as aforementioned, MBS was also able to take a snapshot of the benthic fish fauna by detecting many catfish species typically restricted to river channels (e.g.,^46,47^). For example, MBS detected river-dwelling fishes living near the surface, as well as some deep-water (> 5m depth) inhabitants e.g., *Brachyplatystoma* spp. – goliath catfishes; Pleuronectiformes – flatfishes; and a large number of unidentified species of electric fishes (sequences identities within the range of 80–98.5%) belonging to the families Apteronotidae (10 species) and Sternopygidae (15 species) – common, but often underestimated components of rivers (e.g.,^48,49,50^).

In addition, the sole sample collected in the mouth of the Quixito River (station 3) was substantially different from the five samples collected in the Javari River reflecting the different milieu where the samples were collected. The Javari samples were dominated by Characiformes whereas Gymnotiformes dominated in the Quixito River sample. In the Javari River, samples were collected in fast-flowing water along the edge between a shallow peat bog and the main channel. The Quixito River sample was collected at the mouth of the river, characterized by small slow-flowing channel.

#### 2.2. Stream

Typically, Characiformes, Siluriformes, Gymnotiformes, Cichliformes, Cyprinodontiformes, Beloniformes, and Synbranchiformes are the dominant orders in Amazonian streams (e.g.,^51^). At station 2, both approaches detected species belonging to Characiformes, Siluriformes, Gymnotiformes, and Cichliformes. In addition, CBS found Beloniformes and Cyprinodontiformes whereas MBS detected Synbranchiformes for a total of seven orders. The absence of Beloniformes in the MBS may be due to the poor reference library for comparisons, and the absence of Synbranchiformes in the CBS here could be due to the difficulty in collecting cryptobiotic species. We were able to detect at a fine-scale specialized species assemblage restricted to microhabitats. For example, we captured members of the leaf-dwelling (e.g., *Apistogramma* spp. – dwarf cichlids) and sand-dwelling (e.g., *Gymnorhamphichthys* spp. – sand knifefishes) fish communities. It remains to be determined whether eDNA failed to detect fishes that are residents in the temporary pools (e.g., killifishes – Rivulidae) because of the limitation of its radius of action, or due to the poor reference library for Neotropical fishes.

Species diversity in Amazonian Terra firme streams ranges from ca. 30 to 170 species^52^ with Characiformes and Siluriformes being the most species-rich orders (e.g.,^53^). Quantification of fish richness in these streams depends upon the sampling methodology employed and its substrate composition (for reviews see^54,55,56^). For example, in litter banks-rich streams, Gymnotiformes species diversity can surpass Siluriformes (e.g.,^57^). In station 2, according to CBS, Characiformes and Siluriformes were the dominant orders. In contrast, Characiformes followed by Gymnotiformes were the more species-diverse groups. The extremely high number of species detected by MBS in the sampled stream, more than twice that of CBS, primarily in the two dominant orders, Characiformes and Gymnotiformes, is likely related to four different issues. First, MBS was collected near the confluence between the river and stream, which may have resulted in occasional, wandering river fishes. Second, the CBS was conducted with a standardized sampling effort in a restrict (50-m) stretch of the stream (e.g.,^55^), not including its headwaters and areas near its mouth. Third, Characiformes undoubtedly contain hidden species diversity. This is corroborated by the historical difficulty in identification of small tetra species, wherein one named species may represent several undescribed species, such as in *Astyanax* (e.g.,^58,59^). Fourth, diversity is also underestimated for the Gymnotiformes (e.g.,^60^), for which difficulties in capturing species with cryptobiotic habits possibly play a critical role in the underestimation of their diversity by CBS methods (sub-estimative may reach three times the local species richness and up to 10 times the specimens abundance; JZ, unpublished data).

#### 3. The role of natural history museums in the advance of eDNA studies

The biodiversity crisis is one of the grand challenges of the 21st century^61,62^ with the next two decades critical for the conservation of freshwater environments. Freshwater ecosystems worldwide hold ca. 30% of vertebrate diversity, including ca. 50% of all fish species diversity, and are one of the most vulnerable environments on Earth^35,62,63,64,65^.

Combining specimens, DNA sampling and taxonomic identification is required to obtain a comprehensive assessment of biodiversity. Yet, DNA samples are available for less than 10% of the specimens deposited in most fish repositories. Since most fish specimens deposited in museums and other repositories were collected before the development of PCR, a vast majority were fixed in formalin, a standard method of fixation for over a century. Despite the advances in the techniques of DNA extraction from formalin-fixed materials, the success of these techniques is still limited, especially for specimens stored for long periods in unbuffered solutions^66,67,68^. Thus, well-identified vouchered DNA tissue samples are critical for the identification of unknown DNA in environmental samples. These DNA tissues may be stored as dried, frozen, or alcohol-fixed samples or as cryopreserved living samples that have broad potential applications (e.g.,^69^). However, scientific collections in regions holding most of the fish diversity, such as the Neotropics, often lack the ideal infrastructure to hold long-term genetic resources (e.g. ultrafreezers, liquid nitrogen storage, cryo-facilities). Nevertheless, GGBN has targeted and sometimes funded Neotropical institutions to build biorepository capacity and to make their collections globally discoverable.

These limitations are particularly worrisome given the stark reality of anthropogenic destruction, climate change and the great extent of predicted unknown diversity that remains to be described in the Amazon rainforest^70,71^. These factors make this area and Earth’s other hotspots of biodiversity priority targets for complete species inventories in the next decade before suffering irreversible damage (e.g.,^72^). Another advantage of eDNA is the long-term biodiversity monitoring in preserved areas/conservation units (e.g.,^73^). The use of eDNA is a highly valuable and cost-effective way to monitor biodiversity, especially in areas with low anthropogenic threats^74^. This would allow a better prioritization of scarce resources for research and/or conservation actions.

In the face of these challenges, natural history museums should play a primary role in the development of eDNA as a tool of biodiversity inventories as well as to track changes in biodiversity hotspots by: (1) prioritizing expeditions to jointly secure DNA samples, vouchers, and eDNA in Earth’s hotspots of biodiversity; (2) adapting their biorepositories to archive eDNA samples, which as a consequence, would provide samples not only for analysis with current but heretofore unseen technologies; (3) creating reference libraries for the mitochondrial genome; (4) backing up DNA samples with species-level accuracy on the identification of vouchered specimens; (5) expanding and improving their tissue biobanks. It is crucial that these modifications for eDNA storage also occur in museums in Neotropical and Afrotropical countries, which host most of freshwater fish diversity yet lack the resources to build and maintain these tissue collections in perpetuity^75,76,77^. These efforts would maximize the information extracted from eDNA metabarcoding and DNA samples, facilitate the design of sets of universal primers for broader biodiversity inventories, monitor hotspots of biodiversity, and support taxon-specific surveys; (6) improving public platforms to close gaps in sampling information and making possible access to DNA sequences; (7) training students and researchers to use CBS, MBS, morphology and molecular-based taxonomy to survey and identify biodiversity. By combining eDNA with tissues associated with museum-curated voucher specimens we can continue to fill gaps currently missing in our knowledge of biodiversity. The high frequency of our lowest taxonomic identifications ending with “sp.,” species undetermined, when assessing species diversity using a new technology highlights the need for highly trained taxonomic specialists. Finally, (8) using eDNA research as a gateway to inspire and engage society in natural history and the race against time to survey and protect Earth’s hotspots of biodiversity through education and citizen science programs. Considering the simplicity of implementing MBS in certain aquatic environments, such as rivers (see Methods), scientific communities at natural history museums can launch regional/ global outreach and human resource training initiatives involving citizen scientists, K-12 students, and professional scientists. Likewise, it would create niches for large-scale natural history museums to work with regional-scale scientific institutions worldwide, such as in the training of human resources (e.g., technicians to curate genetic resources) and promoting horizontal transfer of technology in South America and Africa (e.g., eDNA methodology). In sum, activities involving eDNA have the potential to fulfill the priorities of natural history museums in the 21st century: research, collections, training, and outreach.

One successful initiative is the DNA barcoding and metabarcoding libraries for Amazonian fishes supported by Smithsonian’s Global Genome Initiative (GGI), DNA Barcode Alliance, and São Paulo Research Foundation (FAPESP). The current project is the first of many scientific expeditions planned over the next three years to survey fishes in poorly explored areas of the Amazon basin supported by these three initiatives. DNA and eDNA samples and vouchers are being used to develop a robust, well-documented, mitochondrial DNA reference database. This eDNA database is validated by morphological (phenotypic) vouchers. Additional eDNA samples have been collected and deposited in the Smithsonian Institution’s National Museum of Natural History Biorepository. We aim to make available an online platform of DNA sequences of all orders and families, most of the genera, and a significant number of species of Amazonian fishes. Likewise, GGI is also supporting an initiative for African freshwater fishes. These actions together with the ongoing development of eDNA technology and bioinformatics will enable the use of eDNA metabarcoding in fish inventories and the more effective monitoring of hotspots of biodiversity worldwide.

## Acknowledgements

The expedition to the Javari River was funded by São Paulo State Research Foundation grant to GTV (FAPESP #2016/07910-0) and Global Genome Initiative grant to CDS, CBD, and LRP (GGI-Peer-2017-149). A Global Genome Initiative grant to CDS to survey African rivers and build DNA barcoding and metabarcoding archives for freshwater fishes (GGI-Peer-2020-258). JZ received a productivity grant from Brazil’s CNPq (#313183/2014-7). AD, CDS, and NAM are funded by FAPESP (#2016/19075-9). MM and TS were supported by Environment Research and Technology Development Fund (4-1602) of the Ministry of the Environment, Japan.

## Author Contribution

The study was conceived by CDS and designed by CDS and MM. Environmental samples were collected by CDS, DAB, JZ, GTV. MM and TS performed laboratory analyses. Bioinformatic and statistical analyses were performed by CDS and MM. The manuscript was written by CDS. with input from all the authors. CDS coordinated the study.

## Data Availability

Data will be made public on the DRYAD repository upon acceptance.

## Methods

### Study area

The Javari River encompass an area of 109.202 km^2^ with a 1180 km of a main white water river channel (*sensu* Sioli, 1967; i.e., pH-neutral low-transparency, alluvial sediment-laden tributary of the Amazon River forming the border between Brazil, Peru and Colombia for ca. 800 km. The first formal records for the Javari river basin were obtained during the Thayer Expedition to Brazil, in 1865. Most of region remained largely unexplored until our survey conducted along the Javari River basin during the low water season in July-August of 2017.

### Specimens sampling and identification

All samples were collected under Jansen Zuanon permanent permit (SISBIO # 10199-3). Capture-based specimens were sampled at 46 localities along the Javari River basin (Figure 1) during the low water season in July-August, 2017, using gill nets, cast nets, hand nets, and trawl nets in rivers, rapids, beaches, streams, and lakes (Table S1). All fish specimens collected were identified to species level and deposited at the Instituto Nacional de Pesquisas da Amazônia (INPA) under the numbers INPA-ICT 055148 to INPA-ICT 057159, in Brazil.

### Water sampling sites and on-site filtration

Eleven water samples were collected from water surface at three stations to represent the Javari fish fauna: Station 1, Figure 1; JAV2017081606 (5 samples) – Javari River, below Limoeiro (−4.176, −70.779); Station 2, Figure 1; JAV2017082108 (5 samples) – “Terra firme” clearwater stream (locally named as “igarapés”), i.e., acid, highly-transparent and shallow (depth <2 meters), Palmari community (−4.293, −70.291); and Station 3, QUI 2017082906 (1 Sample) – Quixito River (−4.428, −70.260). We used low-tech bucket-sampling to collect freshwater using a 10L polypropylene bucket fastened to a 5 m rope (nylon rope, 6 mm in diameter) to collect 5L of water. Before the water sampling, we wore disposable gloves on both hands and assembled two sets of on-site filtration kits consisting of a Sterivex filter cartridge (pore size 0.45 μm; Merck Millipore, MA, USA) and a 50 mL disposable syringe. Then we thoroughly decontaminated the bucket with a foam-style 10% bleach solution and brought the equipment to the sampling point. We fastened one end of the 5 m rope to the bucket and collected surface freshwater by tossing and retrieving it. We repeated collection of fresh water three times to minimize sampling biases at each station.

We performed on-site filtration using two pairs of the filtration kits described above (filter cartridge + syringe) to obtain duplicate samples. With each collection of fresh water, we removed the filter cartridge from the syringe, drew approximately 50 ml freshwater into the syringe by pulling the plunger, reattached the filter cartridge to the syringe, and pushed the plunger to filter the water. We repeated this step twice in each toss of the bucket sampling so that the final filtration volume reached 100 ml × 2 with three tosses of the bucket. When the filter was clogged before reaching 100-ml filtration, we recorded the total volume of water filtered (70–100 mL from three stations).

After on-site filtration, we sealed an outlet port of the filter cartridge with Parafilm (Bemis NA, Wisconsin, USA), added 2 ml of RNAlater (Thermo Fisher Scientific, DE, USA) into the cartridge from an inlet port of the cartridge using a disposable capillary pipette (Kinglate, USA) to prevent eDNA degradation, and then sealed the inlet port either with Parafilm or a cap for preservation. Filtered cartridges filled with RNAlater were kept in –20°C freezers until shipment to MM’s lab at Natural History Museum and Institute, Chiba, Japan. Samples shipped under export for biological material permit at room temperature using an overseas courier service.

### DNA extraction

All DNA experiments were conducted in MM’s lab. We sterilized the workspace and all equipment before DNA extraction. We used filtered pipette tips and conducted all eDNA-extractions and manipulations in a dedicated room that is physically separated from pre- and post-PCR rooms to safeguard against cross-contamination from PCR products.

We extracted eDNA from the filter cartridges using a DNeasy Blood & Tissue kit (Qiagen, Hilden, Germany) following the methods developed and visualized by ^78^ with slight modifications.

We connected an inlet port of each filter cartridge with a 2.0-ml collection tube and tightly sealed the connection between the cartridge and collection tube with Parafilm. We inserted the combined unit into a 15-ml conical tube and centrifuged the capped conical tube at 6,000xg for 1 min to remove freshwater and RNAlater. After centrifugation we discarded the collection tube and used an aspirator (QIAvac 24 Plus, Qiagen, Hilden, Germany) to completely remove liquid remaining in the cartridge.

We subjected the filter cartridge to lysis using proteinase K. Before the lysis, we mixed PBS (220 μl), proteinase K (20 μl) and buffer AL (200 μl), and gently pipetted the mixed solution into the cartridge from an inlet port of the filter cartridge. We again sealed the inlet port and then placed the cartridge in a 56°C preheated incubator for 20 min while stirring the cartridge using a rotator (Mini Rotator ACR-100, AS ONE, Tokyo, Japan) with a rate of 10 rpm. After the incubation, we removed the film from the inlet port and connected the port with a 2-ml tube (DNA LowBind tube, SARSTEDT, Tokyo, Japan) for DNA collection. We placed the combined unit in a 50-ml conical tube and centrifuged the capped tube at 6,000xg for 1 min to collect the DNA extract.

We purified the collected DNA extract (*ca*. 900 μl) using the DNeasy Blood and Tissue kit following the manufacture’s protocol with a final elution volume of 200 μl. We completed DNA extraction in one round and used one more premix for the extraction blank (EB) to monitor contamination. All DNA extracts were frozen at −20°C until paired-end library preparation.

### Paired-end library preparation and sequencing

We sterilized the workspace and equipment in the pre-PCR area before library preparation. We used filtered pipette tips and performed pre- and post-PCR manipulations in two different, dedicated rooms to safeguard against cross contamination.

We employed a two-step PCR for paired-end library preparation on the MiSeq platform (Illumina, CA, USA) and generally followed the methods developed by ^3^. For the first-round PCR (1st PCR), we used a mixture of the following four primers: MiFish-U-forward (5’–ACA CTC TTT CCC TAC ACG ACG CTC TTC CGA TCT NNN NNN GTC GGT AAA ACT CGT GCC AGC–3’), MiFish-U-reverse (5’–GTG ACT GGA GTT CAG ACG TGT GCT CTT CCG ATC TNN NNN NCA TAG TGG GGT ATC TAA TCC CAG TTT G–3’), MiFish-E-forward-v2 (5’–ACA CTC TTT CCC TAC ACG ACG CTC TTC CGA TCT NNN NNN RGT TGG TAA ATC TCG TGC CAG C–3’) and MiFish-E-reverse-v2 (5’–GTG ACT GGA GTT CAG ACG TGT GCT CTT CCG ATC TNN NNN NGC ATA GTG GGG TAT CTA ATC CTA GTT TG– 3’). These primer pairs amplify a hypervariable region of the mitochondrial 12S rRNA gene (*ca*. 172 bp; hereafter called “MiFish sequence”) and append primer-binding sites (5’ ends of the sequences before six Ns) for sequencing at both ends of the amplicon. We used the six random bases (Ns) in the middle of those primer to enhance cluster separation on the flow cells during initial base call calibrations on the MiSeq platform.

We carried out the 1st PCR with 35 cycles in a 12-μl reaction volume containing 6.0-μl 2 × KAPA HiFi HotStart ReadyMix (KAPA Biosystems, MA, USA), 2.8 μl of a mixture of the four MiFish primers in an equal volume (U/E forward and reverse primers; 5 μM), 1.2-μl sterile distilled H_2_O and 2.0-μl eDNA template (a mixture of the duplicated eDNA extracts in an equal volume). To minimize PCR dropouts during the 1st PCR, we performed 8 replications for the same eDNA template using a strip of 8 tubes (0.2 μl). The thermal cycle profile after an initial 3 min denaturation at 95°C was as follows: denaturation at 98°C for 20 sec, annealing at 65°C for 15 sec and extension at 72°C for 15 sec with the final extension at the same temperature for 5 min. We also made a 1st PCR blank (1B) during this process in addition to EB. Note that we did not perform 8 replications and used a single tube for each of the two blanks (EB, 1B) to minimize cost of the experiments.

After completion of the 1st PCR, we pooled an equal volume of the PCR products from the 8 replications in a single 1.5-ml tube and purified the pooled products using a GeneRead Size Selection kit (Qiagen, Hilden, Germany) following the manufacturer’s protocol for the GeneRead DNA Library Prep I Kit. This protocol repeats the column purification twice to completely remove adapter dimers and monomers. Subsequently we quantified the purified target products (ca. 172 bp) using TapeStation 2200 (Agilent Technologies, Tokyo, Japan), diluted it to 0.1 ng/μl using Milli Q water and used the diluted products as templates for the second-round PCR (2nd PCR). For the two blanks (EB, 1B), we purified the 1st PCR products in the same manner, but did not quantify the purified PCR products, diluted them with an average dilution ratio for the positive samples, and used the diluted products as templates for the 2nd PCR.

For the 2nd PCR, we used the following two primers to append dual-index sequences (8 nucleotides indicated by Xs) and flowcell-binding sites for the MiSeq platform (5’ ends of the sequences before eight Xs): 2nd-PCR-forward (5’–AAT GAT ACG GCG ACC ACC GAG ATC TAC ACX XXX XXX XAC ACT CTT TCC CTA CAC GAC GCT CTT CCG ATC T–3’); and 2nd-PCR-reverse (5’–CAA GCA GAA GAC GGC ATA CGA GAT XXX XXX XXG TGA CTG GAG TTC AGA CGT GTG CTC TTC CGA TCT–3’).

We carried out the 2nd PCR with 10 cycles of a 15-μl reaction volume containing 7.5-μl 2 × KAPA HiFi HotStart ReadyMix, 0.9-μl each primer (5 μM), 3.9-μl sterile distilled H_2_O and 1.9-μl template (0.1 ng/μl with the exceptions of the three blanks). The thermal cycle profile after an initial 3 min denaturation at 95°C was as follows: denaturation at 98°C for 20 sec, annealing and extension combined at 72°C (shuttle PCR) for 15 sec with the final extension at the same temperature for 5 min. We also made a 2nd PCR blank (2B) during this process in addition to EB and 1B.

To monitor for contamination during the DNA extraction, 1st and 2nd PCRs of the 11 samples, we made a total of 3 blanks (EB, 1B, 2B) and subjected them to the above library preparation procedure.

We pooled each individual library in an equal volume into a 1.5-ml tube. Then we electrophoresed the pooled dual-indexed libraries using a 2% E-Gel Size Select agarose gel (Invitrogen, CA, USA) and excised the target amplicons (*ca*. 370 bp) by retrieving them from the recovery wells using a micropipette. The concentration of the size-selected libraries was measured using a Qubit dsDNA HS assay kit and a Qubit fluorometer (Life Technologies, CA, USA), diluted them at 12.0 pM with HT1 buffer (Illumina, CA, USA) and sequenced on the MiSeq platform using a MiSeq v2 Reagent Kit for 2 × 150 bp PE (Illumina, CA, USA) following the manufacturer’s protocol. We subjected the pooled dual-indexed libraries a MiSeq run with a PhiX Control library (v3) spike-in (expected at 5%).

### Data preprocessing and taxonomic assignment

We performed data preprocessing and analysis of MiSeq raw reads using USEARCH v10.0.240^79^ according to the following steps: 1) Forward (R1) and reverse (R2) reads were merged by aligning the two reads using the *fastq_mergepairs* command. During this process, low-quality tail reads with a cut-off threshold set at a quality (Phred) score of 2, too short reads (<100 bp) after tail trimming and those paired reads with too many differences (>5 positions) in the aligned region (*ca*. 65 bp) were discarded; 2) primer sequences were removed from those merged reads using the *fastx_truncate* command; 3) those reads without the primer sequences underwent quality filtering using the *fastq_filter* command to remove low quality reads with an expected error rate of >1% and too short reads of <120 bp; 4) the preprocessed reads were dereplicated using the *fastx_uniques* command and all singletons, doubletons, and tripletons were removed from the subsequent analysis following the recommendation by the author of the program^79^; 5) the dereplicated reads were denoised using the *unoise3* command to generate amplicon sequence variants (ASVs) that remove all putatively chimeric and erroneous sequences^80^; 6) finally ASVs were subjected to taxonomic assignments to species names (metabarcoding operational taxonomic units; MOTUs) using the *usearch_global* command with a sequence identity of >98.5% with the reference sequences and a query coverage of ≥90% (two nucleotide differences allowed).

Those ASVs with the sequence identities of 80–98.5% were tentatively assigned “U98.5” labels before the corresponding species name with the highest identities (*e.g., U98.5_Synbranchus marmoratus*) and they were subjected to clustering at the level of 0.985 using *cluster smallmem* command. An incomplete reference database necessitates this clustering step that enables detection of multiple MOTUs under an identical species name. We annotated such multiple MOTUs with “gotu1, 2, 3…” and tabulated all the outputs (MOTUs plus U98.5_MOTUs) with read abundances. We excluded those ASVs with sequence identities of <80% (saved as “no_hit”) from the above taxonomic assignments and downstream analyses, because all of them were found to be non-fish organisms. For a reference database, we used MiFish DB ver. 36 for taxa assignment, which contained 7,973 species distributed across 464 families and 2,675 genera. In addition, we downloaded all the fish whole mitochondrial genome and 12S rRNA gene sequences from Genbank as of 15 December 2020.

We refined the above automatic taxonomic assignments with reference to a family-level phylogeny based on MiFish sequences from both MOTUs and the reference database. For each family, we assembled representative sequences (most abundant reads) from MOTUs (including U98.5) and added all reference sequences from that family and an outgroup (a sequence from a closely-related family) in FASTA format. We subjected the FASTA file to multiple alignment using MAFFT^81^ with a default set of parameters. We constructed a neighbor-joining (NJ) tree with the aligned sequences in MEGA7^82^ using pairwise deletion of gaps and the Kimura two-parameter distances^83^ with the among-site rate variations modeled with gamma distributions (shape parameter = 1). We assessed statistical support for internal branches of the NJ tree using the bootstrap resampling technique (100 resamplings). In addition, aligned sequences were submitted to Bayesian Inference (BI) analyses run for 10 million generations sampling every 1000 generations to determine posterior probability for each MOTU and reference sequences. Models were obtained on JModeltest2^84^. BI analyses were run in the Mr. Bayes v3.2.7^85^. Some of the BI analyses were conducted on the CIPRES science gateway v3.3^86^. Trees were analyzed and rendered in iTOL v5.7^87^.

The MiSeq paired-end sequencing (2 × 150 bp) of the 11 libraries, together with an additional 88 libraries (total = 99), yielded a total of 5,274,381 reads, with an average of 96.5% base calls, with Phred quality scores of ≥30.0 (Q30; error rate = 0.1% or base call accuracy = 99.9%). This run was highly successful considering the manufacture’s guidelines (Illumina Publication no. 770-2011-001 as of 27 May 2014) are >80% bases ≥Q30 at 2 × 150 bp.

Of the 5,274,381 reads, a total of 1,903,160 reads were assigned to the 11 libraries, and the number of raw reads for each library ranged from 135,818 to 213,952 with an average of 173,015 reads (Table S8). After merging the two overlapping paired-end FASTq files (1,826,828 reads [96.0%]), the primer-trimmed sequences were subjected to quality filtering to remove low-quality reads (1,802,098 reads [94.7%]). The remaining reads were dereplicated for subsequent analysis, and single- to tripletons were removed from the unique sequences as recommended by the author of the program^79^. Then, reads were denoised to remove putatively erroneous and chimeric sequences, and the remaining 1,677,402 reads (88.1% of the raw reads) were subjected to taxon assignments. Of these, 1,671,871 reads (99.7% of the denoised reads) were putatively considered as sequences for fishes, and BLAST searches indicated that non-fish sequences (5,531 reads [0.3%]) mostly consisted of mammals (i.e., cows, pigs, and humans) and a few unknown sequences. The three negative controls (i.e., EB, 1B, and 2B) were subjected to the same analysis pipeline and yielded only 103 denoised reads in total (only 0.006% of the total raw reads), which were not taken into consideration in the subsequent analyses as their subtraction from the corresponding species did not affect the presence/absence data matrix of sequences assignable to fishes. Contamination from non-Amazonian fishes at Miya’s lab was detected and removed (Table S9).

### Statistical analyses

All statistical analyses were conducted in R v.4.0.2^88^.

### Community structure - Molecular-based sampling (MBS)

Evaluation of species richness for eDNA included all 11 samples from the river and stream localities. Specifically for river: five samples from station 1 (JAV2017081606) and one sample from station 3 (QUI 2017082906); stream: five samples from station 2 (JAV2017082108). Species richness between CBS and MBS was performed by comparing fish assemblages captured and detected in stations 1 and 2 only.

Species abundance per order was evaluated by heatmaps produced in *ggplot2*^89^. Composition per Similarity among all 11 samples, three stations, versus stream and river assemblages were calculated using the Pearson correlation coefficient. Then, we calculated Jaccard’s dissimilarities, and the coefficient values were ordinated using non-metric multidimensional scaling (NMDS) to visualize how replicated eDNA data discriminate sites and habitat (streams vs. rivers) patterns and to determine the sampling effort needed to identify community changes among sites in the VEGAN package version 2.4–4^90^. A 3D graph was produced in CAR^91^ and GLR version 0.103.5^92^ packages. Differences in species compositions between sites and habitat types were statistically tested by permutational analysis of similarities (ANOSIM). It allowed for test of the statistical significance of similarity between groups comparing to the within groups similarity using the rank of similarity values^36^. A chord diagram showing the inter-relationship between species composition and habitat (river versus stream) was produced using the Circlize package^93^. Fish silhouettes were produced in Fishsualize v. 0.2.1^94^ with the addition of a species of Gymnotiformes.

### Species richness

Water samples station 1 and station 2: The number of detected taxa between CBS and MBS were represented by Venn diagrams. Rarefaction species accumulation curve for capture-based sampling were calculated for stations 1 and 2^95^ using iNEXT package in R^96^ for Hill number with order *q* = 0 (species richness) with 1000 bootstraps. The dissimilarity species composition among samples in stations 1 and 2 were assessed by calculating pairwise Jaccard’s distances with the function *vegdist.* Bias-corrected estimators Chao II^97^ was applied to calculate species richness detected by MBS, as suggested by ^98^. It was calculated in SpadeR package in R^99^. Species accumulation curves for molecular-based sampling were built using the function *specaccum* in VEGAN package v2.5.4^90^. Graphs were plotted using *ggplot2.*

---

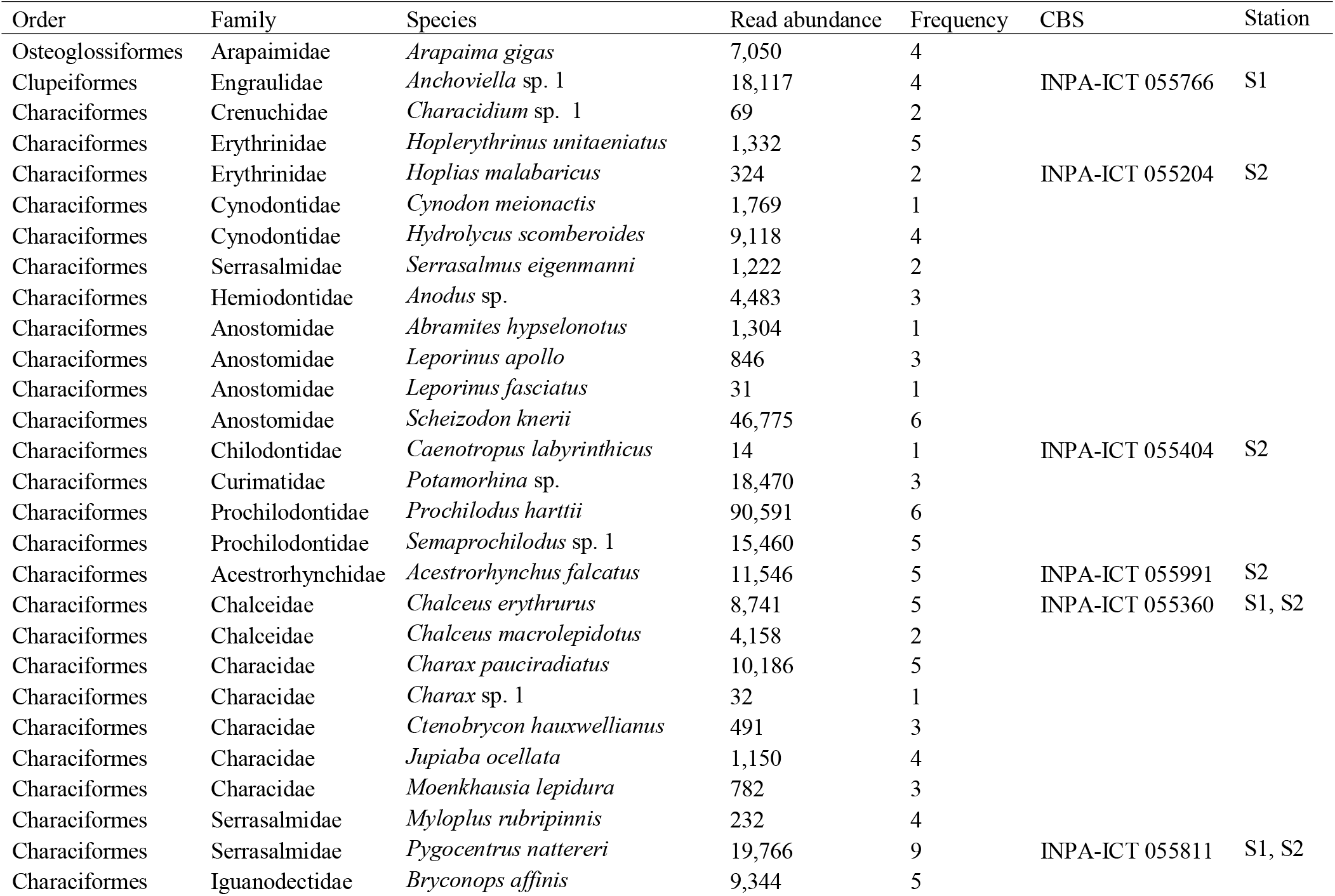
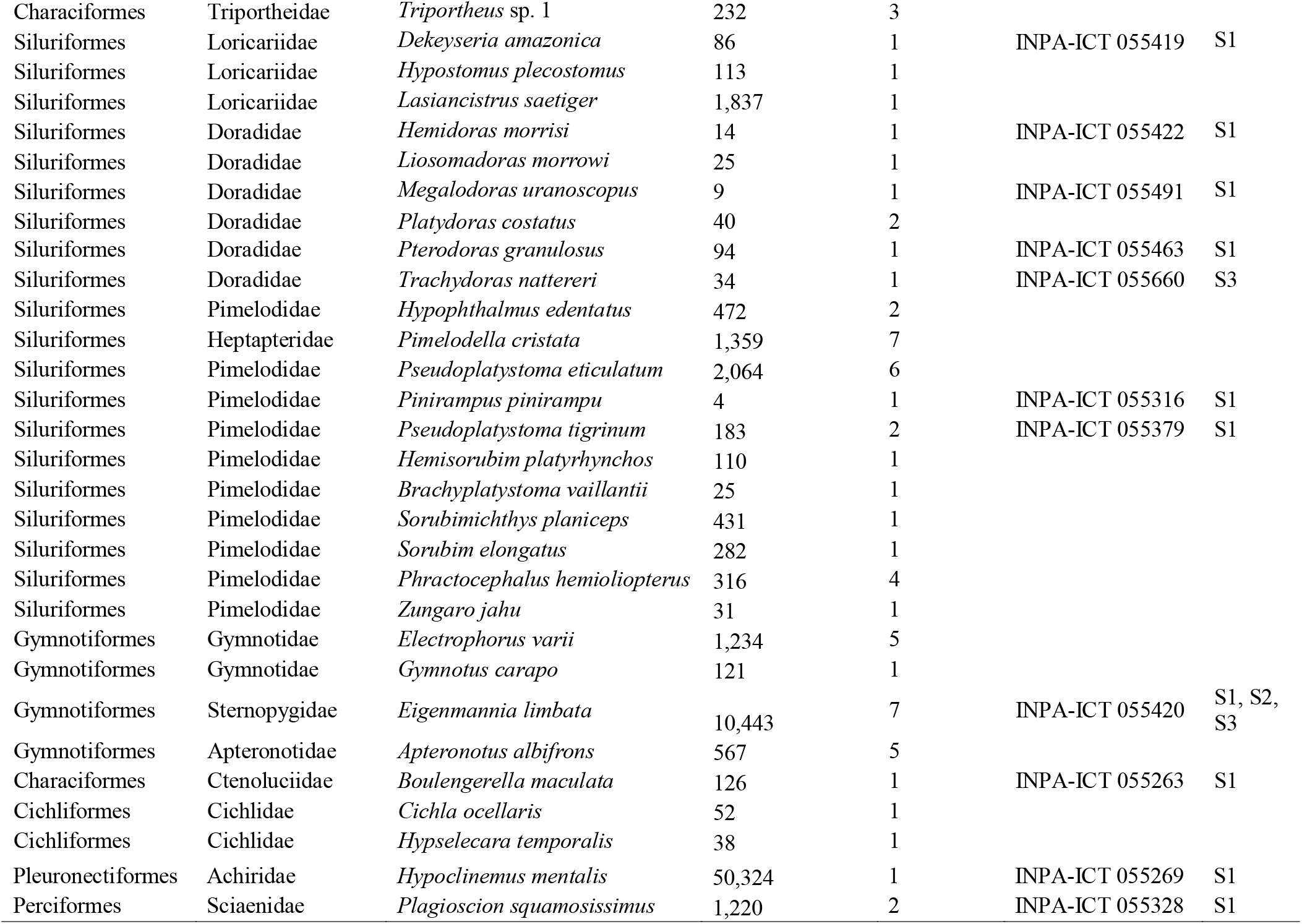

## References

1. Lundberg J. G., M. Kottelat, G. R. Smith, M. L. J. Stiassny, & A. C. Gill. 2000. So Many Fishes, So Little Time: An Overview of Recent Ichthyological Discovery in Continental Waters. Annals of the Missouri Botanical Garden, 87: 26–62.

2. Relyea R. A. 2005. The impact of insecticides and herbicides on the biodiversity and productivity of aquatic communities. Ecological Applications, 15: 618–627.

3. Miya M., Y. Sato, T. Sado, J. Y. Poulsen, K. Sato, T. Minamoto, S. Yamamoto, H. Yamanaka, H. Araki, M. Kondoh, & W. Ywasaki. 2015. MiFish, a set of universal PCR primers for metabarcoding environmental DNA from fishes: detection of more than 230 subtropical marine species. Royal Society Open Science, 2: 150088.

4. Clare A. I. M., M. Knapp, N. J. Gemmell, J. Gert-Jan, M. Bunce, M. D. Lamare, H. R. Taylor, R. Helen. 2019. Beyond Biodiversity: Can Environmental DNA (eDNA) Cut It as a Population Genetics Tool? Genes, 10: 192.

5. Tsuji S., N. Shibata, H. Sawada, & M. Ushio. 2020. Quantitative evaluation of intraspecific genetic diversity in a natural fish population using environmental DNA. Molecular Ecology Resources, 20: 1323–1332.

6. Miya M., R. O. Gotoh, & T. Sado. 2020. MiFish metabarcoding: a high-throughput approach for simultaneous detection of multiple fish species from environmental DNA and other samples. Fisheries Science, 86: 939–970.

7. Dagosta F. C. P. & M. C. C. de Pinna. 2019. The fishes of the Amazon: distribution and biogeographical patterns, with a comprehensive list of species. Bulletin of the American Museum of Natural History, 431.

8. Jézéquel C., P. A. Tedesco, & R. Bigorne. 2020. A database of freshwater fish species of the Amazon Basin. Scientific Data, 7: 96.

9. Reis R. E., S. O. Kullander, & C. J. Ferraris. 2003. Check list of the freshwater fishes of South and Central America. Edipucrs.

10. Tedesco P., et al. 2017. A global database on freshwater fish species occurrence in drainage basins. Scientific Data 4: 170141.

11. Brito P. M., F. J. Meunier, & M. E. C. Leal. 2007. Origine et diversification de líchthyofaune Neotropical: une revue. Cybium, 31: 139–153.

12. Lowe-McConnell R. H. 1987. Ecological studies in tropical fish communities. Cambridge Tropical Biology Series. Cambridge University Press, Cambridge.

13. Bloom D. D. & N. R. Lovejoy. 2017. On the origins of marine derived fishes in South America. Journal of Biogeography, 44: 1927–1938.

14. de Santana C. D., et al. 2019. Unexpected species diversity in electric eels with a description of the strongest living bioelectricity generator. Nature Communications, 10: 4000.

15. Carvalho, L. N., J. Zuanon, & I. Sazima. 2007. Natural history of Amazon fishes. In: K. Del-Claro (Ed.), Tropical Biology and Natural Resources Theme, In: K. Del-Claro & R. J. Marquis (Session Eds. the Natural History Session), Encyclopedia of Life Support Systems (EOLSS). Eolss Publishers, Oxford.

16. Cardoso Y. P., J. J. Rosso, E. Mabragaña, M. Gonzalez-Castro, M. Delpiani, & E. Avigliano. 2018. A continental-wide molecular approach unraveling mtDNA diversity and geographic distribution of the Neotropical genus *Hoplias*. Plos One, 13: e0202024.

17. Hebert P. D. N., A. Cywinska, S. L. Ball, & J. R. de Waard. 2003. Biological identifications through DNA barcodes. Proceedings of the Royal Society London. Series B: Biological Sciences, 270: 313–321.

18. Baldwin, C. C., C. I. Castillo, L. A. Weigt, & B. C. Victor. 2011. Seven new species within western Atlantic *Starksia atlantica, S. lepicoelia*, and *S. sluiteri* (Teleostei, Labrisomidae), with comments on congruence of DNA barcodes and species. ZooKeys, 79: 21–27.

19. Robertson D. R., A. Angulo, C. C. Baldwin, C. Pitassy, E. Diane, A. W. Driskell, L. A. Weigt, A. Lee & I. J. F. Navarro. 2017. Deep-water bony fishes collected by the B/O Miguel Oliver on the shelf edge of Pacific Central America: an annotated, illustrated and DNA-barcoded checklist. Zootaxa, 4348: 1–125.

20. Weigt L. A., C. C. Baldwin, A. Driskell, D.G. Smith, A. Ormos, & E. A. Reyier. 2012. Using DNA Barcoding to Assess Caribbean Reef Fish Biodiversity: Expanding Taxonomic and Geographic Coverage. Plos One, 7: e41059.

21. Seberg O., G. Droege, K. Barker, J. A. Coddington, V. Funk, M. Gostel., G. Petersen, & P. P. Smith. 2016. Global Genome Biodiversity Network: saving a blueprint of the Tree of Life - a botanical perspective. Annals of Botany, 118: 393–399.

22. Parenti L. R., D. Pitassy, Z. Jaafar, K. Vinnikov, N. E. Redmond & K. S. Cole. 2020. Fishes collected during the 2017 MarineGEO assessment of Kāne’ohe Bay, O‘ahu, Hawai‘i. Journal of the Marine Biological Association of the United Kingdom, 100: 607–637.

23. Droege G. et al. 2016. The Global Genome Biodiversity Network (GGBN) Data Standard specification. Database: 10.1093/database/baw125.

24. Marques, V., P. É. Guérin, M. Rocle, A. Valentini, S. Manel, D. Mouillot, & T. Dejean, 2020. Blind assessment of vertebrate taxonomic diversity across spatial scales by clustering environmental DNA metabarcoding sequences. Ecography, 43: 1779–1790.

25. Leray, M., N. Knowlton, Shien-Lei, H., B. N. Nguyen, & R. J. Machida. 2019. GenBank is a reliable resource for 21^st^ biodiversity research. Proceedings National Academy of Sciences, 116: 22651–22656.

26. Dillman C. B., P. Zhuang, T. Zhang, L.-Z. Zhang, N. Mugue, E. J. Hilton. 2014. Forensic investigations into a GenBank anomaly: endangered taxa and the importance of voucher specimens in molecular studies. Journal of Applied Ichthyology, 30: 1300–1309.

27. Locatelli N. S., P. B. McIntyre, N. O. Therkildsen, & D. S. Baetscher. 2020. GenBank’s reliability is uncertain for biodiversity researchers seeking species-level assignment for eDNA. Proceedings of the National Academy of Sciences of the United States of America, 117:32211–32212.

28. Jerde C. L., E. A. Wilson, & T. L. Dressler. 2019. Measuring global fish species richness with eDNA metabarcoding. Molecular Ecology Resources, 19: 19–22.

29. Nobile A. B., D. Freitas-Souza, F.J. Ruiz-Ruano, M. L. M. O. Nobile, G. O. Costa, F. P. de Lima, J. P. M. Camacho, F. Foresti, & C. Oliveira. 2019. DNA metabarcoding of Neotropical ichthyoplankton: enabling high accuracy with lower cost. Metabarcoding and Metagenomics, 3: e.35060.

30. Cilleros K., A. Valentini, A. Cerdan, T. dejean, A. Iribar, P. Taberlet, R. Vigouroux, & R. Brosse. 2019. Unlocking biodiversity and conservation studies in high diversity environments using environmental DNA (eDNA): a text with Guianese freshwater fishes. Molecular Ecology Resources, 19: 27–46.

31. Sales N. G., O. S. Wangensteen, D. C. Carvalho, & S. Mariani. 2019. Influence of preservation methods, sample medium and sampling time on eDNA recovery in a neotropical river. Environmental DNA, 1: 119–130.

32. Jackman J. M. C. et al. 2021. eDNA in a bottleneck: obstacles to fish metabarcoding studies in megadiverse freshwater systems. Environmental DNA: 10.1002/edna3.191

33. Valentini A., et al. 2016. Next-generation monitoring of aquatic biodiversity using environmental DNA metabarcoding. Molecular Ecology, 25: 929–942.

34. McElroy M. E., et al. 2020. Calibrating environmental DNA metabarcoding to conventional surveys for measuring fish species richness. Frontiers in Ecology and Evolution, 8: 276.

35. Dudgeon D. 2020. Freshwater Biodiversity: Status, Threats and Conservation. Cambridge University Press.

36. Clarke K. R. 1993. Non-parametric multivariate analyses of changes in community structure. Austral Ecology, 18: 117–143.

37. Milan D. T., I. S. Mendes & D. C. Carvalho. 2020. New 12S metabarcoding primers for enhanced Neotropical freshwater fish biodiversity assessment. Scientific Reports, 10: 17966.

38. Deagle B. E., S. N. Jarman, E. Coissac, F. Pompanon, & P. Taberlet. 2014. DNA metabarcoding and the cytochrome c oxidase subunit I marker: Not a perfect match. Biology Letters, 10: 20140562.

39. Collins R. A., J. Bakker, O. S. Wangensteen, A. Z. Soto, L. Corrigan, D. W. Sims, M. J. Genner, & S. Mariani. 2019. Non□specific amplification compromises environmental DNA metabarcoding with COI. Methods in Ecology and Evolution, 10: 1985–2001.

40. Antich A., et al. 2021. To denoise or to cluster, that is not the question: optimizing pipelines for COI metabarcoding and metaphylogeography. BMC Bioinformatics 22:177.

41. Vieira T. B., C. S. Pavanelli, L. Casatti, W.S. Smith, E. Benedito, & R. Mazzoni. 2018. A multiple hypothesis approach to explain species richness patterns in neotropical stream-dweller fish communities. Plos One, 13: e0204114.

42. Zuanon J., F. A. Bockmann, & I. Sazima. 2006. A remarkable sand-dwelling fish assemblage from central Amazonia, with comments on the evolution of psammophily in South American freshwater fishes. Neotropical Ichthyology, 4: 107–118.

43. Sazima I., Carvalho L. N., Mendonça F. P., & Zuanon J. 2006. Fallen leaves on the water-bed: diurnal camouflage of three night-active fish species in an Amazonian streamlet. Neotropical Ichthyology, 4: 119–122.

44. Espírito-Santo H. M. V., & J. Zuanon. 2017. Temporary pools provide stability to fish assemblages in Amazon headwater streams. Ecology of Freshwater Fish, 26: 475–483.

45. de Pinna, M. C. C., J. Zuanon, L. R. Rapp-Py-Daniel, & P. Petry. 2018. A new family of neotropical freshwater fishes from deep fossorial Amazonian habitat, with a reappraisal of morphological characiform phylogeny (Teleostei: Ostariophysi). Zoological Journal of the Linnean Society, 182: 76–106.

46. López-Rojas H., J. G. Lundberg, & E. Marsh. 1984. Design and operation of a small trawling apparatus for use with dugout canoes. North American Journal of Fisheries Management, 4:331–334.

47. Marrero C. & D. C. Taphorn. 1991. Notas sobre la historia natural y la distribution de los peces Gymnotiformes in la cuenca del Rio Apure y otros rios de la Orinoquia. Biollania, 8: 123–142.

48. Cox-Fernandes C., J. Podos, & J. G. Lundberg. 2004. Amazonian Ecology: tributaries enhance the diversity of electric fishes. Science, 305: 1960–1962.

49. Peixoto L. A. W., G. M. Dutra, & W. B. Wosiack. 2015. The electric. Glassknife fishes of the *Eigenmannia trilineata* group (Gymnotiformes: Sternopygidae): monophyly and description of seven new species. Zoological Journal of the Linnean Society, 175: 384–414.

50. de Santana C. D. & R. P. Vari. 2010. Electric fishes of the genus *Sternarchorhynchus* (Teleostei, Ostariophysi, Gymnotiformes); phylogenetic and revisionary studies. Zoological Journal of the Linnean Society, 159: 223–371.

51. Castro R. M. C. 1999. Evolução da ictiofauna de riachos sul-americanos: padrões gerais e possíveis processos causais. In: Caramaschi EP, Mazzoni R, Peres-Neto PR. (Eds) Ecologia de peixes de riachos. Série Oecologia Brasiliensis volume VI, PPGE-UFRJ, Rio de Janeiro, 139–155.

52. Mojica J. I., Castellanos C. & J. Lobón-Cerviá. 2009. High temporal species turnover enhances the complexity of fish assemblages in Amazonian Terra firme streams. Ecology of Freshwater Fish, 18: 518–526.

53. de Oliveira R. R., M. M. Rocha, M. B. Anjos, J. Zuanon, & L. H. Rapp Py-Daniel. 2009. Fish fauna of small streams of the Catua-Ipixuna Extractive Reserve, State of Amazonas, Brazil. Check List, 5: 154–172.

54. Caramaschi E., P. R. Mazzoni, C. R. S. F. Bizerril, & P. R. Peres-Neto. 1999. Ecologia de Peixes de Riachos: Estado Atual e Perspectivas. Oecologia Brasiliensis, v. VI, Rio de Janeiro.

55. Anjos M. B. & J. Zuanon. 2007. Sampling effort and fish species richness in small Terra firme forest streams of central Amazonia, Brazil. Neotropical Ichthyology, 5: 45–52.

56. Mojica J. I., J. Lobón-Cerviá, & C. Castellanos. 2014. Quantifying fish species richness and abundance in Amazonian streams: assessment of a multiple gear method suitable for Terra firme stream fish assemblages. Fisheries Management and Ecology, 21: 220–233.

57. Barros D. F., J. Zuanon, F. P. Mendonça, H. M. V. Espirito-Santo, A. V. Galuch, & A. L. M Albernaz. 2011. The fish fauna of streams in the Madeira-Purus interfluvial region, Brazilian Amazon. Check List, 7: 768–773.

58. Escobar-Camacho D., R. Barriga, & S. R. Ron. 2015. Discovering Hidden Diversity of Characins (Teleostei: Characiformes) in Ecuador’s Yasuní National Park. Plos One, 10: e0135569.

59. Ramirez J. L, J. L. Birindelli, D. C. Carvalho, P. R. A. M. Affonso, P. C. Venere, H. Ortega, M. Carrillo-Avila, J.A. Rodríguez-Pulido, & P. M. Galetti Jr. 2017. Revealing Hidden Diversity of the Underestimated Neotropical Ichthyofauna: DNA Barcoding in the Recently Described Genus *Megaleporinus* (Characiformes: Anostomidae). Frontiers in Genetics, 8: 149.

60. Crampton W. G. R., C. D. de Santana, J. C. Waddell, & N. R. Lovejoy. 2016. The Neotropical electric fish genus *Brachyhypopomus* (Ostariophysi: Gymnotiformes: Hypopomidae): taxonomy and biology, with descriptions of 15 new species. Neotropical Ichthyology, 14: 639–790.

61. Abel R. 2002. Conservation biology for the biodiversity crisis: a freshwater follow-up. Conservation Biology, 5: 1435–1437.

62. Dudgeon D. 2010. Prospects for sustaining freshwater biodiversity in the 21st century: linking ecosystem structure and function. Current Opinion in Environmental Sustainability, 5: 422–430.

63. Jenkins M. 2003. Prospects for biodiversity. Science, 302: 1175–1177.

64. Bunn S. E., C. A. Sullivan, C. R. Liermann, P. M. Davies, C. J. Vörösmarty, P. B. McIntire, M. O. Gessner, D. Dudgeon, A. Prusevich, P. Green & S. Glidden. 2010. Global threats to human water security and river biodiversity. Nature, 467: 555–561.

65. Albert J. S., et al. 2020. Scientists’ warning to humanity on the freshwater biodiversity crisis. Ambio, 50: 85–94.

66. Gilbert M. T. P., T. Haselkorn, M. Bunce, J. J. Sanchez, S. B. Lucas, L. D. Jewell, & M. Worobey. 2007. The isolation of nucleic acids from fixed, paraffin-embedded tissues– which methods are useful when? Plos One, 2: e537.

67. Campos P. F., & T. M. Gilbert. 2012. DNA extraction from formalin-fixed material. In Ancient DNA (pp. 81–85). Humana Press.

68. Hykin S. M., K. Bi, & J. A. McGuire. 2015. Fixing formalin: a method to recover genomic-scale DNA sequence data from formalin-fixed museum specimens using high-throughput sequencing. Plos One, 10: e0141579.

69. Hagedorn M. M., J. P. Daly, V. L. Carter, K. S. Cole, Z. Jaafar, C. V. Lager & L. R. Parenti. 2018. Cryopreservation of fish spermatogonial cells: the future of natural history collections. Scientific Reports, 8: 6149.

70. Albert J. & R. E. Reis. 2011. Historical biogeography of Neotropical freshwater fishes. University of California Press, Berkeley and Los Angeles, California.

71. Sabaj Pérez M. H. 2015. Where the Xingu bends and will soon break. American Scientist, 103: 395–403.

72. Amigo I. 2020. When will the Amazon hit a tipping point? Nature, 578: 505–507.

73. Murienne J., I. Cantera, A. Cerdan, J.B. Decotte, T. Dejean, R. Vigouroux, & S. Brosse. 2019. Aquatic DNA for monitoring French Guiana biodiversity. Biodiversity Data Journal, 7: e37518.

74. McDevitt A. D., et al. 2019. Environmental DNA metabarcoding as an effective and rapid tool for fish monitoring in canals. Journal of Fish Biology, 95: 679–682.

75. Fernandes G. W., M. M. Vale, G. E. Overbeck, M. M. Bustamante, C. E. Grelle, H. G. Bergallo & J. Araújo. 2017. Dismantling Brazil’s science threatens global biodiversity heritage. Perspectives in Ecology and Conservation, 15: 239–243.

76. Alves R. J. V., M. Weksler, J. A. Oliveira, P. A. Buckup, H. R. Santana, A. L. Peracchi & A. L. Albernaz. 2018. Brazilian legislation on genetic heritage harms Biodiversity Convention goals and threatens basic biology research and education. Anais da Academia Brasileira de Ciências, 90: 1279–1284.

77. Overbeck G. E., H. G. Bergallo, C. E. Grelle, A. Akama, F. Bravo, G. R. Colli & G. W. Fernandes. 2018. Global biodiversity threatened by science budget cuts in Brazil. BioScience, 68: 11–12.

78. Miya M., T. Minamoto, H. Yamanaka, S. Oka, K. Sato, S. Yamamoto, T. Sado, & H. Doi. 2016. Use of a filter cartridge for filtration of water samples and extraction of environmental DNA. Journal of Visualized Experiments, 117: 54741.

79. Edgar R. C. 2010. Search and clustering orders of magnitude faster than BLAST. Bioinformatics, 26: 2460–2461.

80. Callahan B. J., P. J. McMurdie & S. P. Holmes. 2017. Exact sequence variants should replace operational taxonomic units in marker-gene data analysis. The ISME Journal, 11: 2639–2643.

81. Katoh K. & D. M. Standley. 2013. MAFFT multiple sequence alignment software version 7: Improvements in performance and usability. Molecular Biology and Evolution, 30: 772–780.

82. Kumar S., G. Stecher, & K. Tamura. 2016. MEGA7: Molecular evolutionary genetics analysis version 7.0 for bigger datasets. Molecular Biology and Evolution, 33: 1870–1874.

83. Kimura M. 1980. A simple method for estimating evolutionary rates of base substitutions through comparative studies of nucleotide sequences. Journal of Molecular Evolution, 16: 111–120.

84. Darriba D., G. L. Taboada, R. Doallo, & D. Posada. 2012. jModelTest 2: more models, new heuristics and parallel computing. Nature Methods, 9: 772.

85. Ronquist F., M. Teslenko, P. van der Mark, D.L. Ayres, A. Darling, S. Höhna, B. Larget, L. Liu, M.A. Suchard & J. P. Huelsenbeck. 2012. MrBayes 3.2: efficient Bayesian phylogenetic inference and model choice across a large model space. Systematics Biology, 61: 539–542.

86. Miller M. A., et al. 2015. A RESTful API for Access to Phylogenetic Tools via the CIPRES Science Gateway. Evolutionary Bioinformatics, 11: 43–48.

87. Ciccarelli F. D., T. Goerks, C. von Mering, C. J. Creevery, B. Snel, & P. Bork. 2006. Toward Automatic Reconstruction of a Highly Resolved Tree of Life. Science, 311: 1283–1287.

88. R Core Team. 2020. R: A language and environment for statistical computing. R Foundation for Statistical Computing, Vienna, Austria.https://www.Rproject.org/

89. Wickham H. 2016. Ggplot2: Elegant graphics for data analysis. Houston, TX: Springer.

90. Oksanen J., R. Kindt, & B. O’Hara. 2013. Package VEGAN. Community Ecology Package, Version 2.

91. Fox J. & S. Weisberg. 2019. An R Companion to Applied Regression, Third edition. Sage, Thousand Oaks CA.

92. Adler D., O. Nenadic, & W. Zucchini. 2013. rgl: 3D visualization device system (OpenGL). R package version 0.93.945, URL http://CRAN.R-project.org/package=rgl.

93. Gu Z. 2014. Circlize implements and enhances circular visualization in R. Bioinformatics, 30:2811–2812.

94. Schiettekatte N. M. D., S. J. Brandl, & J. M. Casey. 2019. Fishualize: color palettes based on fish species. CRAN version 0.2.0.

95. Chao A. 1989. Estimating Population Size for Sparse Data in Capture-Recapture Experiments. Biometrics, 45: 427.

96. Hsieh T. C., K. H. Ma, A. Chao. 2020. iNEXT: Interpolation and Extrapolation for Species Diversity. R package version 2.0.20

97. Chao A., R. L. Chazdon, R. K. Colwell, & T.-J. Shen. 2005. A new statistical approach for assessing compositional similarity based on incidence and abundance data. Ecology Letters, 8: 148–15.

98. Olds B. P., C. L. Jerde, M. A. Renshaw, L. Yiuan, N. T. Evans, C. R. Turner, K. Deiner, A. R. Mahon, M. A. Brueseke, P. D. Shirey, M. E. Pfrende, D. M. Lodge, & G. A. Lambert. 2016. Estimating species richness using environmental DNA. Ecology and Evolution, 6: 4214–4226.

99. Chao A., K. H. Ma, T. C. Hsieh, T. C. & C. H. Chiu. 2016. SpadeR (Species-richness Prediction And Diversity Estimation in R): an R package in CRAN. Program and User’s Guide also published at http://chao.stat.nthu.edu.tw/wordpress/software_download/.

