## Supplementary material for "The critical role of natural history museums in advancing eDNA for biodiversity studies: a case study with Amazonian fishes": Supp information

**Supplementary information**

**Species composition among samples and sampling influence on detecting species richness in MBS**

To assess species composition among samples and the influence of sampling on detecting species richness in river and stream, we compared samples collected in stations 1 and 2. Species detection per sample - 100 ml of filtered water - ranged from 62 to 87 in the stream and 34 to 48 species in the river (Supplementary Figure 1A). Stream species richness varied from 3 to 13 species between samples. As for the river, species richness varied by up to 19 species between samples. Species composition was more similar among stream samples than river samples as indicated by Jaccard’s similarity indices, which ranged from 0.49 to 0.61 in the stream and 0.21 to 0.30 in the river (Supplementary Figure 1B) with significantly higher similarity among samples in the stream (Kruskal-Wallis rank sum test χ^2^ = 13.449, p = 0.009) than in the river (Kruskal-Wallis rank sum test χ^2^ = 5.6308, p = 0.228). Thus, in the stream, according to the observed curve of species richness, sample 1 contributed 66.7% of the total species diversity, followed by 19% of sample 2; 5.5% in sample 3; 2.4% in sample 4; and 6.3% in sample 5. In contrast, river sample 1 contribute only 31% of the detected species diversity, samples 2, 3, 4 and 5 contributed 22%, 13%, 21.5%, and 14%, respectively. Based on random species accumulation curves (Supplementary Figure 1C), when samples are combined, five samples or 500 ml of filtered water from each locality, can be considerate a robust estimate of the actual species richness (Supplementary Figure 1C).

### Supplementary Figure 1. Environmental DNA analyses for Javari river (top) and stream (bottom) samples. (A) Violin plot illustrating species abundance per order, probability density (distribution), median, interquartile ranges and total species detected per sample. (B) Heatmap of Jaccard’s similarity coefficients. Note that higher similarity among the five stream samples than in the five river samples. (C) Species accumulation curves. In both figures, blue line are curves calculated based on the sum of species when filter samples are combined. 95% confidence intervals refer to the blue curves and boxplots of these curves show distribution of species diversity as inferred from the method “random”, which add sites in random order and was used for the species accumulation curves.

**Supplementary tables**

**Supplementary table 1.** List and vouchers numbers for 443 species collected in the Javari River basin. CBS = Capture-based sampling; MBS = Molecular-based sampling. S = Station. S1 = Javari River; S2 = Terra firme stream; S3 = Quixito River.

**Supplementary Table 2.** Summary of read numbers after data preprocessing (the second to fifth columns) and after taxon assignment (the last two columns) for the 11 libraries and three blanks. The numbers in parentheses in the third to fifth columns are percentages of the raw read numbers, while those in the last two columns are percentages of denoised read numbers. EB, 1B, and 2B indicate blank samples from DNA extraction, 1st PCR, and 2nd PCR, respectively.

**Supplementary Table 3.** Summary of environmental DNA data analyses for 11 samples collected at the Javari River basin. Station 1 = Javari River; Station 2 = Terra firme stream; Station 3 = Quixito River.

**Supplementary Table 4.** Representative 12S rRNA sequences for the 222 taxa detected by molecular-based capture in the three stations at the Javari River Basin.

**Supplementary Table 5.** List and abundance of species detected by Capture-based sampling in the Javari River – Station 1.

**Supplementary Table 6.** List and frequency of species detected by Molecular-based sampling in the Javari River – Station 1.

**Supplementary Table 7.** List and abundance of species detected by Capture-based sampling in the Terra firme stream – Station 2.

**Supplementary Table 8.** List and frequency of species detected by Molecular-based sampling in the Terra firme stream – Station 2.

**Supplementary Table 9.** Contamination list from non-Amazonian fishes removed from analyses.
