## Supplementary figures and images for "The critical role of natural history museums in advancing eDNA for biodiversity studies: a case study with Amazonian fishes"

### Figure S1

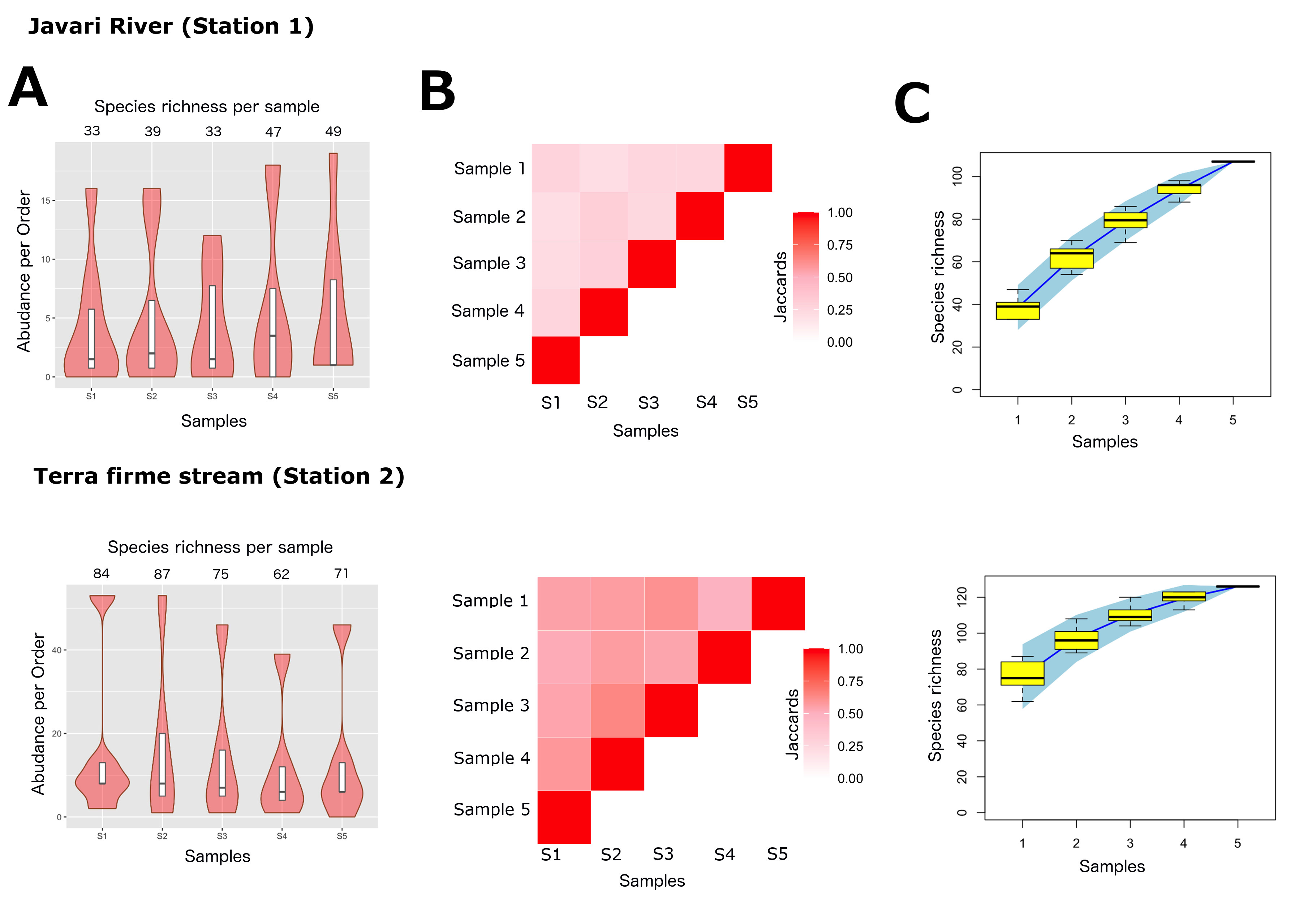
