## Supplementary material for "The critical role of natural history museums in advancing eDNA for biodiversity studies: a case study with Amazonian fishes": Table S1

| **Order** | **Family** | **Genus** | **Species** | **Authority** | **Voucher** | **CBS** |
| --- | --- | --- | --- | --- | --- | --- |
| Characiformes | Acestrorhynchidae | *Acestrorhynchus* | *abbreviatus* | (Cope 1878) | INPA-ICT 055385 | S1 |
| Characiformes | Acestrorhynchidae | *Acestrorhynchus* | *falcatus* | (Bloch 1794) | INPA-ICT 055991 | S2 |
| Characiformes | Acestrorhynchidae | *Acestrorhynchus* | *falcirostris* | (Cuvier 1819) | INPA-ICT 055386 | S1 |
| Characiformes | Acestrorhynchidae | *Acestrorhynchus* | *heterolepis* | (Cope 1878) | INPA-ICT 055182 |  |
| Characiformes | Acestrorhynchidae | *Acestrorhynchus* | *microlepis* | (Jardine 1841) | INPA-ICT 055387 | S1 |
| Pleuronectiformes | Achiridae | *Apionichthys* | *nattereri* | (Steindachner 1876) | INPA-ICT 055487 | S1 |
| Pleuronectiformes | Achiridae | *Hypoclinemus* | *mentalis* | (Günther 1862) | INPA-ICT 055269 | S1, S3 |
| Characiformes | Chalceidae | *Chalceus* | *epakros* | Zanata & Toledo-Piza 2004 | INPA-ICT 055264 |  |
| Characiformes | Chalceidae | *Chalceus* | *erythrurus* | (Cope 1870) | INPA-ICT 055360 | S1 |
| Characiformes | Anostomidae | *Abramites* | *hypselonotus* | (Günther 1868) | INPA-ICT 056093 |  |
| Characiformes | Anostomidae | *Anostomoides* | *atrianalis* | Pellegrin 1909 | INPA-ICT 055393 | S1, S3 |
| Characiformes | Anostomidae | *Laemolyta* | *proxima* | (Garman 1890) | INPA-ICT 056347 |  |
| Characiformes | Anostomidae | *Leporinus* | *agassizii* | Steindachner 1876 | INPA-ICT 055244 | S1, S2 |
| Characiformes | Anostomidae | *Leporinus* | *cylindriformis* | Borodin 1929 | INPA-ICT 055440 | S1, S2, S3 |
| Characiformes | Anostomidae | *Leporinus* | *jatuncochi* | Ovchynnyk 1971 | INPA-ICT 055822 |  |
| Characiformes | Anostomidae | *Leporinus* | *fasciatus* | (Bloch 1794) | INPA-ICT 056126 |  |
| Characiformes | Anostomidae | *Leporinus* | *friderici* | (Bloch 1794) | INPA-ICT 055803 |  |
| Characiformes | Anostomidae | *Leporinus* | *jamesi* | Garman 1929 | INPA-ICT 056127 |  |
| Characiformes | Anostomidae | *Pseudanos* | *trimaculatus* | (Kner 1858) | INPA-ICT 055763 | S2 |
| Characiformes | Anostomidae | *Rhytiodus* | *argenteofuscus* | Kner 1858 | INPA-ICT 056153 |  |
| Characiformes | Anostomidae | *Rhytiodus* | *microlepis* | Kner 1858 | INPA-ICT 055876 |  |
| Characiformes | Anostomidae | *Schizodon* | *fasciatus* | Spix & Agassiz 1829 | INPA-ICT 055546 | S1 |
| Gymnotiformes | Apteronotidae | *Adontosternarchus* | *balaenops* | (Cope 1878) | INPA-ICT 055317 | S1 |
| Gymnotiformes | Apteronotidae | *Adontosternarchus* | *clarkae* | Mago-Leccia, Lundberg & Baskin 1985 | INPA-ICT 055312 | S1 |
| Gymnotiformes | Apteronotidae | *Adontosternarchus* | *nebulosus* | Lundberg & Cox Fernandes 2007 | INPA-ICT 055344 |  |
| Gymnotiformes | Apteronotidae | *Adontosternarchus* | *sachsi* | (Peters 1877) | INPA-ICT 055318 | S1 |
| Gymnotiformes | Apteronotidae | *Apteronotus* | *bonapartii* | (Castelnau 1855) | INPA-ICT 055959 |  |
| Gymnotiformes | Apteronotidae | *Melanosternarchus* | *amaru* | Bernt, Crampton, Orfinger & Albert 2018 | INPA-ICT 055933 |  |
| Gymnotiformes | Apteronotidae | *Compsaraia* | *compsa* | (Mago-Leccia 1994) | INPA-ICT 055506 | S1 |
| Gymnotiformes | Apteronotidae | *Platyurosternarchus* | *macrostomus* | (Günther 1870) | INPA-ICT 056232 |  |
| Gymnotiformes | Apteronotidae | *Porotergus* | *gimbeli* | Ellis 1912 | INPA-ICT 055514 | S1 |
| Gymnotiformes | Apteronotidae | *Sternarchella* | *calhamazon* | Lundberg, Cox Fernandes, Campos-da-Paz & Sullivan 2013 | INPA-ICT 055177 |  |
| Gymnotiformes | Apteronotidae | *Sternarchella* | *orthos* | Mago-Leccia 1994 | INPA-ICT 055178 |  |
| Gymnotiformes | Apteronotidae | *Sternarchella* | *schotti* | (Steindachner 1868) | INPA-ICT 055179 |  |
| Gymnotiformes | Apteronotidae | *Sternarchella* | *sima* | Starks 1913 | INPA-ICT 055694 |  |
| Gymnotiformes | Apteronotidae | *Sternarchogiton* | *nattereri* | (Steindachner 1868) | INPA-ICT 055333 |  |
| Gymnotiformes | Apteronotidae | *Sternarchogiton* | *porcinum* | Eigenmann & Allen 1942 | INPA-ICT 055334 |  |
| Gymnotiformes | Apteronotidae | *Tenebrosternarchus* | *preto* | (de Santana & Crampton 2007) | INPA-ICT 055968 |  |
| Gymnotiformes | Apteronotidae | *Sternarchorhamphus* | *muelleri* | (Steindachner 1881) | INPA-ICT 055919 |  |
| Gymnotiformes | Apteronotidae | *Sternarchorhynchus* | *cramptoni* | de Santana & Vari 2010 | INPA-ICT 055335 | S1 |
| Gymnotiformes | Apteronotidae | *Sternarchorhynchus* | *goeldii* | de Santana & Vari 2010 | INPA-ICT 055520 |  |
| Gymnotiformes | Apteronotidae | *Sternarchorhynchus* | *montanus* | de Santana & Vari 2010 | INPA-ICT 055695 |  |
| Osteoglossiformes | Arapaimidae | *Arapaima* | *gigas* | (Schinz 1822) | INPA-ICT 055148 |  |
| Siluriformes | Aspredinidae | *Bunocephalus* | *aleuropsis* | Cope 1870 | INPA-ICT 055319 |  |
| Siluriformes | Aspredinidae | *Bunocephalus* | *coracoideus* | (Cope 1874) | INPA-ICT 055194 |  |
| Siluriformes | Aspredinidae | *Bunocephalus* | *verrucosus* | (Walbaum 1792) | INPA-ICT 055833 |  |
| Siluriformes | Aspredinidae | *Pseudobunocephalus* | *amazonicus* | (Mees 1989) | INPA-ICT 055859 |  |
| Siluriformes | Aspredinidae | *Pseudobunocephalus* | *quadriradiatus* | (Mees 1989) | INPA-ICT 056185 |  |
| Siluriformes | Aspredinidae | *Pseudobunocephalus* | sp. |  | INPA-ICT 056186 |  |
| Siluriformes | Aspredinidae | *Pterobunocephalus* | cf. *dolichurus* | (Delsman 1941) | INPA-ICT 055560 |  |
| Siluriformes | Aspredinidae | *Pterobunocephalus* | *depressus* | (Haseman 1911) | INPA-ICT 056572 |  |
| Siluriformes | Auchenipteridae | *Ageneiosus* | *dentatus* | Kner 1858 | INPA-ICT 055642 |  |
| Siluriformes | Auchenipteridae | *Ageneiosus* | *inermis* | (Linnaeus 1766) | INPA-ICT 055389 | S1 |
| Siluriformes | Auchenipteridae | *Ageneiosus* | *lineatus* | Ribeiro, Rapp Py-Daniel & Walsh 2017 | INPA-ICT 055505 | S1 |
| Siluriformes | Auchenipteridae | *Ageneiosus* | *ucayalensis* | Castelnau 1855 | INPA-ICT 055298 | S1 |
| Siluriformes | Auchenipteridae | *Ageneiosus* | *vittatus* | Steindachner 1908 | INPA-ICT 056471 |  |
| Siluriformes | Auchenipteridae | *Auchenipterichthys* | *coracoideus* | (Eigenmann & Allen 1942) | INPA-ICT 055262 | S1, S3 |
| Siluriformes | Auchenipteridae | *Auchenipterus* | *brachyurus* | (Cope 1878) | INPA-ICT 055398 | S1, S3 |
| Siluriformes | Auchenipteridae | *Auchenipterus* | *fordicei* | Eigenmann & Eigenmann 1888 | INPA-ICT 056243 |  |
| Siluriformes | Auchenipteridae | *Auchenipterus* | *nuchalis* | (Spix & Agassiz 1829) | INPA-ICT 055526 | S1 |
| Siluriformes | Auchenipteridae | *Centromochlus* | *existimatus* | Mees 1974 | INPA-ICT 055406 | S1 |
| Siluriformes | Auchenipteridae | *Epapterus* | *dispilurus* | Cope 1878 | INPA-ICT 055421 | S1, S3 |
| Siluriformes | Auchenipteridae | *Trachelyopterus* | *galeatus* | (Linnaeus 1766) | INPA-ICT 056730 | S1 |
| Siluriformes | Auchenipteridae | *Centromochlus* | aff. *melanoleuca* | (Vari & Calegari 2014) | INPA-ICT 055987 |  |
| Siluriformes | Auchenipteridae | *Tatia* | *intermedia* | (Steindachner 1877) | INPA-ICT 055214 | S1 |
| Siluriformes | Auchenipteridae | *Tympanopleura* | *atronasus* | (Eigenmann & Eigenmann 1888) | INPA-ICT 055661 |  |
| Siluriformes | Auchenipteridae | *Tympanopleura* | *brevis* | (Steindachner 1881) | INPA-ICT 055180 | S1 |
| Siluriformes | Auchenipteridae | *Tympanopleura* | *piperata* | Eigenmann 1912 | INPA-ICT 055181 | S1 |
| Beloniformes | Belonidae | *Potamorrhaphis* | *guianensis* | (Jardine 1843) | INPA-ICT 055254 | S2 |
| Beloniformes | Belonidae | *Pseudotylosurus* | *angusticeps* | (Günther 1866) | INPA-ICT 055380 |  |
| Beloniformes | Belonidae | *Pseudotylosurus* | *microps* | (Günther 1866) | INPA-ICT 055462 | S1 |
| Beloniformes | Belonidae | *Pseudotylosurus* | sp. |  | INPA-ICT 055898 |  |
| Siluriformes | Callichthyidae | *Corydoras* | aff. *trilineatus* | Cope 1872 | INPA-ICT 056429 |  |
| Siluriformes | Callichthyidae | *Corydoras* | *agassizii* | Steindachner 1876 | INPA-ICT 056174 |  |
| Siluriformes | Callichthyidae | *Corydoras* | *ambiacus* | Cope 1872 | INPA-ICT 056005 | S2 |
| Siluriformes | Callichthyidae | *Corydoras* | *granti* | Tencatt, Lima & Britto 2019 | INPA-ICT 055196 | S2 |
| Siluriformes | Callichthyidae | *Corydoras* | *armatus* | (Günther 1868) | INPA-ICT 055197 |  |
| Siluriformes | Callichthyidae | *Corydoras* | *elegans* | Steindachner 1876 | INPA-ICT 056487 |  |
| Siluriformes | Callichthyidae | *Corydoras* | *melanistius* | Regan 1912 | INPA-ICT 056007 | S2 |
| Siluriformes | Callichthyidae | *Corydoras* | *reticulatus* | Fraser-Brunner 1938 | INPA-ICT 056008 | S2 |
| Siluriformes | Callichthyidae | *Corydoras* | *sodalis* | Nijssen & Isbrücker 1986 | INPA-ICT 056333 |  |
| Siluriformes | Callichthyidae | *Corydoras* | *zygatus* | Eigenmann & Allen 1942 | INPA-ICT 056176 |  |
| Siluriformes | Callichthyidae | *Dianema* | *longibarbis* | Cope 1872 | INPA-ICT 056177 |  |
| Siluriformes | Callichthyidae | *Lepthoplosternum* | *altamazonicum* | Reis 1997 | INPA-ICT 056181 |  |
| Siluriformes | Callichthyidae | *Lepthoplosternum* | *beni* | Reis 1997 | INPA-ICT 055804 |  |
| Siluriformes | Callichthyidae | *Megalechis* | *thoracata* | (Valenciennes 1840) | INPA-ICT 056183 |  |
| Siluriformes | Callichthyidae | *Megalechis* | *picta* | (Müller & Troschel 1849) | INPA-ICT 056223 |  |
| Siluriformes | Cetopsidae | *Cetopsis* | *candiru* | Spix & Agassiz 1829 | INPA-ICT 056654 |  |
| Siluriformes | Cetopsidae | *Cetopsis* | *coecutiens* | (Lichtenstein 1819) | INPA-ICT 055934 |  |
| Characiformes | Characidae | *Aphyocharax* | *avary* | Fowler 1913 | INPA-ICT 055216 | S1, S3 |
| Characiformes | Characidae | *Aphyocharax* | sp. |  | INPA-ICT 055701 |  |
| Characiformes | Characidae | *Aphyodite* | *grammica* | Eigenmann 1912 | INPA-ICT 056370 |  |
| Characiformes | Characidae | *Astyanax* | aff. *elachylepis* | Bertaco & Lucinda 2005 | INPA-ICT 055994 | S2 |
| Characiformes | Characidae | *Makunaima* | aff. *multidens* | (Eigenmann 1908) | INPA-ICT 055217 |  |
| Characiformes | Characidae | *Jupiaba* | *anterior* | (Eigenmann 1908) | INPA-ICT 055479 |  |
| Characiformes | Characidae | *Astyanax* | *bimaculatus* | (Linnaeus 1758) | INPA-ICT 056032 | S2 |
| Characiformes | Characidae | *Makunaima* | cf. *guianensis* | (Eigenmann 1909) | INPA-ICT 055188 | S1 |
| Characiformes | Characidae | *Brachychalcinus* | *copei* | (Steindachner 1882) | INPA-ICT 055995 | S2 |
| Characiformes | Bryconidae | *Brycon* | aff. *pesu* | Müller & Troschel 1845 | INPA-ICT 055402 | S1 |
| Characiformes | Bryconidae | *Brycon* | *amazonicus* | (Agassiz 1829) | INPA-ICT 055193 | S2 |
| Characiformes | Bryconidae | *Brycon* | *melanopterus* | (Cope 1872) | INPA-ICT 055998 | S2 |
| Characiformes | Characidae | *Bryconella* | *pallidifrons* | (Fowler 1946) | INPA-ICT 055787 |  |
| Characiformes | Iguanodectidae | *Bryconops* | *alburnoides* | Kner 1858 | INPA-ICT 055403 | S1 |
| Characiformes | Iguanodectidae | *Bryconops* | *caudomaculatus* | (Günther 1864) | INPA-ICT 055999 | S2 |
| Characiformes | Iguanodectidae | *Bryconops* | *inpai* | Knöppel, Junk & Géry 1968 | INPA-ICT 056000 | S2 |
| Characiformes | Iguanodectidae | *Bryconops* | *magoi* | Chernoff & Machado-Allison 2005 | INPA-ICT 055218 |  |
| Characiformes | Characidae | *Charax* | *michaeli* | Lucena 1989 | INPA-ICT 055293 |  |
| Characiformes | Characidae | *Charax* | *tectifer* | (Cope 1870) | INPA-ICT 056712 |  |
| Characiformes | Characidae | *Creagrutus* | *cochui* | Géry 1964 | INPA-ICT 056036 | S2 |
| Characiformes | Characidae | *Ctenobrycon* | *spilurus* | (Valenciennes 1850) | INPA-ICT 055153 | S1 |
| Characiformes | Characidae | *Cynopotamus* | *amazonum* | (Günther 1868) | INPA-ICT 055586 |  |
| Characiformes | Characidae | *Galeocharax* | *gulo* | (Cope 1870) | INPA-ICT 055530 | S1 |
| Characiformes | Acestrorhynchidae | *Gnathocharax* | *steindachneri* | Fowler 1913 | INPA-ICT 056498 |  |
| Characiformes | Characidae | *Gymnocorymbus* | *thayeri* | Eigenmann 1908 | INPA-ICT 056011 | S2 |
| Characiformes | Characidae | *Hemigrammus* | aff. *geisleri* | Zarske & Géry 2007 | INPA-ICT 055231 |  |
| Characiformes | Characidae | *Hemigrammus* | aff. *lunatus* | Durbin 1918 | INPA-ICT 056339 |  |
| Characiformes | Characidae | *Hemigrammus* | *analis* | Durbin 1909 | INPA-ICT 056500 |  |
| Characiformes | Characidae | *Hemigrammus* | *bellottii* | (Steindachner 1882) | INPA-ICT 055202 |  |
| Characiformes | Characidae | *Hemigrammus* | *geisleri* | Zarske & Géry 2007 | INPA-ICT 055154 |  |
| Characiformes | Characidae | *Hemigrammus* | *haraldi* | Géry 1961 | INPA-ICT 055233 |  |
| Characiformes | Characidae | *Hemigrammus* | *lunatus* | Durbin 1918 | INPA-ICT 055234 |  |
| Characiformes | Characidae | *Hemigrammus* | *machadoi* | Ota, Lima & Pavanelli 2014 | INPA-ICT 056504 |  |
| Characiformes | Characidae | *Hemigrammus* | *newboldi* | (Fernández-Yépez 1949) | INPA-ICT 055773 |  |
| Characiformes | Characidae | *Hemigrammus* | *ocellifer* | (Steindachner 1882) | INPA-ICT 055203 |  |
| Characiformes | Characidae | *Hemigrammus* | *schmardae* | (Steindachner 1882) | INPA-ICT 056392 |  |
| Characiformes | Characidae | *Hemigrammus* | sp. |  | INPA-ICT 055848 |  |
| Characiformes | Characidae | *Hemigrammus* | *unilineatus* | (Gill 1858) | INPA-ICT 055235 |  |
| Characiformes | Acestrorhynchidae | *Heterocharax* | *macrolepis* | Eigenmann 1912 | INPA-ICT 056507 |  |
| Characiformes | Characidae | *Hyphessobrycon* | aff. *bentosi* | Durbin 1908 | INPA-ICT 056344 |  |
| Characiformes | Characidae | *Hyphessobrycon* | aff. *heterorhabdus* | (Ulrey 1894) | INPA-ICT 056016 | S2 |
| Characiformes | Characidae | *Hyphessobrycon* | *agulha* | Fowler 1913 | INPA-ICT 056017 | S2 |
| Characiformes | Characidae | *Hyphessobrycon* | *bentosi* | Durbin 1908 | INPA-ICT 055236 |  |
| Characiformes | Characidae | *Hyphessobrycon* | sp. |  | INPA-ICT 055237 | S2 |
| Characiformes | Characidae | *Hyphessobrycon* | *dorsalis* | Zarske 2014 | INPA-ICT 055887 |  |
| Characiformes | Characidae | *Hyphessobrycon* | *hasemani* | Fowler 1913 | INPA-ICT 055238 |  |
| Characiformes | Characidae | *Hyphessobrycon* | *peruvianus* | Ladiges 1938 | INPA-ICT 055239 |  |
| Characiformes | Characidae | *Hyphessobrycon* | sp. 1 |  | INPA-ICT 055798 |  |
| Characiformes | Characidae | *Hyphessobrycon* | *sweglesi* | (Géry 1961) | INPA-ICT 055240 |  |
| Characiformes | Iguanodectidae | *Iguanodectes* | *purusii* | (Steindachner 1908) | INPA-ICT 055751 |  |
| Characiformes | Iguanodectidae | *Iguanodectes* | *spilurus* | (Günther 1864) | INPA-ICT 055597 | S2 |
| Characiformes | Characidae | *Knodus* | *aff. megalops* | Myers 1929 | INPA-ICT 056719 |  |
| Characiformes | Characidae | *Knodus* | *cf. heteresthes* | (Eigenmann 1908) | INPA-ICT 056263 |  |
| Characiformes | Characidae | *Knodus* | *smithi* | (Fowler 1913) | INPA-ICT 055158 | S1, S3 |
| Characiformes | Characidae | *Protocheirodon* | *pi* | (Vari 1978) | INPA-ICT 055274 | S1 |
| Characiformes | Characidae | *Moenkhausia* | *aff. cotinho* | Eigenmann 1908 | INPA-ICT 055246 |  |
| Characiformes | Characidae | *Moenkhausia* | *ceros* | Eigenmann 1908 | INPA-ICT 056225 |  |
| Characiformes | Characidae | *Moenkhausia* | *chrysargyrea* | (Günther 1864) | INPA-ICT 056041 | S2 |
| Characiformes | Characidae | *Moenkhausia* | *collettii* | (Steindachner 1882) | INPA-ICT 055247 | S2 |
| Characiformes | Characidae | *Moenkhausia* | *comma* | Eigenmann 1908 | INPA-ICT 056721 |  |
| Characiformes | Characidae | *Moenkhausia* | *cotinho* | Eigenmann 1908 | INPA-ICT 055444 | S1 |
| Characiformes | Characidae | *Moenkhausia* | *gracilima* | Eigenmann 1908 | INPA-ICT 055160 | S1, S3 |
| Characiformes | Characidae | *Moenkhausia* | *grandisquamis* | (Müller & Troschel 1845) | INPA-ICT 055277 | S1, S2 |
| Characiformes | Characidae | *Moenkhausia* | *intermedia* | Eigenmann 1908 | INPA-ICT 055613 |  |
| Characiformes | Characidae | *Moenkhausia* | *jamesi* | Eigenmann 1908 | INPA-ICT 055447 | S1 |
| Characiformes | Characidae | *Moenkhausia* | *oligolepis* | (Günther 1864) | INPA-ICT 055207 | S2, S3 |
| Characiformes | Characidae | *Moenkhausia* | sp. 1 |  | INPA-ICT 055615 | S2 |
| Characiformes | Characidae | *Moenkhausia* | sp. 2 |  | INPA-ICT 055448 | S1 |
| Characiformes | Characidae | *Odontostilbe* | *fugitiva* | Cope 1870 | INPA-ICT 055162 |  |
| Characiformes | Characidae | *Odontostilbe* | *nareuda* | Bührnheim & Malabarba 2006 | INPA-ICT 055759 |  |
| Characiformes | Characidae | *Paracheirodon* | *innesi* | (Myers 1936) | INPA-ICT 055808 |  |
| Characiformes | Characidae | *Paragoniates* | *alburnus* | Steindachner 1876 | INPA-ICT 055279 | S1 |
| Characiformes | Characidae | *Phenacogaster* | *aff. beni* | Eigenmann 1911 | INPA-ICT 055857 | S2 |
| Characiformes | Characidae | *Phenacogaster* | *aff. pectinatus* | (Cope 1870) | INPA-ICT 055760 | S2 |
| Characiformes | Characidae | *Phenacogaster* | *cf. napoatilis* | Lucena & Malabarba 2010 | INPA-ICT 056527 |  |
| Characiformes | Characidae | *Poptella* | *compressa* | (Günther 1864) | INPA-ICT 055762 |  |
| Characiformes | Characidae | *Prionobrama* | *filigera* | (Cope 1870) | INPA-ICT 055167 | S1, S3 |
| Characiformes | Characidae | *Roeboides* | *affinis* | (Günther 1868) | INPA-ICT 055170 | S1, S3 |
| Characiformes | Characidae | *Roeboides* | *biserialis* | (Garman 1890) | INPA-ICT 055625 |  |
| Characiformes | Characidae | *Roeboides* | *myersii* | Gill 1870 | INPA-ICT 055469 | S1 |
| Characiformes | Acestrorhynchidae | *Roestes* | *molossus* | (Kner 1858) | INPA-ICT 056538 |  |
| Characiformes | Characidae | *Stethaprion* | *erythrops* | Cope 1870 | INPA-ICT 055477 | S1 |
| Characiformes | Characidae | *Stichonodon* | *insignis* | (Steindachner 1876) | INPA-ICT 056161 |  |
| Characiformes | Characidae | *Tetragonopterus* | *argenteus* | Cuvier 1816 | INPA-ICT 055171 | S1 |
| Characiformes | Characidae | *Thayeria* | sp. |  | INPA-ICT 056411 |  |
| Characiformes | Triportheidae | *Triportheus* | *albus* | Cope 1872 | INPA-ICT 055482 | S1 |
| Characiformes | Triportheidae | *Triportheus* | *angulatus* | (Spix & Agassiz 1829) | INPA-ICT 055172 | S1, S3 |
| Characiformes | Triportheidae | *Triportheus* | *auritus* | (Valenciennes 1850) | INPA-ICT 055383 | S1 |
| Characiformes | Triportheidae | *Triportheus* | *culter* | (Cope 1872) | INPA-ICT 056164 |  |
| Characiformes | Triportheidae | *Triportheus* | *pictus* | (Garman 1890) | INPA-ICT 056364 | S2 |
| Characiformes | Characidae | *Tyttocharax* | *cochui* | (Ladiges 1949) | INPA-ICT 055819 | S2 |
| Characiformes | Characidae | *Xenurobrycon* | *polyancistrus* | Weitzman 1987 | INPA-ICT 055260 |  |
| Characiformes | Chilodontidae | *Caenotropus* | *labyrinthicus* | (Kner 1858) | INPA-ICT 055404 | S1 |
| Characiformes | Chilodontidae | *Chilodus* | *punctatus* | Müller & Troschel 1844 | INPA-ICT 055220 | S2 |
| Cichliformes | Cichlidae | *Aequidens* | *pallidus* | (Heckel 1840) | INPA-ICT 055992 | S2 |
| Cichliformes | Cichlidae | *Aequidens* | *tetramerus* | (Heckel 1840) | INPA-ICT 055183 |  |
| Cichliformes | Cichlidae | *Apistogramma* | *agassizii* | (Steindachner 1875) | INPA-ICT 055186 |  |
| Cichliformes | Cichlidae | *Apistogramma* | *bitaeniata* | Pellegrin 1936 | INPA-ICT 055786 |  |
| Cichliformes | Cichlidae | *Apistogramma* | *eunotus* | Kullander 1981 | INPA-ICT 055187 |  |
| Cichliformes | Cichlidae | *Apistogramma* | sp. |  | INPA-ICT 056168 |  |
| Cichliformes | Cichlidae | *Astronotus* | *ocellatus* | (Agassiz 1831) | INPA-ICT 055831 |  |
| Cichliformes | Cichlidae | *Biotodoma* | *cupido* | (Heckel 1840) | INPA-ICT 055399 | S1 |
| Cichliformes | Cichlidae | *Bujurquina* | *syspilus* | (Cope 1872) | INPA-ICT 056586 |  |
| Cichliformes | Cichlidae | *Chaetobranchus* | *flavescens* | Heckel 1840 | INPA-ICT 056376 |  |
| Cichliformes | Cichlidae | *Cichla* | *monoculus* | Spix & Agassiz 1831 | INPA-ICT 055410 | S1 |
| Cichliformes | Cichlidae | *Cichlasoma* | *amazonarum* | Kullander 1983 | INPA-ICT 056055 |  |
| Cichliformes | Cichlidae | *Crenicara* | *punctulatum* | (Günther 1863) | INPA-ICT 055198 |  |
| Cichliformes | Cichlidae | *Crenicichla* | *cf. inpa* | Ploeg 1991 | INPA-ICT 056381 |  |
| Cichliformes | Cichlidae | *Crenicichla* | *cincta* | Regan 1905 | INPA-ICT 055411 | S1 |
| Cichliformes | Cichlidae | *Crenicichla* | *inpa* | Ploeg 1991 | INPA-ICT 055199 | S1, S2 |
| Cichliformes | Cichlidae | *Crenicichla* | *johanna* | Heckel 1840 | INPA-ICT 056059 |  |
| Cichliformes | Cichlidae | *Crenicichla* | *regani* | Ploeg 1989 | INPA-ICT 055222 | S2 |
| Cichliformes | Cichlidae | *Crenicichla* | *reticulata* | (Heckel 1840) | INPA-ICT 055413 | S1 |
| Cichliformes | Cichlidae | *Heros* | *efasciatus* | Heckel 1840 | INPA-ICT 056342 |  |
| Cichliformes | Cichlidae | *Hypselecara* | *coryphaenoides* | (Heckel 1840) | INPA-ICT 056594 |  |
| Cichliformes | Cichlidae | *Hypselecara* | *temporalis* | (Günther 1862) | INPA-ICT 055853 |  |
| Cichliformes | Cichlidae | *Laetacara* | *thayeri* | (Steindachner 1875) | INPA-ICT 056514 |  |
| Cichliformes | Cichlidae | *Mesonauta* | *festivus* | (Heckel 1840) | INPA-ICT 056129 |  |
| Cichliformes | Cichlidae | *Pterophyllum* | *scalare* | (Schultze 1823) | INPA-ICT 055860 |  |
| Cichliformes | Cichlidae | *Satanoperca* | sp. |  | INPA-ICT 055863 |  |
| Cupleiformes | Pristigasteridae | *Pellona* | *castelnaeana* | Valenciennes 1847 | INPA-ICT 055873 |  |
| Cupleiformes | Pristigasteridae | *Pristigaster* | *cayana* | Cuvier 1829 | INPA-ICT 056147 |  |
| Characiformes | Crenuchidae | *Ammocryptocharax* | *elegans* | Weitzman & Kanazawa 1976 | INPA-ICT 056048 |  |
| Characiformes | Crenuchidae | *Characidium* | *aff. longum* | Taphorn, Montaña & Buckup 2006 | INPA-ICT 055219 | S1, S2, S3 |
| Characiformes | Crenuchidae | *Characidium* | *aff. pteroides* | Eigenmann 1909 | INPA-ICT 056002 | S2 |
| Characiformes | Crenuchidae | *Characidium* | *etheostoma* | Cope 1872 | INPA-ICT 056034 | S2 |
| Characiformes | Crenuchidae | *Characidium* | *longum* | Taphorn, Montaña & Buckup 2006 | INPA-ICT 056082 |  |
| Characiformes | Crenuchidae | *Elachocharax* | *pulcher* | Myers 1927 | INPA-ICT 055227 |  |
| Characiformes | Crenuchidae | *Klausewitzia* | *ritae* | Géry 1965 | INPA-ICT 055801 |  |
| Characiformes | Crenuchidae | *Melanocharacidium* | *dispilomma* | Buckup 1993 | INPA-ICT 056021 | S2 |
| Characiformes | Crenuchidae | *Melanocharacidium* | *melanopteron* | Buckup 1993 | INPA-ICT 055443 | S1 |
| Characiformes | Crenuchidae | *Odontocharacidium* | sp. |  | INPA-ICT 055252 |  |
| Characiformes | Ctenoluciidae | *Boulengerella* | *maculata* | (Valenciennes 1850) | INPA-ICT 055263 | S1 |
| Characiformes | Curimatidae | *Curimata* | *roseni* | Vari 1989 | INPA-ICT 055527 | S1 |
| Characiformes | Curimatidae | *Curimatella* | *dorsalis* | (Eigenmann & Eigenmann 1889) | INPA-ICT 055415 | S1 |
| Characiformes | Curimatidae | *Curimatella* | *immaculata* | (Fernández-Yépez 1948) | INPA-ICT 055416 | S1 |
| Characiformes | Curimatidae | *Curimatella* | *meyeri* | (Steindachner 1882) | INPA-ICT 055417 | S1 |
| Characiformes | Curimatidae | *Cyphocharax* | *aff. spiluropsis* | (Eigenmann & Eigenmann 1889) | INPA-ICT 056060 |  |
| Characiformes | Curimatidae | *Cyphocharax* | *spiluropsis* | (Eigenmann & Eigenmann 1889) | INPA-ICT 055741 |  |
| Characiformes | Curimatidae | *Cyphocharax* | sp. |  | INPA-ICT 055224 |  |
| Characiformes | Curimatidae | *Potamorhina* | *pristigaster* | (Steindachner 1876) | INPA-ICT 056146 |  |
| Characiformes | Curimatidae | *Psectrogaster* | *amazonica* | Eigenmann & Eigenmann 1889 | INPA-ICT 055621 |  |
| Characiformes | Curimatidae | *Steindachnerina* | *bimaculata* | (Steindachner 1876) | INPA-ICT 055377 | S1, S3 |
| Characiformes | Cynodontidae | *Cynodon* | *gibbus* | (Spix & Agassiz 1829) | INPA-ICT 055418 | S1, S3 |
| Characiformes | Cynodontidae | *Hydrolycus* | *scomberoides* | (Cuvier 1819) | INPA-ICT 055303 |  |
| Characiformes | Cynodontidae | *Hydrolycus* | *tatauaia* | Toledo-Piza, Menezes & Santos 1999 | INPA-ICT 055426 | S1 |
| Characiformes | Cynodontidae | *Rhaphiodon* | *vulpinus* | Spix & Agassiz 1829 | INPA-ICT 055465 | S1, S3 |
| Cyprinodontiformes | Rivulidae | *Anablepsoides* | *atratus* | (Garman 1895) | INPA-ICT 055245 |  |
| Cyprinodontiformes | Rivulidae | *Anablepsoides* | sp. |  | INPA-ICT 056031 | S2 |
| Cyprinodontiformes | Rivulidae | *Laimosemion* | sp. |  | INPA-ICT 056039 | S2 |
| Siluriformes | Doradidae | *Acanthodoras* | *cataphractus* | (Linnaeus 1758) | INPA-ICT 055726 |  |
| Siluriformes | Doradidae | *Acanthodoras* | *spinosissimus* | (Eigenmann & Eigenmann 1888) | INPA-ICT 055384 | S1 |
| Siluriformes | Doradidae | *Agamyxis* | *pectinifrons* | (Cope 1870) | INPA-ICT 055388 | S1 |
| Siluriformes | Doradidae | *Amblydoras* | *affinis* | (Kner 1855) | INPA-ICT 055184 |  |
| Siluriformes | Doradidae | *Anadoras* | *grypus* | (Cope 1872) | INPA-ICT 055827 |  |
| Siluriformes | Doradidae | *Hemidoras* | *morrisi* | Eigenmann 1925 | INPA-ICT 055422 | S1, S3 |
| Siluriformes | Doradidae | *Hemidoras* | *stenopeltis* | (Kner 1855) | INPA-ICT 055321 | S1 |
| Siluriformes | Doradidae | *Hypodoras* | *forficulatus* | Eigenmann 1925 | INPA-ICT 056261 |  |
| Siluriformes | Doradidae | *Megalodoras* | *uranoscopus* | (Eigenmann & Eigenmann 1888) | INPA-ICT 055491 | S1 |
| Siluriformes | Doradidae | *Tenellus* | *cristinae* | (Sabaj Pérez, Arce H., Sousa & Birindelli 2014) | INPA-ICT 056138 |  |
| Siluriformes | Doradidae | *Nemadoras* | *elongatus* | (Boulenger 1898) | INPA-ICT 055325 | S1 |
| Siluriformes | Doradidae | *Nemadoras* | *humeralis* | (Kner 1855) | INPA-ICT 055449 | S1, S3 |
| Siluriformes | Doradidae | *Hemidoras* | *boulengeri* | Steindachner 1915 | INPA-ICT 055651 |  |
| Siluriformes | Doradidae | *Hemidoras* | *stuebelii* | (Steindachner 1882) | INPA-ICT 055948 |  |
| Siluriformes | Doradidae | *Ossancora* | *punctata* | (Kner 1855) | INPA-ICT 055652 |  |
| Siluriformes | Doradidae | *Oxydoras* | *niger* | (Valenciennes 1821) | INPA-ICT 055371 |  |
| Siluriformes | Doradidae | *Physopyxis* | *ananas* | Sousa & Rapp Py-Daniel 2005 | INPA-ICT 055209 |  |
| Siluriformes | Doradidae | *Physopyxis* | *lyra* | Cope 1872 | INPA-ICT 055809 |  |
| Siluriformes | Doradidae | *Platydoras* | *armatulus* | (Valenciennes 1840) | INPA-ICT 055458 | S1 |
| Siluriformes | Doradidae | *Pterodoras* | *granulosus* | (Valenciennes 1821) | INPA-ICT 055463 | S1 |
| Siluriformes | Doradidae | *Rhinodoras* | *boehlkei* | Glodek, Whitmire & Orcés V. 1976 | INPA-ICT 055874 |  |
| Siluriformes | Doradidae | *Rhynchodoras* | *woodsi* | Glodek 1976 | INPA-ICT 055466 | S1 |
| Siluriformes | Doradidae | *Tenellus* | *ternetzi* | (Eigenmann 1925) | INPA-ICT 055337 |  |
| Siluriformes | Doradidae | *Tenellus* | *trimaculatus* | (Boulenger 1898) | INPA-ICT 055381 |  |
| Siluriformes | Doradidae | *Trachydoras* | *brevis* | (Kner 1853) | INPA-ICT 055582 |  |
| Siluriformes | Doradidae | *Trachydoras* | *nattereri* | (Steindachner 1881) | INPA-ICT 055660 |  |
| Siluriformes | Doradidae | *Trachydoras* | sp. |  | INPA-ICT 055338 | S1, S3 |
| Siluriformes | Doradidae | *Trachydoras* | *steindachneri* | (Perugia 1897) | INPA-ICT 055521 | S1 |
| Gobiiformes | Eleotridae | *Microphilypnus* | *ternetzi* | Myers 1927 | INPA-ICT 055752 |  |
| Cupleiformes | Engraulidae | *Amazonsprattus* | *scintilla* | Roberts 1984 | INPA-ICT 056602 |  |
| Cupleiformes | Engraulidae | *Anchoviella* | *guianensis* | (Eigenmann 1912) | INPA-ICT 055766 |  |
| Cupleiformes | Engraulidae | *Anchoviella* | *jamesi* | (Jordan & Seale 1926) | INPA-ICT 055698 |  |
| Cupleiformes | Engraulidae | *Anchoviella* | sp. |  | INPA-ICT 055391 | S1, S3 |
| Cupleiformes | Engraulidae | *Lycengraulis* | *batesii* | (Günther 1868) | INPA-ICT 056128 |  |
| Characiformes | Erythrinidae | *Erythrinus* | *erythrinus* | (Bloch & Schneider 1801) | INPA-ICT 055200 | S2 |
| Characiformes | Erythrinidae | *Hoplerythrinus* | *unitaeniatus* | (Spix & Agassiz 1829) | INPA-ICT 056180 |  |
| Characiformes | Erythrinidae | *Hoplias* | *malabaricus* | (Bloch 1794) | INPA-ICT 055204 | S1, S2, S3 |
| Characiformes | Gasteropelecidae | *Carnegiella* | *myersi* | Fernández-Yépez 1950 | INPA-ICT 055734 |  |
| Characiformes | Gasteropelecidae | *Carnegiella* | *strigata* | (Günther 1864) | INPA-ICT 055195 | S2 |
| Characiformes | Gasteropelecidae | *Gasteropelecus* | *sternicla* | (Linnaeus 1758) | INPA-ICT 055229 |  |
| Characiformes | Gasteropelecidae | *Thoracocharax* | *stellatus* | (Kner 1858) | INPA-ICT 055288 | S1, S3 |
| Gymnotiformes | Gymnotidae | *Electrophorus* | *varii* | de Santana, Wosiacki, Crampton, Sabaj, Dillman, Mendes-Júnior & Castro e C. 2019 | INPA-ICT 056179 |  |
| Gymnotiformes | Gymnotidae | *Gymnotus* | *coropinae* | Hoedeman 1962 | INPA-ICT 056013 | S2 |
| Gymnotiformes | Gymnotidae | *Gymnotus* | *curupira* | Crampton, Thorsen & Albert 2005 | INPA-ICT 056014 | S2 |
| Gymnotiformes | Gymnotidae | *Gymnotus* | *javari* | Albert, Crampton & Hagedorn 2003 | INPA-ICT 056338 |  |
| Gymnotiformes | Gymnotidae | *Gymnotus* | *ucamara* | Crampton, Lovejoy & Albert 2003 | INPA-ICT 055794 |  |
| Characiformes | Hemiodontidae | *Anodus* | *orinocensis* | (Steindachner 1887) | INPA-ICT 055867 |  |
| Characiformes | Hemiodontidae | *Anodus* | sp. |  | INPA-ICT 056320 |  |
| Characiformes | Hemiodontidae | *Hemiodus* | *microlepis* | Kner 1858 | INPA-ICT 055423 | S1 |
| Characiformes | Hemiodontidae | *Hemiodus* | *unimaculatus* | (Bloch 1794) | INPA-ICT 055156 | S1 |
| Siluriformes | Heptapteridae | *Brachyrhamdia* | *meesi* | Sands & Black 1985 | INPA-ICT 055996 | S2 |
| Siluriformes | Heptapteridae | *Brachyrhamdia* | sp. |  | INPA-ICT 055192 |  |
| Siluriformes | Heptapteridae | *Chasmocranus* | sp. |  | INPA-ICT 056003 | S2 |
| Siluriformes | Heptapteridae | *Gladioglanis* | *conquistador* | Lundberg, Bornbusch & Mago-Leccia 1991 | INPA-ICT 055201 |  |
| Siluriformes | Heptapteridae | *Goeldiella* | *eques* | (Müller & Troschel 1849) | INPA-ICT 055230 |  |
| Siluriformes | Heptapteridae | *Imparfinis* | *stictonotus* | (Fowler 1940) | INPA-ICT 055242 | S1 |
| Siluriformes | Heptapteridae | *Mastiglanis* | sp. |  | INPA-ICT 055324 | S1, S3 |
| Siluriformes | Heptapteridae | *Myoglanis* | *koepckei* | Chang 1999 | INPA-ICT 056024 | S2 |
| Siluriformes | Heptapteridae | *Phenacorhamdia* | *aff. boliviana* | (Pearson 1924) | INPA-ICT 056144 |  |
| Siluriformes | Heptapteridae | *Phenacorhamdia* | sp. |  | INPA-ICT 056268 |  |
| Siluriformes | Heptapteridae | *Pimelodella* | *aff. boliviana* | Eigenmann 1917 | INPA-ICT 055541 | S1 |
| Siluriformes | Heptapteridae | *Pimelodella* | *boliviana* | Eigenmann 1917 | INPA-ICT 055164 | S1, S2, S3 |
| Siluriformes | Heptapteridae | *Pimelodella* | *howesi* | Fowler 1940 | INPA-ICT 055494 | S1 |
| Siluriformes | Heptapteridae | *Pimelodella* | *serrata* | Eigenmann 1917 | INPA-ICT 055165 | S1, S3 |
| Siluriformes | Heptapteridae | *Pimelodella* | sp. |  | INPA-ICT 055455 | S1 |
| Siluriformes | Heptapteridae | *Rhamdia* | *laukidi* | Bleeker 1858 | INPA-ICT 055212 |  |
| Gymnotiformes | Hypopomidae | *Brachyhypopomus* | *batesi* | Crampton, de Santana, Waddell & Lovejoy 2017 | INPA-ICT 056326 |  |
| Gymnotiformes | Hypopomidae | *Brachyhypopomus* | *beebei* | (Schultz 1944) | INPA-ICT 056169 |  |
| Gymnotiformes | Hypopomidae | *Brachyhypopomus* | *brevirostris* | (Steindachner 1868) | INPA-ICT 055189 |  |
| Gymnotiformes | Hypopomidae | *Brachyhypopomus* | *hamiltoni* | Crampton, de Santana, Waddell & Lovejoy 2017 | INPA-ICT 056170 |  |
| Gymnotiformes | Hypopomidae | *Brachyhypopomus* | sp. |  | INPA-ICT 055190 |  |
| Gymnotiformes | Hypopomidae | *Brachyhypopomus* | *sullivani* | Crampton, de Santana, Waddell & Lovejoy 2017 | INPA-ICT 056418 |  |
| Gymnotiformes | Hypopomidae | *Brachyhypopomus* | *walteri* | Sullivan, Zuanon & Cox Fernandes 2013 | INPA-ICT 055191 |  |
| Gymnotiformes | Rhamphichthyidae | *Hypopygus* | *lepturus* | Hoedeman 1962 | INPA-ICT 055241 | S2 |
| Gymnotiformes | Rhamphichthyidae | *Steatogenys* | *elegans* | (Steindachner 1880) | INPA-ICT 055176 | S1 |
| Gymnotiformes | Rhamphichthyidae | *Steatogenys* | sp. |  | INPA-ICT 055331 | S1 |
| Characiformes | Lebiasinidae | *Copeina* | *guttata* | (Steindachner 1876) | INPA-ICT 056004 | S2 |
| Characiformes | Lebiasinidae | *Copella* | *callolepis* | (Regan 1912) | INPA-ICT 055221 |  |
| Characiformes | Lebiasinidae | *Nannostomus* | *digrammus* | (Fowler 1913) | INPA-ICT 055756 |  |
| Characiformes | Lebiasinidae | *Nannostomus* | *eques* | Steindachner 1876 | INPA-ICT 055249 |  |
| Characiformes | Lebiasinidae | *Nannostomus* | *trifasciatus* | Steindachner 1876 | INPA-ICT 055250 |  |
| Characiformes | Lebiasinidae | *Pyrrhulina* | *laeta* | (Cope 1872) | INPA-ICT 055210 |  |
| Characiformes | Lebiasinidae | *Pyrrhulina* | *lugubris* | Eigenmann 1922 | INPA-ICT 055990 |  |
| Characiformes | Lebiasinidae | *Pyrrhulina* | *semifasciata* | Steindachner 1876 | INPA-ICT 055211 |  |
| Siluriformes | Loricariidae | *Ancistrus* | sp. |  | INPA-ICT 055185 | S1, S2 |
| Siluriformes | Loricariidae | *Aphanotorulus* | *emarginatus* | (Valenciennes 1840) | INPA-ICT 055261 | S1, S3 |
| Siluriformes | Loricariidae | *Dekeyseria* | *amazonica* | Rapp Py-Daniel 1985 | INPA-ICT 055419 | S1 |
| Siluriformes | Loricariidae | *Farlowella* | *amazonum* | (Günther 1864) | INPA-ICT 055228 |  |
| Siluriformes | Loricariidae | *Farlowella* | *oxyrryncha* | (Kner 1853) | INPA-ICT 055320 |  |
| Siluriformes | Loricariidae | *Farlowella* | *smithi* | Fowler 1913 | INPA-ICT 056716 |  |
| Siluriformes | Loricariidae | *Hemiodontichthys* | *acipenserinus* | (Kner 1853) | INPA-ICT 055155 |  |
| Siluriformes | Loricariidae | *Hypoptopoma* | *gulare* | Cope 1878 | INPA-ICT 055428 | S1, S3 |
| Siluriformes | Loricariidae | *Hypoptopoma* | *steindachneri* | Boulenger 1895 | INPA-ICT 055270 | S1 |
| Siluriformes | Loricariidae | *Hypoptopoma* | *sternoptychum* | (Schaefer 1996) | INPA-ICT 055323 | S1 |
| Siluriformes | Loricariidae | *Hypoptopoma* | *thoracatum* | Günther 1868 | INPA-ICT 055205 | S1 |
| Siluriformes | Loricariidae | *Hypostomus* | *cf. plecostomus* | (Linnaeus 1758) | INPA-ICT 055206 | S1 |
| Siluriformes | Loricariidae | *Hypostomus* | *pyrineusi* | (Miranda Ribeiro 1920) | INPA-ICT 055433 | S1 |
| Siluriformes | Loricariidae | *Lamontichthys* | *filamentosus* | (LaMonte 1935) | INPA-ICT 055436 | S1 |
| Siluriformes | Loricariidae | *Lasiancistrus* | *aff. schomburgkii* | (Günther 1864) | INPA-ICT 055437 | S1 |
| Siluriformes | Loricariidae | *Limatulichthys* | *griseus* | (Eigenmann 1909) | INPA-ICT 055159 | S1 |
| Siluriformes | Loricariidae | *Loricaria* | *cataphracta* | Linnaeus 1758 | INPA-ICT 055276 | S1 |
| Siluriformes | Loricariidae | *Otocinclus* | aff. *macrospilus* | Eigenmann & Allen 1942 | INPA-ICT 056184 |  |
| Siluriformes | Loricariidae | *Otocinclus* | *macrospilus* | Eigenmann & Allen 1942 | INPA-ICT 056526 |  |
| Siluriformes | Loricariidae | *Oxyropsis* | *wrightiana* | Eigenmann & Eigenmann 1889 | INPA-ICT 055208 |  |
| Siluriformes | Loricariidae | *Panaqolus* | *purusiensis* | (LaMonte 1935) | INPA-ICT 055450 | S1 |
| Siluriformes | Loricariidae | *Panaqolus* | sp. |  | INPA-ICT 057159 | S1 |
| Siluriformes | Loricariidae | *Peckoltia* | *brevis* | (LaMonte 1935) | INPA-ICT 055452 | S1 |
| Siluriformes | Loricariidae | *Peckoltichthys* | *bachi* | (Boulenger 1898) | INPA-ICT 055453 | S1 |
| Siluriformes | Loricariidae | *Pseudacanthicus* | sp. |  | INPA-ICT 055461 | S1 |
| Siluriformes | Loricariidae | *Pseudorinelepis* | *genibarbis* | (Valenciennes 1840) | INPA-ICT 055150 |  |
| Siluriformes | Loricariidae | *Pterygoplichthys* | *gibbiceps* | (Kner 1854) | INPA-ICT 055464 | S1 |
| Siluriformes | Loricariidae | *Pterygoplichthys* | *pardalis* | (Castelnau 1855) | INPA-ICT 056357 |  |
| Siluriformes | Loricariidae | *Rineloricaria* | *lanceolata* | (Günther 1868) | INPA-ICT 055467 | S1 |
| Siluriformes | Loricariidae | *Rineloricaria* | *phoxocephala* | (Eigenmann & Eigenmann 1889) | INPA-ICT 055169 |  |
| Siluriformes | Loricariidae | *Rineloricaria* | sp. |  | INPA-ICT 055656 |  |
| Siluriformes | Loricariidae | *Spatuloricaria* | sp. |  | INPA-ICT 055474 | S1 |
| Siluriformes | Loricariidae | *Sturisoma* | sp. |  | INPA-ICT 055286 |  |
| Osteoglossiformes | Osteoglossidae | *Osteoglossum* | *bicirrhosum* | (Cuvier 1829) | INPA-ICT 056354 |  |
| Characiformes | Parodontidae | *Parodon* | aff. *pongoensis* | (Allen 1942) | INPA-ICT 056729 |  |
| Siluriformes | Pimelodidae | *Brachyplatystoma* | *juruense* | (Boulenger 1898) | INPA-ICT 055401 | S1 |
| Siluriformes | Pimelodidae | *Brachyplatystoma* | *vaillantii* | (Valenciennes 1840) | INPA-ICT 056703 |  |
| Siluriformes | Pimelodidae | *Calophysus* | *macropterus* | (Lichtenstein 1819) | INPA-ICT 055339 | S1 |
| Siluriformes | Pimelodidae | *Cheirocerus* | *eques* | Eigenmann 1917 | INPA-ICT 055151 |  |
| Siluriformes | Pimelodidae | *Cheirocerus* | *goeldii* | (Steindachner 1908) | INPA-ICT 055409 | S1 |
| Siluriformes | Pimelodidae | *Hemisorubim* | *platyrhynchos* | (Valenciennes 1840) | INPA-ICT 055267 |  |
| Siluriformes | Pimelodidae | *Hypophthalmus* | *fimbriatus* | Kner 1858 | INPA-ICT 056688 |  |
| Siluriformes | Pimelodidae | *Hypophthalmus* | *marginatus* | Valenciennes 1840 | INPA-ICT 055157 | S1 |
| Siluriformes | Pimelodidae | *Hypophthalmus* | *oremaculatus* | Nani & Fuster de Plaza 1947 | INPA-ICT 056117 |  |
| Siluriformes | Pimelodidae | *Leiarius* | *marmoratus* | (Gill 1870) | INPA-ICT 055976 |  |
| Siluriformes | Pimelodidae | *Leiarius* | *pictus* | (Müller & Troschel 1849) | INPA-ICT 055438 | S1 |
| Siluriformes | Pimelodidae | *Megalonema* | *amaxanthum* | Lundberg & Dahdul 2008 | INPA-ICT 055510 | S1 |
| Siluriformes | Pimelodidae | *Phractocephalus* | *hemioliopterus* | (Bloch & Schneider 1801) | INPA-ICT 055372 |  |
| Siluriformes | Pimelodidae | *Pimelodina* | *flavipinnis* | Steindachner 1876 | INPA-ICT 055495 | S1 |
| Siluriformes | Pimelodidae | *Pimelodus* | *albofasciatus* | Mees 1974 | INPA-ICT 056027 | S2 |
| Siluriformes | Pimelodidae | *Pimelodus* | *altissimus* | Eigenmann & Pearson 1942 | INPA-ICT 055687 |  |
| Siluriformes | Pimelodidae | *Pimelodus* | *blochii* | Valenciennes 1840 | INPA-ICT 055166 | S1, S3 |
| Siluriformes | Pimelodidae | *Pinirampus* | *pirinampu* | (Spix & Agassiz 1829) | INPA-ICT 055316 |  |
| Siluriformes | Pimelodidae | *Platynematichthys* | *notatus* | (Jardine 1841) | INPA-ICT 056704 |  |
| Siluriformes | Pimelodidae | *Platysilurus* | *mucosus* | (Vaillant 1880) | INPA-ICT 055513 | S1 |
| Siluriformes | Pimelodidae | *Propimelodus* | *caesius* | Parisi, Lundberg & DoNascimiento 2006 | INPA-ICT 055515 | S1 |
| Siluriformes | Pimelodidae | Propimelodus | sp. |  | INPA-ICT 055460 | S1 |
| Siluriformes | Pimelodidae | *Pseudoplatystoma* | *punctifer* | (Castelnau 1855) | INPA-ICT 055622 |  |
| Siluriformes | Pimelodidae | *Pseudoplatystoma* | *tigrinum* | (Valenciennes 1840) | INPA-ICT 055379 |  |
| Siluriformes | Pimelodidae | *Sorubim* | *elongatus* | Littmann, Burr, Schmidt & Isern 2001 | INPA-ICT 055657 |  |
| Siluriformes | Pimelodidae | *Sorubim* | *lima* | (Bloch & Schneider 1801) | INPA-ICT 056274 |  |
| Siluriformes | Pimelodidae | *Sorubim* | *maniradii* | Littmann, Burr & Buitrago-Suarez 2001 | INPA-ICT 055473 | S1 |
| Siluriformes | Pimelodidae | *Sorubimichthys* | *planiceps* | (Spix & Agassiz 1829) | INPA-ICT 055376 |  |
| "Perciformes" | Polycentridae | *Monocirrhus* | *polyacanthus* | Heckel 1840 | INPA-ICT 056352 |  |
| Myliobatiformes | Potamotrygonidae | *Potamotrygon* | *motoro* | (Müller & Henle 1841) | INPA-ICT 055552 | S1 |
| Myliobatiformes | Potamotrygonidae | *Potamotrygon* | *orbignyi* | (Castelnau 1855) | INPA-ICT 055897 |  |
| Myliobatiformes | Potamotrygonidae | *Potamotrygon* | *scobina* | Garman 1913 | INPA-ICT 055553 | S1 |
| Characiformes | Prochilodontidae | *Prochilodus* | *nigricans* | Spix & Agassiz 1829 | INPA-ICT 055168 | S1, S3 |
| Characiformes | Prochilodontidae | *Semaprochilodus* | *insignis* | (Jardine 1841) | INPA-ICT 056029 | S2 |
| Siluriformes | Pseudopimelodidae | *Batrochoglanis* | cf. *raninus* | (Valenciennes 1840) | INPA-ICT 056416 |  |
| Siluriformes | Pseudopimelodidae | *Batrochoglanis* | cf. *villosus* | (Eigenmann 1912) | INPA-ICT 056101 |  |
| Siluriformes | Pseudopimelodidae | *Microglanis* | cf. *poecilus* | Eigenmann 1912 | INPA-ICT 055854 |  |
| Siluriformes | Pseudopimelodidae | *Microglanis* | cf. *secundus* | Mees 1974 | INPA-ICT 056132 |  |
| Gymnotiformes | Rhamphichthyidae | *Gymnorhamphichthys* | *rondoni* | (Miranda Ribeiro 1920) | INPA-ICT 056012 | S2 |
| Gymnotiformes | Rhamphichthyidae | *Rhamphichthys* | *pantherinus* | Castelnau 1855 | INPA-ICT 055578 |  |
| Gymnotiformes | Rhamphichthyidae | *Rhamphichthys* | *rostratus* | (Linnaeus 1766) | INPA-ICT 055175 |  |
| Perciformes | Sciaenidae | *Pachyurus* | *gabrielensis* | Casatti 2001 | INPA-ICT 055163 |  |
| Perciformes | Sciaenidae | *Pachyurus* | *schomburgkii* | Günther 1860 | INPA-ICT 055721 |  |
| Perciformes | Sciaenidae | *Plagioscion* | *squamosissimus* | (Heckel 1840) | INPA-ICT 055328 | S1, S3 |
| Siluriformes | Scoloplacidae | *Scoloplax* | *baskini* | Rocha, de Oliveira & Rapp Py-Daniel 2008 | INPA-ICT 056316 |  |
| Siluriformes | Scoloplacidae | *Scoloplax* | *dicra* | Bailey & Baskin 1976 | INPA-ICT 055213 |  |
| Characiformes | Serrasalmidae | *Colossoma* | *macropomum* | (Cuvier 1816) | INPA-ICT 056330 |  |
| Characiformes | Serrasalmidae | *Metynnis* | *altidorsalis* | Ahl 1923 | INPA-ICT 056130 |  |
| Characiformes | Serrasalmidae | *Metynnis* | *luna* | Cope 1878 | INPA-ICT 055612 |  |
| Characiformes | Serrasalmidae | *Myloplus* | *asterias* | (Müller & Troschel 1844) | INPA-ICT 055149 |  |
| Characiformes | Serrasalmidae | *Myloplus* | *lobatus* | (Valenciennes 1850) | INPA-ICT 056136 |  |
| Characiformes | Serrasalmidae | *Mylossoma* | *aureum* | (Spix & Agassiz 1829) | INPA-ICT 055294 | S1, S3 |
| Characiformes | Serrasalmidae | *Mylossoma* | *duriventre* | (Cuvier 1818) | INPA-ICT 055295 | S1 |
| Characiformes | Serrasalmidae | *Pristobrycon* | *calmoni* | (Steindachner 1908) | INPA-ICT 056356 |  |
| Characiformes | Serrasalmidae | *Pygocentrus* | *nattereri* | Kner 1858 | INPA-ICT 055811 |  |
| Characiformes | Serrasalmidae | *Serrasalmus* | *compressus* | Jégu, Leão & Santos 1991 | INPA-ICT 055470 | S1, S3 |
| Characiformes | Serrasalmidae | *Serrasalmus* | *elongatus* | Kner 1858 | INPA-ICT 056157 |  |
| Characiformes | Serrasalmidae | *Serrasalmus* | *hollandi* | Eigenmann 1915 | INPA-ICT 056272 |  |
| Characiformes | Serrasalmidae | *Serrasalmus* | *rhombeus* | (Linnaeus 1766) | INPA-ICT 055471 | S1 |
| Characiformes | Serrasalmidae | *Serrasalmus* | sp. |  | INPA-ICT 055472 | S1 |
| Characiformes | Serrasalmidae | *Serrasalmus* | *spilopleura* | Kner 1858 | INPA-ICT 056362 |  |
| Gymnotiformes | Sternopygidae | *Distocyclus* | *conirostris* | (Eigenmann & Allen 1942) | INPA-ICT 055300 |  |
| Gymnotiformes | Sternopygidae | *Eigenmannia* | af*f. trilineata* | López & Castello 1966 | INPA-ICT 055225 |  |
| Gymnotiformes | Sternopygidae | *Eigenmannia* | *limbata* | (Schreiner & Miranda Ribeiro 1903) | INPA-ICT 055420 | S1, S3 |
| Gymnotiformes | Sternopygidae | *Eigenmannia* | *macrops* | (Boulenger 1897) | INPA-ICT 055226 | S1, S3 |
| Gymnotiformes | Sternopygidae | *Eigenmannia* | sp. |  | INPA-ICT 055302 |  |
| Gymnotiformes | Sternopygidae | *Rhabdolichops* | aff. *electrogrammus* | Lundberg & Mago-Leccia 1986 | INPA-ICT 056692 |  |
| Gymnotiformes | Sternopygidae | *Rhabdolichops* | *caviceps* | (Fernández-Yépez 1968) | INPA-ICT 055329 | S1 |
| Gymnotiformes | Sternopygidae | *Rhabdolichops* | *eastwardi* | Lundberg & Mago-Leccia 1986 | INPA-ICT 055305 | S1 |
| Gymnotiformes | Sternopygidae | *Rhabdolichops* | *lundbergi* | Correa, Crampton & Albert 2006 | INPA-ICT 055306 | S1, S3 |
| Gymnotiformes | Sternopygidae | *Rhabdolichops* | *navalha* | Correa, Crampton & Albert 2006 | INPA-ICT 055174 | S1 |
| Gymnotiformes | Sternopygidae | *Rhabdolichops* | *troscheli* | (Kaup 1856) | INPA-ICT 055330 | S1 |
| Gymnotiformes | Sternopygidae | *Sternopygus* | *macrurus* | (Bloch & Schneider 1801) | INPA-ICT 055309 | S1 |
| Synbranchiformes | Synbranchidae | *Synbranchus* | sp. |  | INPA-ICT 055815 |  |
| Tetraodontiformes | Tetraodontidae | *Colomesus* | *asellus* | (Müller & Troschel 1849) | INPA-ICT 055152 |  |
| Siluriformes | Trichomycteridae | *Henonemus* | *punctatus* | (Boulenger 1887) | INPA-ICT 055268 | S1 |
| Siluriformes | Trichomycteridae | *Ituglanis* | *amazonicus* | (Steindachner 1882) | INPA-ICT 056346 |  |
| Siluriformes | Trichomycteridae | *Ituglanis* | *parkoi* | (Miranda Ribeiro 1944) | INPA-ICT 056122 |  |
| Siluriformes | Trichomycteridae | *Ochmacanthus* | *reinhardtii* | (Steindachner 1882) | INPA-ICT 055161 |  |
| Siluriformes | Trichomycteridae | *Ochmacanthus* | sp. |  | INPA-ICT 055600 |  |
| Siluriformes | Trichomycteridae | *Paravandellia* | sp. |  | INPA-ICT 055922 |  |
| Siluriformes | Trichomycteridae | *Tridens* | *melanops* | Eigenmann & Eigenmann 1889 | INPA-ICT 056579 |  |
| Siluriformes | Trichomycteridae | *Tridens* | sp. |  | INPA-ICT 055258 |  |
| Siluriformes | Trichomycteridae | *Vandellia* | *cirrhosa* | Valenciennes 1846 | INPA-ICT 055291 |  |
| Characiformes | Triportheidae | *Engraulisoma* | *taeniatum* | Castro 1981 | INPA-ICT 055709 |  |
