## Supplementary material for "The critical role of natural history museums in advancing eDNA for biodiversity studies: a case study with Amazonian fishes": Table S2

| Library | Raw read | Merged | Q filtered | Denoised | Fish | Non-fish |
| --- | --- | --- | --- | --- | --- | --- |
| Sample 1 | 199,323 | 192,517 (96.6%) | 190,348 (95.5%) | 178,079 (89.3%) | 178,011 (100.0%) | 68 (0.0%) |
| Sample 2 | 195,723 | 185,268 (94.7%) | 182,705 (93.4%) | 168,440 (86.1%) | 168,410 (100.0%) | 30 (0.0%) |
| Sample 3 | 213,952 | 206,875 (96.7%) | 204,424 (95.6%) | 191,076 (89.3%) | 190,697 (99.8%) | 379 (0.2%) |
| Sample 4 | 185,255 | 179,218 (96.7%) | 177,224 (95.7%) | 165,841 (89.5%) | 165,771 (100.0%) | 70 (0.0%) |
| Sample 5 | 141,670 | 138,448 (97.7%) | 137,235 (96.9%) | 128,337 (90.6%) | 127,829 (99.6%) | 508 (0.4%) |
| Sample 6 | 135,818 | 131,947 (97.2%) | 130,262 (95.9%) | 122,057 (89.9%) | 120,512 (98.7%) | 1,545 (1.3%) |
| Sample 7 | 159,113 | 144,883 (91.1%) | 142,239 (89.4%) | 130,381 (81.9%) | 129,495 (99.3%) | 886 (0.7%) |
| Sample 8 | 164,230 | 157,926 (96.2%) | 155,317 (94.6%) | 145,128 (88.4%) | 145,108 (100.0%) | 20 (0.0%) |
| Sample 9 | 173,925 | 168,894 (97.1%) | 166,543 (95.8%) | 154,978 (89.1%) | 154,369 (99.6%) | 609 (0.4%) |
| Sample 10 | 162,904 | 155,536 (95.5%) | 152,780 (93.8%) | 140,909 (86.5%) | 140,531 (99.7%) | 378 (0.3%) |
| Sample 11 | 171,247 | 165,316 (96.5%) | 163,021 (95.2%) | 152,176 (88.9%) | 151,138 (99.3%) | 1,038 (0.7%) |
| Total | 1,903,160 | 1,826,828 (96.0) | 1,802,098 (94.7) | 1,677,402 (88.2%) | 1,671,871 (99.6) | 5,531 (0.4%) |
| EB | 101 | 90 (89.1%) | 89 (88.1%) | 54 (53.5%) | 54 (100.0%) | 0 (0.0%) |
| 1B | 80 | 76 (95.0%) | 73 (91.3%) | 49 (61.3%) | 49 (100.0%) | 0 (0.0%) |
| 2B | 13 | 11 (84.6%) | 10 (76.9%) | 0 (0.0%) | 0 (0.0%) | 0 (0.0%) |
| Total | 194 | 177 (89.6%) | 172 (97.2%) | 103 (58.1%) | 103 (58.1%) | 0 (0.0%) |
