## Supplementary material for "The critical role of natural history museums in advancing eDNA for biodiversity studies: a case study with Amazonian fishes": Table S3

| Order | Family | Scientific Name | Ave. Identity | Ave. LOD | Total read | Freq | 273-A001-Amazon_S129 Station 2 | 274-A002-Amazon_S130 Station 2 | 275-A003-Amazon_S131Station 2 | 276-A004-Amazon_S132 Station 2 | 277-A005-Amazon_S133 Station 2 | 278-A006-Amazon_S134 Station 1 | 279-A007-Amazon_S135 Station 1 | 280-A008-Amazon_S136 Station 1 | 281-A009-Amazon_S137 Station 1 | 282-A010-Amazon_S138 Station 1 | 283-A011-Amazon_S139 Station 3 |
| --- | --- | --- | --- | --- | --- | --- | --- | --- | --- | --- | --- | --- | --- | --- | --- | --- | --- |
|  |  |  |  |  |  |  | 177,577 | 162,385 | 190,494 | 165,388 | 127,669 | 120,512 | 129,495 | 145,108 | 149,726 | 140,509 | 149,602 |
|  |  |  |  |  |  |  | 84 | 87 | 75 | 62 | 71 | 33 | 39 | 33 | 47 | 41 | 38 |
| Osteoglossiformes | Arapaimidae | *Arapaima gigas* | 100.0 | HIGH | 7,050 | 4 | 0 | 0 | 0 | 0 | 0 | 1,794 | 3,584 | 1,454 | 0 | 218 | 0 |
| Clupeiformes | Engraulidae | *Anchoviella*_sp_1 | 98.8 | HIGH | 18,117 | 4 | 0 | 0 | 0 | 0 | 0 | 0 | 1,670 | 6,299 | 9,778 | 370 | 0 |
| Clupeiformes | Engraulidae | *Anchoviella*_sp_2 | 95.3 | LOW | 2,970 | 1 | 0 | 0 | 0 | 0 | 0 | 0 | 2,970 | 0 | 0 | 0 | 0 |
| Clupeiformes | Clupeidae | *Pellona* sp. 1 | 88.8 | LOW | 1,251 | 1 | 0 | 0 | 0 | 0 | 0 | 0 | 0 | 0 | 1,251 | 0 | 0 |
| Clupeiformes | Clupeidae | *Pellona* sp. 2 | 90.0 | LOW | 840 | 1 | 0 | 0 | 0 | 0 | 0 | 0 | 0 | 0 | 840 | 0 | 0 |
| Characiformes | Crenuchidae | *Characidium* sp. 1 | 98.8 | HIGH | 69 | 2 | 0 | 26 | 0 | 0 | 43 | 0 | 0 | 0 | 0 | 0 | 0 |
| Characiformes | Crenuchidae | *Characidium* sp. 2 | 85.2 | LOW | 2,685 | 6 | 203 | 93 | 168 | 80 | 153 | 0 | 0 | 0 | 0 | 0 | 1,988 |
| Characiformes | Crenuchidae | *Characidium* sp. 3 | 97.0 | HIGH | 99 | 2 | 42 | 57 | 0 | 0 | 0 | 0 | 0 | 0 | 0 | 0 | 0 |
| Characiformes | Crenuchidae | *Characidium* sp. 4 | 89.8 | LOW | 217 | 2 | 0 | 0 | 125 | 0 | 92 | 0 | 0 | 0 | 0 | 0 | 0 |
| Characiformes | Crenuchidae | *Elachocharax*_sp. 1 | 85.9 | LOW | 201 | 3 | 53 | 0 | 73 | 0 | 75 | 0 | 0 | 0 | 0 | 0 | 0 |
| Characiformes | Erythrinidae | *Hoplerythrinus unitaeniatus* | 99.4 | LOW | 1,332 | 5 | 342 | 272 | 190 | 244 | 284 | 0 | 0 | 0 | 0 | 0 | 0 |
| Characiformes | Erythrinidae | *Hoplias malabaricus* | 99.1 | HIGH | 324 | 2 | 241 | 83 | 0 | 0 | 0 | 0 | 0 | 0 | 0 | 0 | 0 |
| Characiformes | Erythrinidae | *Hoplerythrinus* sp. | 87.9 | LOW | 783 | 5 | 135 | 113 | 306 | 111 | 118 | 0 | 0 | 0 | 0 | 0 | 0 |
| Characiformes | Erythrinidae | *Hoplias* sp. | 97.6 | HIGH | 3,721 | 7 | 610 | 58 | 161 | 77 | 36 | 0 | 1,554 | 0 | 1,225 | 0 | 0 |
| Characiformes | Cynodontidae | *Cynodon meionactis* | 98.8 | HIGH | 1,769 | 1 | 0 | 0 | 0 | 0 | 0 | 0 | 0 | 0 | 0 | 1,769 | 0 |
| Characiformes | Cynodontidae | *Hydrolycus scomberoides* | 100.0 | HIGH | 9,118 | 4 | 0 | 0 | 0 | 0 | 0 | 0 | 127 | 4,162 | 4,157 | 672 | 0 |
| Characiformes | Cynodontidae | *Hydrolycus* sp. 1 | 87.7 | LOW | 807 | 3 | 0 | 0 | 0 | 0 | 0 | 770 | 33 | 0 | 4 | 0 | 0 |
| Characiformes | Cynodontidae | *Hydrolycus* sp. 2 | 88.9 | LOW | 147 | 1 | 0 | 147 | 0 | 0 | 0 | 0 | 0 | 0 | 0 | 0 | 0 |
| Characiformes | Serrasalmidae | *Serrasalmus eigenmanni* | 100.0 | HIGH | 1,222 | 2 | 37 | 0 | 0 | 0 | 0 | 0 | 0 | 0 | 0 | 0 | 1,185 |
| Characiformes | Serrasalmidae | *Myleus* sp. | 90.4 | LOW | 949 | 1 | 0 | 0 | 0 | 0 | 0 | 0 | 0 | 0 | 949 | 0 | 0 |
| Characiformes | Hemiodontidae | *Anodus* sp. | 100.0 | LOW | 4,483 | 3 | 0 | 0 | 0 | 0 | 0 | 489 | 3,152 | 0 | 842 | 0 | 0 |
| Characiformes | Anostomidae | *Abramites hypselonotus* | 100.0 | HIGH | 1,304 | 1 | 0 | 0 | 0 | 0 | 0 | 0 | 0 | 0 | 0 | 0 | 1,304 |
| Characiformes | Anostomidae | *Leporinus apollo* | 98.8 | HIGH | 846 | 3 | 0 | 69 | 0 | 0 | 76 | 0 | 0 | 0 | 0 | 701 | 0 |
| Characiformes | Anostomidae | *Leporinus fasciatus* | 100.0 | LOW | 31 | 1 | 31 | 0 | 0 | 0 | 0 | 0 | 0 | 0 | 0 | 0 | 0 |
| Characiformes | Anostomidae | *Anostomus* sp. | 92.4 | LOW | 39,567 | 5 | 9,832 | 6,784 | 10,675 | 6,555 | 5,721 | 0 | 0 | 0 | 0 | 0 | 0 |
| Characiformes | Anostomidae | *Leporinus* sp. 1 | 92.9 | LOW | 350 | 1 | 0 | 0 | 0 | 0 | 0 | 0 | 0 | 0 | 350 | 0 | 0 |
| Characiformes | Anostomidae | *Leporinus* sp. 2 | 92.4 | LOW | 69 | 1 | 0 | 0 | 0 | 0 | 69 | 0 | 0 | 0 | 0 | 0 | 0 |
| Characiformes | Anostomidae | *Leporinus* sp. 3 | 93.6 | LOW | 30 | 1 | 0 | 30 | 0 | 0 | 0 | 0 | 0 | 0 | 0 | 0 | 0 |
| Characiformes | Anostomidae | *Scheizodon knerii* | 100.0 | LOW | 46,775 | 6 | 0 | 0 | 0 | 0 | 0 | 16,370 | 5,944 | 5,951 | 10,218 | 7,017 | 1,275 |
| Characiformes | Anostomidae | *Leporinus* sp. 4 | 98.2 | MODERATE | 173,589 | 6 | 35,887 | 35,144 | 40,068 | 32,987 | 27,560 | 0 | 0 | 1,943 | 0 | 0 | 0 |
| Characiformes | Anostomidae | *Leporinus* sp. 5 | 93.4 | LOW | 24,887 | 9 | 1,874 | 2,090 | 957 | 1,476 | 1,506 | 6,893 | 0 | 5,093 | 1,365 | 3,633 | 0 |
| Characiformes | Anostomidae | *Leporinus* sp. 6 | 92.8 | LOW | 6,498 | 7 | 171 | 373 | 319 | 318 | 56 | 2,265 | 2,996 | 0 | 0 | 0 | 0 |
| Characiformes | Anostomidae | *Leporinus* sp. 7 | 93.5 | LOW | 4,173 | 5 | 1,007 | 541 | 1,273 | 933 | 419 | 0 | 0 | 0 | 0 | 0 | 0 |
| Characiformes | Anostomidae | *Leporinus* sp. 8 | 92.9 | LOW | 665 | 4 | 129 | 114 | 179 | 243 | 0 | 0 | 0 | 0 | 0 | 0 | 0 |
| Characiformes | Anostomidae | *Leporinus* sp. 9 | 93.5 | LOW | 371 | 4 | 73 | 55 | 127 | 116 | 0 | 0 | 0 | 0 | 0 | 0 | 0 |
| Characiformes | Anostomidae | *Leporinus* sp. 10 | 92.4 | LOW | 91 | 2 | 0 | 0 | 12 | 0 | 79 | 0 | 0 | 0 | 0 | 0 | 0 |
| Characiformes | Anostomidae | *Leporinus* sp. 11 | 90.0 | LOW | 2,755 | 2 | 0 | 0 | 0 | 0 | 0 | 0 | 0 | 0 | 994 | 0 | 1,761 |
| Characiformes | Anostomidae | *Megaleporinus* sp. | 91.7 | LOW | 959 | 1 | 0 | 0 | 0 | 0 | 0 | 0 | 0 | 0 | 0 | 959 | 0 |
| Characiformes | Chilodontidae | *Caenotropus labyrinthicus* | 99.4 | HIGH | 14 | 1 | 14 | 0 | 0 | 0 | 0 | 0 | 0 | 0 | 0 | 0 | 0 |
| Characiformes | Curimatidae | *Potamorhina* sp. | 98.7 | LOW | 18,470 | 3 | 0 | 0 | 0 | 0 | 0 | 0 | 9,701 | 0 | 7,405 | 1,364 | 0 |
| Characiformes | Curimatidae | *Cyphocharax* sp. | 95.9 | LOW | 845 | 1 | 0 | 0 | 0 | 0 | 0 | 0 | 0 | 0 | 0 | 845 | 0 |
| Characiformes | Prochilodontidae | *Prochilodus harttii* | 100.0 | LOW | 90,591 | 6 | 0 | 0 | 0 | 0 | 0 | 15,687 | 20,841 | 9,164 | 42,809 | 1,024 | 1,066 |
| Characiformes | Prochilodontidae | *Semaprochilodus* sp. 1 | 100.0 | HIGH | 15,460 | 5 | 2,425 | 2,635 | 5,186 | 3,153 | 2,061 | 0 | 0 | 0 | 0 | 0 | 0 |
| Characiformes | Prochilodontidae | *Chilodus* sp. | 94.0 | MODERATE | 37 | 2 | 0 | 11 | 26 | 0 | 0 | 0 | 0 | 0 | 0 | 0 | 0 |
| Characiformes | Prochilodontidae | *Prochilodus* sp. | 85.9 | LOW | 46,510 | 3 | 0 | 0 | 0 | 0 | 0 | 2,907 | 0 | 0 | 0 | 13 | 43,590 |
| Characiformes | Prochilodontidae | *Semaprochilodus* sp. 2 | 84.1 | LOW | 272 | 2 | 0 | 23 | 249 | 0 | 0 | 0 | 0 | 0 | 0 | 0 | 0 |
| Characiformes | Prochilodontidae | *Semaprochilodus* sp. 3 | 84.7 | LOW | 31 | 2 | 4 | 0 | 27 | 0 | 0 | 0 | 0 | 0 | 0 | 0 | 0 |
| Characiformes | Prochilodontidae | *Semaprochilodus* sp. 4 | 91.6 | LOW | 1,694 | 2 | 0 | 0 | 0 | 0 | 0 | 0 | 1,179 | 515 | 0 | 0 | 0 |
| Characiformes | Prochilodontidae | *Semaprochilodus* sp. 5 | 83.5 | LOW | 179 | 2 | 0 | 58 | 0 | 121 | 0 | 0 | 0 | 0 | 0 | 0 | 0 |
| Characiformes | Acestrorhynchidae | *Acestrorhynchus falcatus* | 99.4 | HIGH | 11,546 | 5 | 1,189 | 3,075 | 2,408 | 3,092 | 1,782 | 0 | 0 | 0 | 0 | 0 | 0 |
| Characiformes | Acestrorhynchidae | *Acestrorhynchus* sp. 1 | 91.7 | MODERATE | 833 | 1 | 0 | 0 | 0 | 0 | 0 | 833 | 0 | 0 | 0 | 0 | 0 |
| Characiformes | Acestrorhynchidae | *Acestrorhynchus* sp. 2 | 84.0 | LOW | 7 | 1 | 7 | 0 | 0 | 0 | 0 | 0 | 0 | 0 | 0 | 0 | 0 |
| Characiformes | Chalceidae | *Chalceus erythrurus* | 100.0 | HIGH | 8,741 | 5 | 0 | 0 | 90 | 0 | 44 | 0 | 3,974 | 1,525 | 0 | 3,108 | 0 |
| Characiformes | Chalceidae | *Chalceus macrolepidotus* | 100.0 | HIGH | 4,158 | 2 | 0 | 0 | 0 | 0 | 0 | 0 | 0 | 0 | 1,037 | 3,121 | 0 |
| Characiformes | Characidae | *Charax pauciradiatus* | 99.4 | HIGH | 10,186 | 5 | 2,505 | 1,852 | 1,715 | 2,334 | 1,780 | 0 | 0 | 0 | 0 | 0 | 0 |
| Characiformes | Characidae | *Charax* sp. 1 | 99.4 | HIGH | 32 | 1 | 0 | 0 | 0 | 0 | 32 | 0 | 0 | 0 | 0 | 0 | 0 |
| Characiformes | Characidae | *Ctenobrycon hauxwellianus* | 100.0 | HIGH | 491 | 3 | 73 | 85 | 0 | 0 | 333 | 0 | 0 | 0 | 0 | 0 | 0 |
| Characiformes | Characidae | *Jupiaba ocellata* | 98.8 | HIGH | 1,150 | 4 | 0 | 331 | 341 | 270 | 208 | 0 | 0 | 0 | 0 | 0 | 0 |
| Characiformes | Characidae | *Moenkhausia_lepidura* | 99.4 | HIGH | 782 | 3 | 64 | 661 | 0 | 57 | 0 | 0 | 0 | 0 | 0 | 0 | 0 |
| Characiformes | Serrasalmidae | *Myloplus rubripinnis* | 99.3 | HIGH | 232 | 4 | 57 | 33 | 114 | 0 | 28 | 0 | 0 | 0 | 0 | 0 | 0 |
| Characiformes | Serrasalmidae | *Pygocentrus nattereri* | 99.9 | HIGH | 19,766 | 9 | 344 | 453 | 329 | 414 | 54 | 5,256 | 7,461 | 0 | 1,539 | 3,916 | 0 |
| Characiformes | Characidae | *Astyanax* sp. 1 | 97.4 | LOW | 4,347 | 5 | 1,061 | 1,208 | 1,333 | 724 | 21 | 0 | 0 | 0 | 0 | 0 | 0 |
| Characiformes | Characidae | *Astyanax* sp. 2 | 97.0 | MODERATE | 4,542 | 5 | 1,206 | 978 | 1,130 | 667 | 561 | 0 | 0 | 0 | 0 | 0 | 0 |
| Characiformes | Characidae | *Astyanax* sp. 3 | 88.6 | LOW | 3,172 | 5 | 746 | 311 | 1,181 | 204 | 730 | 0 | 0 | 0 | 0 | 0 | 0 |
| Characiformes | Characidae | *Astyanax* sp. 4 | 96.4 | LOW | 23 | 1 | 23 | 0 | 0 | 0 | 0 | 0 | 0 | 0 | 0 | 0 | 0 |
| Characiformes | Characidae | *Astyanax* sp. 5 | 95.8 | LOW | 14 | 1 | 14 | 0 | 0 | 0 | 0 | 0 | 0 | 0 | 0 | 0 | 0 |
| Characiformes | Bryconidae | *Brycon* sp. 1 | 96.4 | HIGH | 228,753 | 5 | 61,363 | 43,361 | 46,231 | 43,837 | 33,961 | 0 | 0 | 0 | 0 | 0 | 0 |
| Characiformes | Bryconidae | *Brycon* sp. 2 | 89.2 | LOW | 158,513 | 6 | 28,812 | 30,693 | 36,614 | 36,066 | 26,324 | 0 | 0 | 4 | 0 | 0 | 0 |
| Characiformes | Bryconidae | *Brycon* sp. 3 | 87.0 | LOW | 54,689 | 5 | 9,467 | 10,637 | 12,926 | 12,161 | 9,498 | 0 | 0 | 0 | 0 | 0 | 0 |
| Characiformes | Bryconidae | *Brycon* sp. 4 | 96.4 | HIGH | 2,095 | 5 | 218 | 406 | 650 | 610 | 211 | 0 | 0 | 0 | 0 | 0 | 0 |
| Characiformes | Bryconidae | *Brycon* sp. 5 | 93.0 | LOW | 1,451 | 1 | 0 | 0 | 0 | 0 | 0 | 0 | 0 | 1,451 | 0 | 0 | 0 |
| Characiformes | Iguanodectidae | *Bryconops* sp. 1 | 96.0 | LOW | 5,379 | 5 | 1,565 | 923 | 774 | 1,097 | 1,020 | 0 | 0 | 0 | 0 | 0 | 0 |
| Characiformes | Iguanodectidae | *Bryconops* sp. 2 | 96.5 | LOW | 200 | 5 | 53 | 9 | 30 | 23 | 85 | 0 | 0 | 0 | 0 | 0 | 0 |
| Characiformes | Characidae | *Charax* sp. 2 | 95.3 | LOW | 9,541 | 3 | 0 | 28 | 0 | 0 | 0 | 3,619 | 5,894 | 0 | 0 | 0 | 0 |
| Characiformes | Characidae | *Charax* sp. 3 | 89.2 | LOW | 5,298 | 2 | 0 | 0 | 0 | 0 | 0 | 5,030 | 0 | 0 | 0 | 0 | 268 |
| Characiformes | Characidae | *Charax* sp. 4 | 95.9 | LOW | 142 | 2 | 54 | 0 | 0 | 88 | 0 | 0 | 0 | 0 | 0 | 0 | 0 |
| Characiformes | Characidae | *Charax* sp. 5 | 98.2 | HIGH | 38 | 1 | 0 | 0 | 0 | 0 | 38 | 0 | 0 | 0 | 0 | 0 | 0 |
| Characiformes | Serrasalmidae | *Colossoma* sp. 1 | 93.6 | LOW | 224,686 | 6 | 4 | 0 | 0 | 0 | 0 | 45,419 | 21,671 | 75,422 | 35,247 | 46,923 | 0 |
| Characiformes | Serrasalmidae | *Colossoma* sp. 2 | 89.3 | LOW | 2,473 | 2 | 0 | 0 | 84 | 0 | 0 | 2,389 | 0 | 0 | 0 | 0 | 0 |
| Characiformes | Characidae | *Corynopoma* sp. | 92.4 | LOW | 1,580 | 2 | 0 | 57 | 0 | 0 | 0 | 0 | 1,523 | 0 | 0 | 0 | 0 |
| Characiformes | Characidae | *Cyanocharax* sp. 1 | 91.9 | LOW | 41 | 1 | 41 | 0 | 0 | 0 | 0 | 0 | 0 | 0 | 0 | 0 | 0 |
| Characiformes | Characidae | *Cyanocharax* sp. 2 | 90.8 | LOW | 2,606 | 2 | 0 | 0 | 0 | 0 | 0 | 0 | 0 | 1,245 | 1,361 | 0 | 0 |
| Characiformes | Characidae | *Hasemania* sp. | 85.8 | LOW | 39 | 3 | 11 | 17 | 0 | 11 | 0 | 0 | 0 | 0 | 0 | 0 | 0 |
| Characiformes | Characidae | *Hemigrammus* sp. 1 | 91.7 | LOW | 212 | 3 | 106 | 81 | 0 | 0 | 25 | 0 | 0 | 0 | 0 | 0 | 0 |
| Characiformes | Characidae | *Hyphessobrycon* sp. 1 | 92.7 | MODERATE | 17 | 1 | 0 | 0 | 17 | 0 | 0 | 0 | 0 | 0 | 0 | 0 | 0 |
| Characiformes | Characidae | *Hyphessobrycon* sp. 2 | 86.6 | LOW | 264 | 4 | 29 | 88 | 95 | 0 | 52 | 0 | 0 | 0 | 0 | 0 | 0 |
| Characiformes | Iguanodectidae | *Iguanodectes* sp. | 94.2 | LOW | 145 | 3 | 15 | 71 | 0 | 59 | 0 | 0 | 0 | 0 | 0 | 0 | 0 |
| Characiformes | Characidae | *Jupiaba* sp. 1 | 82.1 | HIGH | 400 | 1 | 0 | 0 | 0 | 0 | 0 | 400 | 0 | 0 | 0 | 0 | 0 |
| Characiformes | Characidae | *Jupiaba* sp. 2 | 94.6 | LOW | 3,339 | 7 | 430 | 329 | 575 | 341 | 153 | 0 | 0 | 0 | 677 | 834 | 0 |
| Characiformes | Characidae | *Moenkhausia* sp. 1 | 97.6 | HIGH | 22 | 1 | 0 | 0 | 0 | 0 | 22 | 0 | 0 | 0 | 0 | 0 | 0 |
| Characiformes | Characidae | *Moenkhausia* sp. 2 | 93.6 | LOW | 684 | 5 | 106 | 97 | 154 | 69 | 258 | 0 | 0 | 0 | 0 | 0 | 0 |
| Characiformes | Characidae | *Moenkhausia* sp. 3 | 97.1 | MODERATE | 5 | 1 | 0 | 0 | 0 | 0 | 0 | 5 | 0 | 0 | 0 | 0 | 0 |
| Characiformes | Characidae | *Moenkhausia* sp. 4 | 91.1 | MODERATE | 688 | 5 | 101 | 309 | 34 | 128 | 116 | 0 | 0 | 0 | 0 | 0 | 0 |
| Characiformes | Characidae | *Moenkhausia* sp. 5 | 89.9 | MODERATE | 14 | 1 | 0 | 0 | 0 | 14 | 0 | 0 | 0 | 0 | 0 | 0 | 0 |
| Characiformes | Characidae | *Paracheirodon* sp. | 88.1 | LOW | 89 | 1 | 0 | 0 | 89 | 0 | 0 | 0 | 0 | 0 | 0 | 0 | 0 |
| Characiformes | Characidae | *Poptella* sp. | 95.2 | LOW | 4,950 | 5 | 1,141 | 1,099 | 1,102 | 1,001 | 607 | 0 | 0 | 0 | 0 | 0 | 0 |
| Characiformes | Serrasalmidae | *Serrasalmus* sp. | 98.2 | LOW | 3,335 | 1 | 0 | 0 | 0 | 0 | 0 | 0 | 3,335 | 0 | 0 | 0 | 0 |
| Characiformes | Iguanodectidae | *Bryconops affinis* | 100.0 | HIGH | 9,344 | 5 | 3,039 | 1,399 | 1,651 | 1,946 | 1,309 | 0 | 0 | 0 | 0 | 0 | 0 |
| Characiformes | Characidae | *Tetragonopterus* sp. | 96.5 | MODERATE | 3,300 | 6 | 1,716 | 127 | 242 | 335 | 223 | 0 | 0 | 0 | 0 | 657 | 0 |
| Characiformes | Characidae | *Tyttocharax* sp. | 94.8 | LOW | 1,572 | 1 | 0 | 0 | 0 | 0 | 0 | 0 | 0 | 0 | 0 | 1,572 | 0 |
| Characiformes | Triportheidae | *Triportheus* sp. 1 | 99.8 | HIGH | 232 | 3 | 0 | 18 | 69 | 0 | 0 | 0 | 0 | 0 | 0 | 0 | 145 |
| Characiformes | Gasteropelecidae | *Thoracocharax* sp. | 93.9 | MODERATE | 1,275 | 1 | 0 | 0 | 0 | 0 | 0 | 0 | 0 | 0 | 1,275 | 0 | 0 |
| Characiformes | Crenuchidae | *Elachocharax*_sp. 2 | 84.3 | LOW | 84 | 3 | 12 | 28 | 0 | 0 | 44 | 0 | 0 | 0 | 0 | 0 | 0 |
| Characiformes | Triportheidae | *Triportheus* sp. 2 | 95.7 | MODERATE | 5,745 | 3 | 0 | 0 | 0 | 0 | 0 | 0 | 5,051 | 0 | 0 | 194 | 500 |
| Characiformes | Triportheidae | *Triportheus* sp. 3 | 91.3 | LOW | 6,537 | 3 | 0 | 0 | 0 | 0 | 0 | 576 | 0 | 4,748 | 0 | 1,213 | 0 |
| Siluriformes | Cetopsidae | *Helogenes* sp. | 90.7 | LOW | 24 | 1 | 0 | 0 | 0 | 0 | 24 | 0 | 0 | 0 | 0 | 0 | 0 |
| Siluriformes | Callichthyidae | *Corydoras* sp. 1 | 95.8 | LOW | 152 | 4 | 57 | 47 | 44 | 4 | 0 | 0 | 0 | 0 | 0 | 0 | 0 |
| Siluriformes | Callichthyidae | *Corydoras* sp. 2 | 98.2 | LOW | 13 | 2 | 0 | 5 | 8 | 0 | 0 | 0 | 0 | 0 | 0 | 0 | 0 |
| Siluriformes | Loricariidae | *Dekeyseria amazonica* | 100.0 | HIGH | 86 | 1 | 0 | 0 | 0 | 0 | 0 | 0 | 0 | 0 | 86 | 0 | 0 |
| Siluriformes | Loricariidae | *Hypostomus plecostomus* | 98.8 | LOW | 113 | 1 | 0 | 0 | 0 | 0 | 0 | 0 | 113 | 0 | 0 | 0 | 0 |
| Siluriformes | Loricariidae | *Lasiancistrus saetiger* | 98.8 | LOW | 1,837 | 1 | 0 | 0 | 0 | 0 | 0 | 0 | 0 | 1,837 | 0 | 0 | 0 |
| Siluriformes | Loricariidae | *Ancistrus* sp. | 97.0 | LOW | 938 | 5 | 246 | 132 | 235 | 166 | 159 | 0 | 0 | 0 | 0 | 0 | 0 |
| Siluriformes | Loricariidae | *Hypoptopoma* sp. 1 | 87.7 | LOW | 116 | 1 | 0 | 0 | 0 | 0 | 0 | 0 | 116 | 0 | 0 | 0 | 0 |
| Siluriformes | Loricariidae | *Hypoptopoma* sp. 2 | 98.1 | MODERATE | 314 | 3 | 0 | 0 | 0 | 0 | 0 | 0 | 63 | 14 | 237 | 0 | 0 |
| Siluriformes | Loricariidae | *Hypoptopoma* sp. 3 | 91.9 | LOW | 1,139 | 1 | 0 | 0 | 0 | 0 | 0 | 0 | 0 | 1,139 | 0 | 0 | 0 |
| Siluriformes | Loricariidae | *Hypostomus* sp. 1 | 98.2 | LOW | 190 | 1 | 0 | 0 | 0 | 0 | 0 | 0 | 0 | 0 | 190 | 0 | 0 |
| Siluriformes | Loricariidae | *Hypostomus* sp. 2 | 98.2 | LOW | 111 | 1 | 0 | 0 | 0 | 0 | 0 | 0 | 0 | 111 | 0 | 0 | 0 |
| Siluriformes | Loricariidae | *Pseudacanthicus* sp. | 97.0 | HIGH | 746 | 1 | 0 | 0 | 0 | 0 | 0 | 0 | 0 | 0 | 746 | 0 | 0 |
| Siluriformes | Loricariidae | *Spatuloricaria* sp. | 98.3 | LOW | 482 | 1 | 0 | 0 | 0 | 0 | 0 | 0 | 482 | 0 | 0 | 0 | 0 |
| Siluriformes | Doradidae | *Hemidoras morrisi* | 99.4 | LOW | 14 | 1 | 0 | 0 | 0 | 0 | 0 | 0 | 0 | 0 | 0 | 14 | 0 |
| Siluriformes | Doradidae | *Liosomadoras morrowi* | 99.4 | HIGH | 25 | 1 | 0 | 0 | 0 | 0 | 0 | 0 | 0 | 0 | 25 | 0 | 0 |
| Siluriformes | Doradidae | *Megalodoras uranoscopus* | 100.0 | HIGH | 9 | 1 | 0 | 0 | 0 | 0 | 0 | 0 | 9 | 0 | 0 | 0 | 0 |
| Siluriformes | Doradidae | *Platydoras costatus* | 100.0 | HIGH | 40 | 2 | 0 | 0 | 0 | 0 | 0 | 0 | 0 | 0 | 10 | 30 | 0 |
| Siluriformes | Doradidae | *Pterodoras granulosus* | 98.8 | HIGH | 94 | 1 | 0 | 0 | 0 | 0 | 0 | 0 | 0 | 0 | 0 | 94 | 0 |
| Siluriformes | Doradidae | *Trachydoras nattereri* | 99.4 | HIGH | 34 | 1 | 0 | 0 | 0 | 0 | 0 | 0 | 0 | 0 | 0 | 0 | 34 |
| Siluriformes | Doradidae | *Nemadoras* sp. | 96.4 | MODERATE | 346 | 1 | 0 | 0 | 0 | 0 | 0 | 0 | 0 | 0 | 0 | 0 | 346 |
| Siluriformes | Doradidae | *Opsodoras* sp. | 96.4 | LOW | 19 | 1 | 0 | 0 | 0 | 0 | 0 | 0 | 0 | 0 | 0 | 0 | 19 |
| Siluriformes | Auchenipteridae | *Ageneiosus* sp. | 95.3 | LOW | 40 | 1 | 0 | 0 | 0 | 0 | 0 | 0 | 40 | 0 | 0 | 0 | 0 |
| Siluriformes | Auchenipteridae | *Auchenipterus* sp. | 97.1 | HIGH | 13 | 1 | 0 | 0 | 0 | 0 | 0 | 0 | 13 | 0 | 0 | 0 | 0 |
| Siluriformes | Auchenipteridae | *Centromochlus* sp. | 89.5 | LOW | 835 | 1 | 0 | 0 | 0 | 0 | 0 | 0 | 835 | 0 | 0 | 0 | 0 |
| Siluriformes | Heptapteridae | *Chasmocranus* sp. | 90.4 | MODERATE | 132 | 4 | 87 | 0 | 4 | 30 | 11 | 0 | 0 | 0 | 0 | 0 | 0 |
| Siluriformes | Heptapteridae | *Imparfinis* sp. | 81.5 | LOW | 26 | 1 | 0 | 0 | 0 | 0 | 26 | 0 | 0 | 0 | 0 | 0 | 0 |
| Siluriformes | Heptapteridae | *Pimelodella* sp. 1 | 98.3 | HIGH | 427 | 1 | 0 | 0 | 0 | 0 | 0 | 0 | 0 | 0 | 427 | 0 | 0 |
| Siluriformes | Heptapteridae | *Pimelodella* sp. 2 | 97.7 | HIGH | 35 | 1 | 0 | 0 | 0 | 0 | 0 | 0 | 0 | 0 | 0 | 0 | 35 |
| Siluriformes | Heptapteridae | *Rhamdia* sp. | 97.7 | LOW | 22 | 1 | 22 | 0 | 0 | 0 | 0 | 0 | 0 | 0 | 0 | 0 | 0 |
| Siluriformes | Pimelodidae | *Hypophthalmus edentatus* | 98.8 | HIGH | 472 | 2 | 0 | 0 | 0 | 0 | 0 | 0 | 76 | 0 | 396 | 0 | 0 |
| Siluriformes | Heptapteridae | *Pimelodella cristata* | 98.9 | HIGH | 1,359 | 7 | 220 | 257 | 315 | 156 | 180 | 0 | 0 | 0 | 217 | 0 | 14 |
| Siluriformes | Pimelodidae | *Pseudoplatystoma eticulatum* | 99.1 | MODERATE | 2,064 | 6 | 0 | 0 | 0 | 0 | 0 | 399 | 728 | 272 | 434 | 57 | 174 |
| Siluriformes | Pimelodidae | *Hypophthalmus* sp. | 97.1 | MODERATE | 102 | 1 | 0 | 0 | 0 | 0 | 0 | 0 | 0 | 0 | 0 | 0 | 102 |
| Siluriformes | Pimelodidae | *Pimelodus* sp. 1 | 97.2 | LOW | 739 | 5 | 0 | 0 | 0 | 0 | 0 | 22 | 77 | 17 | 238 | 385 | 0 |
| Siluriformes | Pimelodidae | *Pimelodus* sp. 2 | 98.3 | LOW | 619 | 4 | 0 | 0 | 0 | 0 | 0 | 0 | 40 | 40 | 356 | 0 | 183 |
| Siluriformes | Pimelodidae | *Pimelodus* sp. 3 | 97.7 | LOW | 58 | 4 | 10 | 22 | 22 | 0 | 4 | 0 | 0 | 0 | 0 | 0 | 0 |
| Siluriformes | Pimelodidae | *Pinirampus pinirampu* | 99.4 | LOW | 4 | 1 | 0 | 0 | 0 | 0 | 0 | 0 | 4 | 0 | 0 | 0 | 0 |
| Siluriformes | Pimelodidae | *Pimelodus* sp. 4 | 97.8 | MODERATE | 2,522 | 5 | 0 | 0 | 0 | 0 | 0 | 509 | 761 | 111 | 516 | 625 | 0 |
| Siluriformes | Pimelodidae | *Pimelodus* sp. 5 | 94.8 | LOW | 140 | 1 | 0 | 0 | 0 | 0 | 0 | 140 | 0 | 0 | 0 | 0 | 0 |
| Siluriformes | Pimelodidae | *Pimelodus* sp. 6 | 92.5 | LOW | 7 | 1 | 0 | 0 | 0 | 0 | 0 | 0 | 0 | 0 | 0 | 0 | 7 |
| Siluriformes | Pimelodidae | *Pimelodus* sp. 7 | 98.3 | HIGH | 10 | 2 | 6 | 0 | 4 | 0 | 0 | 0 | 0 | 0 | 0 | 0 | 0 |
| Siluriformes | Pimelodidae | *Pimelodus* sp. 8 | 97.7 | LOW | 14 | 1 | 0 | 0 | 0 | 0 | 0 | 14 | 0 | 0 | 0 | 0 | 0 |
| Siluriformes | Pimelodidae | *Pimelodus* sp. 9 | 95.4 | LOW | 4 | 1 | 0 | 0 | 0 | 0 | 0 | 0 | 0 | 0 | 0 | 0 | 4 |
| Siluriformes | Pimelodidae | *Pseudoplatystoma tigrinum* | 100.0 | LOW | 183 | 2 | 0 | 0 | 0 | 0 | 0 | 158 | 0 | 0 | 0 | 25 | 0 |
| Siluriformes | Pimelodidae | *Hemisorubim platyrhynchos* | 100.0 | LOW | 110 | 1 | 0 | 0 | 0 | 0 | 0 | 0 | 0 | 110 | 0 | 0 | 0 |
| Siluriformes | Pimelodidae | *Brachyplatystoma vaillantii* | 100.0 | LOW | 25 | 1 | 0 | 0 | 0 | 0 | 0 | 25 | 0 | 0 | 0 | 0 | 0 |
| Siluriformes | Pimelodidae | *Sorubimichthys planiceps* | 99.4 | LOW | 431 | 1 | 0 | 0 | 0 | 0 | 0 | 0 | 0 | 0 | 431 | 0 | 0 |
| Siluriformes | Pimelodidae | *Pseudoplatystoma* sp. | 95.4 | LOW | 39 | 1 | 0 | 0 | 0 | 0 | 0 | 0 | 0 | 0 | 0 | 39 | 0 |
| Siluriformes | Pimelodidae | *Sorubim elongatus* | 100.0 | LOW | 282 | 1 | 0 | 0 | 0 | 0 | 0 | 0 | 0 | 0 | 282 | 0 | 0 |
| Siluriformes | Pimelodidae | *Sorubim* sp. | 97.1 | LOW | 338 | 2 | 0 | 0 | 0 | 0 | 0 | 0 | 0 | 0 | 190 | 0 | 148 |
| Siluriformes | Pimelodidae | *Phractocephalus hemioliopterus* | 100.0 | LOW | 316 | 4 | 0 | 0 | 0 | 0 | 0 | 26 | 0 | 58 | 16 | 216 | 0 |
| Siluriformes | Pimelodidae | *Zungaro* sp. | 86.3 | LOW | 39 | 2 | 4 | 0 | 0 | 35 | 0 | 0 | 0 | 0 | 0 | 0 | 0 |
| Siluriformes | Pimelodidae | *Zungaro jahu* | 100.0 | HIGH | 31 | 1 | 0 | 0 | 0 | 0 | 0 | 0 | 0 | 0 | 31 | 0 | 0 |
| Siluriformes | Pseudopimelodidae | *Pseudopimelodus pulcher* | 94.2 | LOW | 7 | 1 | 0 | 0 | 0 | 7 | 0 | 0 | 0 | 0 | 0 | 0 | 0 |
| Gymnotiformes | Gymnotidae | *Electrophorus varii* | 100.0 | HIGH | 1,234 | 5 | 753 | 253 | 46 | 144 | 38 | 0 | 0 | 0 | 0 | 0 | 0 |
| Gymnotiformes | Gymnotidae | *Gymnotus carapo* | 100.0 | HIGH | 121 | 1 | 0 | 0 | 0 | 121 | 0 | 0 | 0 | 0 | 0 | 0 | 0 |
| Gymnotiformes | Gymnotidae | *Electrophorus* sp. | 98.2 | HIGH | 8,275 | 1 | 0 | 0 | 0 | 0 | 0 | 0 | 0 | 0 | 0 | 0 | 8,275 |
| Gymnotiformes | Gymnotidae | *Gymnotus* sp. 1 | 93.5 | MODERATE | 2,152 | 1 | 0 | 0 | 0 | 0 | 0 | 0 | 0 | 2,152 | 0 | 0 | 0 |
| Gymnotiformes | Gymnotidae | *Gymnotus* sp. 2 | 88.3 | LOW | 892 | 1 | 0 | 0 | 0 | 0 | 0 | 0 | 0 | 0 | 0 | 0 | 892 |
| Gymnotiformes | Gymnotidae | *Gymnotus* sp. 3 | 88.9 | MODERATE | 1,735 | 4 | 677 | 210 | 386 | 0 | 462 | 0 | 0 | 0 | 0 | 0 | 0 |
| Gymnotiformes | Gymnotidae | *Gymnotus* sp. 4 | 92.4 | LOW | 9,561 | 2 | 0 | 0 | 0 | 0 | 136 | 0 | 0 | 0 | 0 | 0 | 9,425 |
| Gymnotiformes | Gymnotidae | *Gymnotus* sp. 5 | 95.9 | MODERATE | 477 | 4 | 228 | 90 | 123 | 0 | 36 | 0 | 0 | 0 | 0 | 0 | 0 |
| Characiformes | Characidae | *Gephyrocharax* sp. 1 | 97.6 | LOW | 36,775 | 5 | 1,681 | 8,357 | 11,493 | 9,287 | 5,957 | 0 | 0 | 0 | 0 | 0 | 0 |
| Characiformes | Characidae | *Gephyrocharax* sp. 2 | 98.2 | LOW | 462 | 5 | 117 | 92 | 103 | 100 | 50 | 0 | 0 | 0 | 0 | 0 | 0 |
| Gymnotiformes | Rhamphichthyidae | *Gymnorhamphichthys* sp. | 95.8 | HIGH | 1,299 | 4 | 226 | 579 | 361 | 0 | 133 | 0 | 0 | 0 | 0 | 0 | 0 |
| Gymnotiformes | Sternopygidae | *Eigenmannia limbata* | 98.9 | HIGH | 10,443 | 7 | 455 | 234 | 132 | 0 | 320 | 0 | 0 | 0 | 7,540 | 627 | 1,135 |
| Gymnotiformes | Sternopygidae | *Eigenmannia* sp. 1 | 87.8 | LOW | 19,687 | 6 | 86 | 707 | 104 | 0 | 64 | 0 | 0 | 0 | 0 | 5 | 18,721 |
| Gymnotiformes | Sternopygidae | *Eigenmannia* sp. 2 | 96.6 | LOW | 2,200 | 2 | 0 | 84 | 2,116 | 0 | 0 | 0 | 0 | 0 | 0 | 0 | 0 |
| Gymnotiformes | Sternopygidae | *Eigenmannia* sp. 3 | 90.2 | LOW | 224 | 2 | 0 | 75 | 0 | 149 | 0 | 0 | 0 | 0 | 0 | 0 | 0 |
| Gymnotiformes | Sternopygidae | *Eigenmannia* sp. 4 | 89.1 | LOW | 73 | 1 | 0 | 73 | 0 | 0 | 0 | 0 | 0 | 0 | 0 | 0 | 0 |
| Gymnotiformes | Sternopygidae | *Eigenmannia* sp. 5 | 98.3 | LOW | 6,246 | 1 | 0 | 0 | 0 | 0 | 0 | 0 | 0 | 0 | 0 | 0 | 6,246 |
| Gymnotiformes | Sternopygidae | *Eigenmannia* sp. 6 | 94.8 | LOW | 27 | 1 | 0 | 0 | 0 | 0 | 0 | 27 | 0 | 0 | 0 | 0 | 0 |
| Gymnotiformes | Sternopygidae | *Eigenmannia* sp. 7 | 90.0 | LOW | 7 | 1 | 7 | 0 | 0 | 0 | 0 | 0 | 0 | 0 | 0 | 0 | 0 |
| Gymnotiformes | Sternopygidae | *Eigenmannia* sp. 8 | 96.4 | LOW | 8,263 | 5 | 0 | 49 | 36 | 152 | 0 | 0 | 5,150 | 2,876 | 0 | 0 | 0 |
| Gymnotiformes | Sternopygidae | *Eigenmannia* sp. 9 | 94.3 | LOW | 8,680 | 6 | 0 | 17 | 0 | 353 | 0 | 2,828 | 0 | 3,138 | 698 | 1,646 | 0 |
| Gymnotiformes | Sternopygidae | *Eigenmannia* sp. 10 | 91.5 | LOW | 1,038 | 1 | 0 | 0 | 0 | 0 | 0 | 0 | 0 | 0 | 0 | 0 | 1,038 |
| Gymnotiformes | Sternopygidae | *Eigenmannia* sp. 11 | 92.6 | LOW | 600 | 1 | 0 | 0 | 0 | 0 | 0 | 0 | 0 | 0 | 0 | 0 | 600 |
| Gymnotiformes | Sternopygidae | *Eigenmannia* sp. 12 | 95.9 | LOW | 1,736 | 5 | 293 | 575 | 115 | 258 | 495 | 0 | 0 | 0 | 0 | 0 | 0 |
| Gymnotiformes | Sternopygidae | *Eigenmannia* sp. 13 | 93.7 | LOW | 368 | 1 | 0 | 0 | 0 | 0 | 0 | 0 | 0 | 0 | 0 | 368 | 0 |
| Gymnotiformes | Sternopygidae | *Eigenmannia* sp. 14 | 91.0 | LOW | 107 | 3 | 0 | 14 | 62 | 31 | 0 | 0 | 0 | 0 | 0 | 0 | 0 |
| Gymnotiformes | Sternopygidae | *Eigenmannia* sp. 15 | 97.7 | LOW | 113 | 3 | 33 | 42 | 0 | 0 | 38 | 0 | 0 | 0 | 0 | 0 | 0 |
| Gymnotiformes | Sternopygidae | *Sternopygus* sp. | 98.3 | HIGH | 35,592 | 10 | 1,750 | 634 | 1,806 | 239 | 706 | 0 | 13,092 | 4,650 | 6,900 | 2,408 | 3,407 |
| Gymnotiformes | Apteronotidae | *Apteronotus albifrons* | 99.4 | HIGH | 567 | 5 | 170 | 62 | 0 | 110 | 25 | 0 | 0 | 0 | 0 | 200 | 0 |
| Gymnotiformes | Apteronotidae | *Apteronotus* sp. 1 | 97.1 | HIGH | 19,574 | 2 | 0 | 0 | 0 | 0 | 0 | 0 | 0 | 0 | 0 | 6 | 19,568 |
| Gymnotiformes | Apteronotidae | *Apteronotus* sp. 2 | 89.4 | LOW | 6 | 1 | 6 | 0 | 0 | 0 | 0 | 0 | 0 | 0 | 0 | 0 | 0 |
| Gymnotiformes | Apteronotidae | *Apteronotus* sp. 3 | 84.7 | LOW | 16,538 | 5 | 0 | 142 | 147 | 0 | 0 | 2,749 | 0 | 4,631 | 0 | 0 | 8,869 |
| Gymnotiformes | Apteronotidae | *Apteronotus* sp. 4 | 97.6 | HIGH | 10,996 | 7 | 0 | 0 | 10 | 745 | 53 | 0 | 938 | 1,747 | 801 | 0 | 6,702 |
| Gymnotiformes | Apteronotidae | *Apteronotus* sp. 5 | 95.3 | MODERATE | 6,412 | 1 | 0 | 0 | 0 | 0 | 0 | 0 | 0 | 0 | 0 | 0 | 6,412 |
| Gymnotiformes | Apteronotidae | *Apteronotus* sp. 6 | 97.1 | HIGH | 4,892 | 7 | 0 | 100 | 81 | 117 | 18 | 379 | 2,233 | 0 | 0 | 0 | 1,964 |
| Gymnotiformes | Apteronotidae | *Apteronotus* sp. 7 | 97.7 | HIGH | 3,381 | 4 | 0 | 827 | 67 | 0 | 0 | 1,310 | 0 | 0 | 0 | 1,177 | 0 |
| Gymnotiformes | Apteronotidae | *Apteronotus* sp. 8 | 97.6 | HIGH | 220 | 1 | 0 | 0 | 0 | 0 | 0 | 0 | 0 | 220 | 0 | 0 | 0 |
| Gymnotiformes | Apteronotidae | *Apteronotus* sp. 9 | 96.5 | HIGH | 65 | 2 | 58 | 0 | 7 | 0 | 0 | 0 | 0 | 0 | 0 | 0 | 0 |
| Gymnotiformes | Apteronotidae | *Apteronotus* sp. 10 | 95.9 | HIGH | 19 | 2 | 0 | 6 | 0 | 13 | 0 | 0 | 0 | 0 | 0 | 0 | 0 |
| Characiformes | Ctenoluciidae | *Boulengerella maculata* | 99.4 | HIGH | 126 | 1 | 0 | 0 | 0 | 0 | 0 | 0 | 0 | 0 | 126 | 0 | 0 |
| Cichliformes | Cichlidae | *Cichla ocellaris* | 100.0 | HIGH | 52 | 1 | 0 | 0 | 0 | 0 | 0 | 0 | 0 | 52 | 0 | 0 | 0 |
| Cichliformes | Cichlidae | *Hypselecara emporalis* | 98.8 | LOW | 38 | 1 | 0 | 38 | 0 | 0 | 0 | 0 | 0 | 0 | 0 | 0 | 0 |
| Cichliformes | Cichlidae | *Aequidens* sp. 1 | 96.4 | MODERATE | 2,807 | 5 | 788 | 315 | 963 | 367 | 374 | 0 | 0 | 0 | 0 | 0 | 0 |
| Cichliformes | Cichlidae | *Aequidens* sp. 2 | 97.8 | MODERATE | 3,031 | 3 | 0 | 19 | 55 | 0 | 0 | 0 | 0 | 2,957 | 0 | 0 | 0 |
| Cichliformes | Cichlidae | *Apistogramma sp.* | 86.3 | LOW | 165 | 3 | 68 | 29 | 0 | 0 | 68 | 0 | 0 | 0 | 0 | 0 | 0 |
| Cichliformes | Cichlidae | *Astronotus* sp. | 97.6 | HIGH | 4,444 | 1 | 0 | 0 | 0 | 0 | 0 | 0 | 0 | 0 | 4,444 | 0 | 0 |
| Cichliformes | Cichlidae | *Bujurquina* sp. | 97.0 | MODERATE | 588 | 1 | 0 | 0 | 0 | 0 | 0 | 0 | 0 | 0 | 0 | 0 | 588 |
| Cichliformes | Cichlidae | *Chaetobranchus* sp. | 98.2 | HIGH | 417 | 1 | 0 | 0 | 0 | 0 | 0 | 0 | 0 | 0 | 0 | 0 | 417 |
| Cichliformes | Cichlidae | *Crenicichla* sp. 1 | 96.5 | LOW | 47 | 2 | 20 | 27 | 0 | 0 | 0 | 0 | 0 | 0 | 0 | 0 | 0 |
| Cichliformes | Cichlidae | *Crenicichla* sp. 2 | 86.0 | LOW | 971 | 2 | 34 | 0 | 0 | 0 | 0 | 0 | 0 | 0 | 937 | 0 | 0 |
| Cichliformes | Cichlidae | *Crenicichla* sp. 3 | 88.4 | LOW | 312 | 3 | 0 | 0 | 0 | 0 | 0 | 113 | 139 | 0 | 60 | 0 | 0 |
| Cichliformes | Cichlidae | *Crenicichla* sp. 4 | 87.8 | LOW | 6,265 | 8 | 607 | 614 | 1,057 | 779 | 388 | 825 | 1,926 | 0 | 69 | 0 | 0 |
| Cichliformes | Cichlidae | *Crenicichla* sp. 5 | 97.1 | LOW | 332 | 5 | 94 | 65 | 134 | 34 | 5 | 0 | 0 | 0 | 0 | 0 | 0 |
| Cichliformes | Cichlidae | *Crenicichla* sp. 6 | 98.3 | LOW | 148 | 5 | 10 | 24 | 79 | 17 | 18 | 0 | 0 | 0 | 0 | 0 | 0 |
| Cichliformes | Cichlidae | *Heros* sp. | 96.5 | MODERATE | 39 | 1 | 39 | 0 | 0 | 0 | 0 | 0 | 0 | 0 | 0 | 0 | 0 |
| Cichliformes | Cichlidae | *Mesonauta* sp. | 97.0 | LOW | 1,206 | 1 | 0 | 0 | 0 | 0 | 0 | 0 | 0 | 0 | 0 | 1,206 | 0 |
| Cichliformes | Cichlidae | *Satanoperca* sp. | 97.0 | MODERATE | 14 | 1 | 0 | 0 | 0 | 0 | 14 | 0 | 0 | 0 | 0 | 0 | 0 |
| Synbranchiformes | Synbranchidae | *Synbranchus* sp. 1 | 94.6 | HIGH | 340 | 3 | 31 | 29 | 280 | 0 | 0 | 0 | 0 | 0 | 0 | 0 | 0 |
| Synbranchiformes | Synbranchidae | *Synbranchus* sp. 2 | 94.6 | HIGH | 24 | 2 | 12 | 0 | 0 | 12 | 0 | 0 | 0 | 0 | 0 | 0 | 0 |
| Pleuronectiformes | Achiridae | *Hypoclinemus mentalis* | 99.4 | LOW | 50,324 | 1 | 0 | 0 | 0 | 0 | 0 | 0 | 0 | 0 | 0 | 50,324 | 0 |
| Perciformes | Sciaenidae | *Plagioscion squamosissimus* | 99.1 | LOW | 1,220 | 2 | 0 | 0 | 0 | 0 | 0 | 286 | 0 | 0 | 0 | 934 | 0 |
| Perciformes | Sciaenidae | *Plagioscion* sp. | 96.4 | LOW | 1,195 | 1 | 0 | 0 | 0 | 0 | 0 | 0 | 0 | 0 | 0 | 0 | 1,195 |
