## Supplementary material for "The critical role of natural history museums in advancing eDNA for biodiversity studies: a case study with Amazonian fishes": Table S4

| Order | Species | Id(%) | Read | Freq | Representative seq |
| --- | --- | --- | --- | --- | --- |
| Osteoglossiformes | *Arapaima gigas* | 100.0 | 7,050 | 4 | ACCGCGGTTATACGAGAGGCTCAAGTTAATACTATCGGCGTAAAGTGTGATTATAGGACCCAATACTAAAGCCAAAAGGCCTCAAAACTGTTATACGCCCATTGAGACTTGTAGGCTCCAAAACGAAAGTAGCTTTAAAACTTTAACCTAGAATTCACGACAGCTAGGAA |
| Cupleiformes | *Anchoviella* sp 1 | 98.8 | 18,117 | 4 | ACCGCGGTTATACGAGAGACCCTAGTTGATTAAAGCGGCGTAAAGAGTGGTTATGGAACTATTCTTTTAAAGCAGAAAACCTCTCAAACTGTTATACGCACCCAGAGGTCTGAATCCCTCACACGAAAGTGACTTTATTTATGCCTACCAGAACCCACGAAAGCTGGGAC |
| Cupleiformes | *Anchoviella* sp 2 | 95.3 | 2,970 | 1 | ACCGCGGTTATACGAGAGACCCTAGTTGATTGAAGCGGCGTAAAGAGTGGTTATGGAACTACCCTCTTAAAGCAGAAAACCTCTCAAACTGTTATACGCATCCAGAGGTCTAAATCCCTCACACGAAAGTAACTTTATTTATGCCCACCAGAACCCACGAAAGCTGGGAC |
| Cupleiformes | *Pellona* sp. 1 | 88.8 | 1,251 | 1 | ACCGCGGTTATACGAGAGGCCCTAGTTGATATGCTCGGCGTAAAGAGTGGTTATGGGGAAAAAACTAAAGCAAAAGACCCCTCAAGCAGTCATACGCACTCAGGAAGTCGAACCACTAACACGAAAGTTGCTTTATCTAACCTACCAGAGTCCACGACAGCCGGGAA |
| Cupleiformes | *Pellona* sp. 2 | 90.0 | 840 | 1 | ACCGCGGTTATACGAGAGGCCCTAGTTGATATGCTCGGCGTAAAGAGTGGTTATGGGAAAAAAACTAAAGCAAAAGACCCCTCAAGCAGTCATACGCACTCAGGAAGTCGAACCACTAACACGAAAGTTGCTTTACCTAACCTACCAGAATCCACGACAGCCGGGAA |
| Characiformes | *Characidium* sp. 1 | 98.8 | 69 | 2 | ACCGCGGTTATACGAGAGACCCTAGTTGACATCTGCGGCGTAAAGAGTGGTTAGAAATATAACATAAAACTAAAGCCAAAGATTTTCCAAGCCGTCGTACGCACCACGAAGACACGAAGCCCAAACACGAAAGTAGCTTTATTATTAAACCGACCCCACGAAAGCTAAGAA |
| Characiformes | *Characidium* sp. 2 | 85.2 | 2,685 | 6 | ACCGCGGTTATACGAGAGACCCGAGTTGATAGCTACGGCGTAAAGGGTGGTTAGGGGACAAAATTTAACTAAAGCTAAAGACCCCCTAAGCTGTCGCACGCATTACGGAGGCACGAAATCCAAAAACGAAAGTAGCTTTACAATAACCCGACCCCACAAAAGCTAAGAA |
| Characiformes | *Characidium* sp. 3 | 97.0 | 99 | 2 | ACCGCGGTTATACGAGAGACCCTAGTTGACACCTACGGCGTAAAGGGTGGTTAGAAATAAATATAAACTAAAGCTAAAGACCTTCCAAGCCGTCGTACGCACCCCGAAGGCACGAAACTCAAATACGAAAGTAGCTTTATTATAACCGACCCCACGAAAGCTAAGGA |
| Characiformes | *Characidium* sp. 4 | 89.8 | 217 | 2 | ACCGCGGTTATACGAGAGACCCTAGTTGATAGTTACGGCGTAAAGAGTGGTTAGTAAACAAACAAAACTAAAGCCAAAGACCTTCCAAGCCGTCGTACGCACCCCGAAGACACGAAGCCCAAGCACGAAAGTAGCTTTATTATCTACCGACCCCACGAAAGCTAAAAA |
| Characiformes | *Elachocharax*_sp. 1 | 85.9 | 201 | 3 | ACCGCGGTTATACGAGAGACCCTAGTTGATTAATAACGGCGTAAAGAGTGGTTAAGGGTAAATAAAATTAAAGCCAAAGACTCCCCAAGCCGTCAAACGCACCCAGAGAGCACGAAGCCCAAACACGAAAGTAGCTCTACTAATAACCCGACCCCACGAAAGCTAAGAA |
| Characiformes | *Hoplerythrinus unitaeniatus* | 99.4 | 1,332 | 5 | ACCGCGGTTATACGAGAGACCCTAGTTGATAACTACGGCGTAAAGAGTGGTTAAGAATAAAACTTAATAAAGCCAAAGACCCCCCAAGCCGTCACACGCACATGCGGGCACGAAGTTCACACACGAAAGTAGCTTTAATTAACTGACGCCACGAAAGCTAAGAA |
| Characiformes | *Hoplias malabaricus* | 99.1 | 324 | 2 | ACCGCGGTTATACGAGAGACCCTAGTTGATAAATACGGCGTAAAGAGTGGTTAGAGGACTCCCATAATAAAGCCAAAGACCCTCCAGGCCGTCACACGCATACGAGGGCACGAAGTTCACACACGAAAGTAGCTTTAAATTACCCGACGCCACGAAAGCTAAGAA |
| Characiformes | *Hoplerythrinus* sp. | 87.9 | 783 | 5 | ACCGCGGTTATACGAGAGACCCTAGTTGATATCTACGGCGTAAAGAGTGGTTAAGAATAAGATTTAATAAAGCCAAAAACCTACCAAGCCGTCACACGCATACGAAGGTATGAGACCCACACACGAAAGTAGCTTTAACAAAACTGACACCACGAAAGCTAAGAA |
| Characiformes | *Hoplias* sp. | 97.6 | 3,721 | 7 | ACCGCGGTTATACGAGATACCCTAGTTGATAAATACGGCGTAAAGAGTGGTTAGAGGACTCTTATAATAAAGCCAAAGACCCTCTGGGCCGTCACACGCATACGAGGGCACGAAGTTCACACACGAAAGTAGCTTTAAATTACCCGACGCCACGAAAGCTAAGAA |
| Characiformes | *Cynodon meionactis* | 98.8 | 1,769 | 1 | ACCGCGGTTAAACGAGAGACCCTAGTTGATCATCACGGCGTAAAGAGTGGTTAGGGGATTATCATAAATAAAGCCAAAGACCTCCCAAGCTGTCGCACGCATTCCGGAGGCGCGAAGCCCACACACGAAAGTAGCTTTAACTATTGACCCTGATGCCACGAAAGCTAAGAA |
| Characiformes | *Hydrolycus scomberoides* | 100.0 | 9,118 | 4 | ACCGCGGTTATACGAGAGACCCTAGTTGACCATCACGGCGTAAAGCGTGGTTAAAGGACCCCTATAAATAAAGCCAAAGACCTCCCAAGCTGTCGCACGCACCCCGGAGGCACGAAGCCCACACACGAAAGTAGCTTTAACTACAAACCTCGATGCCACGAAAGCTAAGAA |
| Characiformes | *Hydrolycus* sp. 1 | 87.7 | 807 | 3 | ACCGCGGTTATACGAGAGACCCAAGTTGATCGTCACGGCGTAAAGTGTGGTTAGGGGCCCCATCAAAATAAAGCCAAAGACCTCCCAAGCTGTCGCACGCTCCCTGGAGGCACGAAGCCCATACACGAAAGTAACTTTAACTAAACCCGATGCCACGAAAGCTAGGAG |
| Characiformes | *Hydrolycus* sp. 2 | 88.9 | 147 | 1 | ACCGCGGTTATACGAGAGACCCTAGTTGATTTACACGGCGTAAAGTGTGGTTAGAGAAACATCACAAATAAAGCCAAAGACCCCCCAAGCTGTCGCACGCACTACGGAGGCACGAAGCCCATACACGAAAGTAGCTTTAACTGAACCTCGATGCCACGAAAGCTAAGAA |
| Characiformes | *Serrasalmus eigenmanni* | 100.0 | 1,222 | 2 | ACCGCGGTTATACGAGAGACTCCAGTTGATAGCTACGGCGTAAAGAGTGGTTTGGGGCCCGCCCAAAATAAAGCCAAAGACCTCCCAAGCCGTCAAACGCACCCCGGAGGCACGAAGTCCTAACGCGAAAGCAACTTTACCTTCCCCGACGCCACGAAAGCTAAGAA |
| Characiformes | *Myleus* sp. | 90.4 | 949 | 1 | ACCGCGGTTATACGAGAGACCCTAGTTGATGGCTACGGCGTAAAGAGTGGTTTGGGGCCCCCACTAAAATAAAGCCAAAGACCTCTCAAGCTGTCAAACGCACCCCAGAGGCACGAAGCCCTCGTACGAAAGTAGCTTTATCTTCTCCCGACACCACGAAAGCTAAGAT |
| Characiformes | *Anodus* sp. | 100.0 | 4,483 | 3 | ACCGCGGTTATACGAGAGACCCTAGTTGATAGCCGCGGCGTAAAGAGTGGTTAGGGATACCCAACAAATAAAGCCAAAGACCTCCCAAGCTGTTACACGCATCTCGGAGGCACGAAGCCCCACTACGAAAGTGACTTTAATCTCTTCCCGACGCCACGAAAGCTAAGAA |
| Characiformes | *Abramites hypselonotus* | 100.0 | 1,304 | 1 | ACCGCGGTTATACGAGAGACCCTAGTTGATAGCTACGGCGTAAAGGGTGGTTCGAGATAAATTACAAATAAAGCTAAAGACCTTCTAAGCTGTTACAAGCACTCCGAAGACACGAAACCCCAACACGAAAGTAGCTTTACTACACTTGACGCCACGAAAGCTAAGAA |
| Characiformes | *Leporinus apollo* | 98.8 | 846 | 3 | ACCGCGGTTATACGAGAGACCCTAATTGATAGCTCACGGCGTAAAGGGTGGTTTGGGGAAACCTCAAAATAAAGCTAAAGACCTTCTAAGCCGTTACACGCATACCGAAGGCACGAGACCCTAGCACGAAAGTAGCTTTACTATTACCCCTGACGCCACGAAAACTAAGAA |
| Characiformes | *Leporinus fasciatus* | 100.0 | 31 | 1 | ACCGCGGTTATACGAGAGACCCTAATTGATAGACACGGCGTAAAGAGTGGTTTGGGAAAGCCCAAAAATAAAGCTAAAGACCTTCTAAGCCGTCACACGCACGCCGAAGGCACGAAGCCCTGACACGAAAGTAGCTTTACTATTACCCCCGACGCCACGAAAGCTAAGAA |
| Characiformes | *Anostomus* sp. | 92.4 | 39,567 | 5 | ACCGCGGTTATACGAGAGACCCTAGTTGACATAAGCGGCGTAAAGAGTGGTTTAGGGAAAACCCCAAATAAAGCCAAACATCTTCCAAGCCGTTACACGCACACTGAAGACATGAGACCCTCATACGAAGGTAGCTTTACTACTTCCCCTGACGCCACGAAAGCTAAGAA |
| Characiformes | *Leporinus* sp. 1 | 92.9 | 350 | 1 | ACCGCGGTTATACGAGAGACCCTAATTGATAGGCACGGCGTAAAGAGTGGTTTGGGACATCCTAAAAATAAAGCTAAAGACCTTCTAAGCCGTCACACGCATGCCGAAGGCGCGAGGCCCTAACACGAAAGTAGCTTTACTCTTATCCCTGACGCCACGAAAGCTAAGGA |
| Characiformes | *Leporinus* sp. 2 | 92.4 | 69 | 1 | ACCGCGGTTATACGAGAGACCCTAGTTGATAGGCACGGCGTAAAGAGTGGTTTGGGACACCCTAAAAATAAAGCTAAAGACCTTCTAAGCCGTTACACGCATGCCGAAGGCGCGAGACCCTAACACGAAAGTAGCTTTACTCTCATCCCTGACGCCACGAAAGCTAAGAA |
| Characiformes | *Leporinus* sp. 3 | 93.6 | 30 | 1 | ACCGCGGTTATACGAGAGACCCTAATTGATAGACACGGCGTAAAGAGTGGTTTGGGGTACCCTAAAAATAAAGCTAAAGACCTTCTAAGCCGTCACACGCATGCCGAAGGCGCGAGACCCTAACACGAAAGTAGCTTTACTCTTATCCCTGACGCCACGAAAGCTAAGAA |
| Characiformes | *Scheizodon knerii* | 100.0 | 46,775 | 6 | ACCGCGGTTATACGAGAGACCCTAATTGATAGGCACGGCGTAAAGAGTGGTTAGGGGTAGACTATAAATAAAGCTAAAGACCTTCTAAGCTGTCATACGCACACCGAAGGCATGAAGTCCTAATACGAAAGTAGCTTTACTATTATCCTTGACGCCACGAAAGCTAAGAA |
| Characiformes | *Leporinus* sp. 4 | 98.2 | 173,589 | 6 | ACCGCGGTTATACGAGAGACCCTAATTGATAGGTACGGCGTAAAGAGTGGTTTGGGATAACCCAAAAATAAAGCTAAAGACCTTCTAAGCCGTCACACGCATGCTGAAGGCACGAGACCCCGACACGAAAGTAGCTTTACTATTACCCCGACGCCACGAAAGCTAAGAA |
| Characiformes | *Leporinus* sp. 5 | 93.4 | 24,887 | 9 | ACCGCGGTTATACGAGAGACCCTAATTGATAGGCACGGCGTAAAGAGTGGTTTGGGACATCCTAAAAATAAAGCTAAAGACCTTCTAAGCCGTCACACGCATGCCGAAGGCGCGAGACCCTAACACGAAAGTAGCTTTACTCTTATCCCTGACGCCACGAAAGCTAAGGA |
| Characiformes | *Leporinus* sp. 6 | 92.8 | 6,498 | 7 | ACCGCGGTTATACGAGAGACCCTAGTTGATAGGCACGGCGTAAAGAGTGGTTTGGGACACCCTAAAAATAAAGCTAAAGACCTTCTAAGCCGTCACACGCATGCCGAAGGCGCGAGACCCTAACACGAAAGTAGCTTTACTCTCATCCCTGACGCCACGAAAGCTAAGAA |
| Characiformes | *Leporinus* sp. 7 | 93.5 | 4,173 | 5 | ACCGCGGTTATACGAGAGACCCTAATTGATAGACACGGCGTAAAGAGTGGTTTGGGACACCCTAAAAATAAAGCTAAAGACCTTCTAAGCCGTCACACGCATGCCGAAGGCGCGAGACCCTAACACGAAAGTAGCTTTACTCTTATCCCTGACGCCACGAAAGCTAAGAA |
| Characiformes | *Leporinus* sp. 8 | 92.9 | 665 | 4 | ACCGCGGTTATACGAGAGACCCTAATTGATAGGCACGGCGTAAAGAGTGGTTTGGGATGGACCAAAAATAAAGCTAAAGACCTTCTAAGCTGTTACACGCATACCGAAAGCACGAAACCCTAACACGAAAGTAGCTTTACTATTATCCTGACACCACGAAAGCTAAGAA |
| Characiformes | *Leporinus* sp. 9 | 93.5 | 371 | 4 | ACCGCGGTTATACGAGAGACCCTAATTGATAGACACGGCGTAAAGAGTGGTTTAGGACAACCCAAAAATAAAGCTAAAGACCTTCTAAGCCGTCACACGCATACCGAAGACATGAAGCCCCAACACGAAAGTAGCTTTACTACTGCCCCTGACGCCACGAAAGCTAAGAA |
| Characiformes | *Leporinus* sp. 10 | 92.4 | 91 | 2 | ACCGCGGTTATACGAGAGACCCTAATTGATAGGCACGGCGTAAAGAGTGGTTAGGGATAATCATAAATAAAGCTAAAGACCTTCTAAGCTGTTACACGCACACCGAAGACACGAAGCCCCAACACGAAAGTAGCTTTACTATTCCCTGACGCCACGAAAGCTAAGAA |
| Characiformes | *Leporinus* sp. 11 | 90.0 | 2,755 | 2 | ACCGCGGTTATACGAGAGACCCTAATTGATAGGTACGGCGTAAAGAGTGGTTAAGGATAAAATATAAATAAAGCTAAAGACCTTCTAAGCTGTTATACGCACACTGAAGGCACGAAGCCCTAATACGAAAGTAGCTTTACTACTATTTCTGACGCCACGAAAGCTAAGAA |
| Characiformes | *Megaleporinus* sp. | 91.7 | 959 | 1 | ACCGCGGTTATACGAGAGGCCCCAATTGACAGCCACGGCGTAAAGAGTGGTTCGGGTAAAGACCAAATGAAGCCAAAGGCCTGCCAAGCTGTTACACGCACCCCGAAGGCACGAAGTCCTAACACGAAAGTAGCTTCACCACCACCCCAACACCACGAAAGCTAAGAA |
| Characiformes | *Caenotropus labyrinthicus* | 99.4 | 14 | 1 | ACCGCGGTTATACGAGAGACCCTAGTTGATATGTACGGCGTAAAGAGTGGTTTGGGACACCTTAATAAATAAAGCCAAAGACCTCCCCAAGCTGTTGTACGCACTCCGGAGGCACGAAGCCCTAATACGAAAGTAGCTTTATTGAGCCCGACGCCACGAAAGCTAAGAA |
| Characiformes | *Potamorhina* sp. | 98.7 | 18,470 | 3 | ACCGCGGTTATACGAGAGACCCTAGTTAATATATACGGCGTAAAGAGTGGTTTGGGGTAAATACTTAATAAAGCCGAAGATCCCCCAAGCCGTCACACGCACTCCGGAGACACGAAGCCCCAGCACGAAAGTAGCTTTACAAAGACCCCCGACGCCACGAAAGCTAAGAA |
| Characiformes | *Cyphocharax* sp. | 95.9 | 845 | 1 | ACCGCGGTTATACGAGAGACCCTAGTTGATATACACGGCGTAAAGAGTGGTTTGGGACATACTCAAAATAAAGCCGAAGATCTCCCAAGCCGTCACACGCACCCCGGAGACACGAAGCCCTGATACGAAGGTAGCTTTATTAATACCCGACGCCACGAAAGCTAAGAA |
| Characiformes | *Prochilodus harttii* | 100.0 | 90,591 | 6 | ACCGCGGTTATACGAGAGACCCTAGTTGATATATACGGCGTAAAGAGTGGTTTGGAATAACCAAGTAATATAGCCAAAGACCTCCCAAGCCGTCACACGCACCCCGGAGGCACGAAGCCCAAACACGAAAGTAGCTTTATTAACTTACCGACGCCACGAAAGCTAAGAA |
| Characiformes | *Semaprochilodus* sp. 1 | 100.0 | 15,460 | 5 | ACCGCGGTTATACGAGAGACCCTAGTTGATATACACGGCGTAAAGAGTGGTTTGGGACAAACCAAATAATAGAGCCAAAGACCTCCCAAGCCGTCACACGCACCCCGGAGGCACGAAGCCCAAGCACGAAAGTAGCTTTATTACACCCCCGACGCCACGAAAGCTAAGAA |
| Characiformes | *Chilodus* sp. | 94.0 | 37 | 2 | ACCGCGGTTATACGAGAGACCCCAGTTGATATACACGGCGTAAAGAGTGGTTTAGGGTATATAATAAATAAAGCCAAAGACCCCCTAGGCTGTTGAACGCACCACGGAGGAGCGAAGCCCCAATACGAAAGTAGCTTTATAAGACCTGACGCCACGAAAGCTAAGAA |
| Characiformes | *Prochilodus* sp. | 87.6 | 46,510 | 3 | ACCGCGGTTATACGAGAGACCCTAGTTGATATATACGGCGTAAAGAGTGGTTAGGAAACATCTATATTTAAAGCCAAACTCCTCCTAAGCCGTCGCACGCCCCCTGGAGGCAGGAAGCCCAAACACGAAAGTAGCTTTATATACCCGACCCCACGAAAGCTAAGAA |
| Characiformes | *Semaprochilodus* sp. 2 | 84.1 | 272 | 2 | ACCGCGGTTATACGAGAGACCCTAGTTGATATACACGGCGTAAAGAGTGGTTAGGAAACACTTATACTTAAAGCCAAACTCCTCCTAAGCCGTCGCACGCCCCCTGGAGGCAGGAAACCCATATACGAAAGTAGCTTTATATACCCGACCCCACGAAAGCTAAGAA |
| Characiformes | *Semaprochilodus* sp. 3 | 84.7 | 31 | 2 | ACCGCGGTTATACGAGAGACCCTAGTTGATATACACGGCGTAAAGAGTGGTTAGGAAACATCTATATTTAAAGCCAAACTCCTCCTAAGCCGTCGCACGCCCCCTGGAGGCAGGAAACCCATATACGAAAGTAGCTTTATATACCCGACCCCACGAAAGCTAAGAA |
| Characiformes | *Semaprochilodus* sp. 4 | 91.6 | 1,694 | 2 | ACCGCGGTTATACGAGAGACCCTAGTTGATACACACGGCGTAAAGAGTGGTTAGGGACAGACCAAAAAATAAAGCCAAAGACCTCCCAAGCCGTCACACGCACCCCGGAGATACGAGACCCAAGCACGAAAGTAGCTTTATCACCCCCCCCCCGACGCCACGAAAGCTAAGAA |
| Characiformes | *Semaprochilodus* sp. 5 | 83.5 | 179 | 2 | ACCGCGGTTATACGAGAGACCCTAGTTGATATACACGGCGTAAAGAGTGGTTAGGAAACATTTATATTTAAAGCCAAACTCCTCCTAAGCCGTCGCACGCCCCCTGGAGGCAGGAAACCCATATACGAAAGTAGCTTTATATGCCCGACCCCACGAAAGCTAAGAA |
| Characiformes | *Acestrorhynchus falcatus* | 99.4 | 11,546 | 5 | ACCGCGGTTACACGAGAGACTCAAGTTAATAGACTACGGCGTAAAGCGTGGTTAGGGGCCCTTAATAACTAAAGCCAAAGATCTTCTATGTCGTCGCACGCAACGCGAAGAAACGAAGCCCAAACACGAAAGTAGCTTTATTTCCCCTGACCCCACGAAAGCTAAGAT |
| Characiformes | *Acestrorhynchus* sp. 1 | 91.7 | 833 | 1 | ACCGCGGTTACACGAGAGACTCAAGTTAATAGACTACGGCGTAAAGCGTGGTTAGGGGCCCCAACAACTAAAGCCAAAGATCTTCTATGTCGTCGCACACACCGCGAAGCTGCGAAGCCCAAACACGAAAGTTGCTTTACTAACCCTGACCCCACGAAAGCTAAGAC |
| Characiformes | *Acestrorhynchus* sp. 2 | 84.0 | 7 | 1 | ACCGCGGTTATACGAAAGACCCAAGTTGATAGATACGGCGTAAAGCGTGGTTAAAGGAACACCATAATTAAAGCCAAAGACCTCCAATGCTGTCAAATATATTTCGGAGGCACGAAGCCCAAACACGAAAGTAGCTTTATTTTCCTTGACTCCACGAAAGCTAAGAA |
| Characiformes | *Chalceus erythrurus* | 100.0 | 8,741 | 5 | ACCGCGGTTATACGAGAGACCCTAGTTGATAGCTACGGCGTAAAGAGTGGTCTAGGACCCACAGCAAATTAAAGCCAAAGACCTCCCAAGCTGTCGCACGCACCCGGAGGCACGAAGCCCAAACACGAAGGTAGCTTTATCACATCTCCTAACCCCACGAAAGCTAAGAA |
| Characiformes | *Chalceus macrolepidotus* | 100.0 | 4,158 | 2 | ACCGCGGTTATACGAGAGACCCTAGTTGATAGATACGGCGTAAAGAGTGGTCTAGGACCCACAACAAAATAAAGCCAAAGACCTCCCAAGCTGTCGCACGCACCCCGGAGGCACGAAGCCCAAACACGAAGGTAGCTTTATTACATTCTCCTAACCCCACGAAAGCTAAGAA |
| Characiformes | *Charax pauciradiatus* | 99.4 | 10,186 | 5 | ACCGCGGTTATACGAGAGACCCAAATTAATAGCTACGGCGTAAAGAGTGGTTTGGGGTAAAATTAATAAAGCCGAATACTCTCCTGGCCGTCGCACGCATTTTGAGAGCATGAAGCCCTATAACGAAAGTAGCTTTACCAATATTTTCCTGACCCCACGAAAGCTAAGAA |
| Characiformes | *Charax* sp. 1 | 99.4 | 32 | 1 | ACCGCGGTTATACGAGAGACCCTAATTAATAGCTACGGCGTAAAGAGTGGTTTAGGGTAAAAATTAATAAAGCCGAAGATTCTCATGGCCGTTGTACGCATTCTGAGAATATGAAGCCCCAACACGAAAGTAGCTTTACCAGTAATTTCCTGACCCCACGAAAGCTAAGAA |
| Characiformes | *Ctenobrycon hauxwellianus* | 100.0 | 491 | 3 | ACCGCGGTTATACGAGAGACCCTAGTTGATAAACACGGCGTAAAGAGTGGTTAGGATAAAAGAAAAATAAAGTCAAATGCCCTCTAGGCCGTTACACGCATTCTGAGAACATGAAGCCCCACTACGAAAGTAACTTTACTATTTCCGACCCCACGAAAGCTAAGAA |
| Characiformes | *Jupiaba ocellata* | 98.8 | 1,150 | 4 | ACCGCGGTTATACGAGAGACCCTAGTTGATAAATACGGCGTAAAGAGTGGTTATGGGAAAAACAAATAAAGTCAAACAACCTCTTAGCTGTTATACGCATTATGAGAGTATGAAGCCCCCTCACGAAAGTAACTTTAATATCTCCTGACCCCACGAAAGCTAAGAA |
| Characiformes | *Moenkhausia lepidura* | 99.4 | 782 | 3 | ACCGCGGTTATACGAGAGACCCTAGTTGATAGCTACGGCGTAAAGCGTGGTTAGGAGAACAATATAAATAAAGTCAAACAATCTCTCGGCCGTTATACGTTATCTGAGAATATGAAGTCCTACCACGAAAGTAACTTTAATTTTTCTGACCCCACGAAAACTGAGGA |
| Characiformes | *Myloplus rubripinnis* | 99.3 | 232 | 4 | ACCGCGGTTATACGAGAGACCCTAGTTGACAGCTACGGCGTAAAGAGTGGTTTGGGGCACCCCATAAAAATAAAGCCAAAGACCTTCCAAGCTGTCAAACGCACTCCGGAGGCACGAAACCCCAACACGAAAGTAGCTTTACCTTCACCCCGACGCCACGAAAGCTAAGAA |
| Characiformes | *Pygocentrus nattereri* | 99.9 | 19,766 | 9 | ACCGCGGTTATACGAGAGACTCCAGTTGATAGCTACGGCGTAAAGAGTGGTTTGGGGCCCCACCCAAAATAAAGCCAAAGACCTCCCAAGCCGTCAAACGCACCCCGGAGGCACGAAGTCCTAACGCGAAAGCAACTTTACCTCCCCCGACGCCACGAAAGCTAAGAA |
| Characiformes | *Astyanax* sp. 1 | 97.4 | 4,347 | 5 | ACCGCGGTTATACGAGAGACCCCAGTTGATATACACGGCGTAAAGAGTGGTTAGGAGAATAAAAAAATAAAGTCAAATAACCTCTAAGCCGTTGTACGCATTACGAGAACATGAAGACCCCCCACGAAAGTAACTTTACCACCTCCGACCCCACGAAAGCTAAGAA |
| Characiformes | *Astyanax* sp. 2 | 97.0 | 4,542 | 5 | ACCGCGGTTATACGAGAGACCCCAGTTGATATATACGGCGTAAAGAGTGGTTAGGAGAATAAATAAATAAAGTCAAATGACCTCTAAGCCGTTGTACGCATTACGAGAACAAGAAGCCCCCCACGAAAGTAACTTTACCACCTCCGACCCCACGAAAGCTAAGAA |
| Characiformes | *Astyanax* sp. 3 | 88.6 | 3,172 | 5 | ACCGCGGTTATACGAGAGACCCGAGTTGATAAATACGGCGTAAAGAGTGGTTAGGAAAACAGAGAAAATAAAGTCAAATGTTCTCCCAGCTGTTCTACGCACTACGAGAATATGAAGCCCCAACGAAAGTAACTTTACCACTTCCGACCCCACGAAAGCTAAGAA |
| Characiformes | *Astyanax* sp. 4 | 96.4 | 23 | 1 | ACCGCGGTTATACGAGAGACCCCAGTTGATACACACGGCGTAAAGAGTGGTTAGGAGAATAAAAAAAATAAAGTCAAATAACCTCTAAGCCGTTGTACGCATTACGAGAACATGAAGGCCCCCCACGAAAGTAACTTTACCACCTCCGACCCCACGAAAGCTAAGAA |
| Characiformes | *Astyanax* sp. 5 | 95.8 | 14 | 1 | ACCGCGGTTATACGAGAGACCCCAGTTGATATACACGGCGTAAAGAGTGGTTAGGAGAATAAAAAAAATAAAGTCAAATAACCTCTAAGCCGTTGTACGCATTACGAGAACATGAAGGCCCCCAACGAAAGTAACTTTACCACCTCCGACCCAACGAAAGCTAAGAA |
| Characiformes | *Brycon* sp. 1 | 96.4 | 228,753 | 5 | ACCGCGGTTATACGAGAGGCCCTAGTTGATAGACACGGCGTAAAGGGTGGTTAGGGATAACTATTAATTAAAGCCGAAGACCTCCAAGTCCGTCATACGATCTCGAAGGCACGAAGTCCCGACACGAAAGTAGCTTTATTACCCCCGACCCCACGAAAGCTAAGAA |
| Characiformes | *Brycon* sp. 2 | 89.2 | 158,513 | 6 | ACCGCGGTTATACGAGAGACCCTAGTTGATAGACCACGGCGTAAAGGGTGGTTAGGGATAACCATTTATTAAAGCCAAAGATCCCCCAAGCTGTCATACGATCCGGAGGATCGAAGCCCCAACACGAAAGTAGCTTTATTACCCCTGACCCCACGAAAACTAAGGA |
| Characiformes | *Brycon* sp. 3 | 87.0 | 54,689 | 5 | ACCGCGGTTATACGAGGGGCCCTAGTTGATGGATCTCGGCGTAAAGGGTGGTTAGGGGTAACCCCTGAATAAAGCCAAAGACCTCCCAACCTGTTATACGGTCCGAAGGCACGAAGTCCCAACACGAAAGTAGCTTTATTATAACCCCGACCCCACGAAAGCTAAGAA |
| Characiformes | *Brycon* sp. 4 | 96.4 | 2,095 | 5 | ACCGCGGTTATACGAGAGACCCTAGTTGATAGACACGGCGTAAAGGGTGGTTAGGGATAACTATTAATTAAAGCCGAAGACCTCCAAGTCCGTCATACGATCTCGGAAGCACGAAGCCCCAACACGAAAGTAGCTTTATTACCCCCGACTCCACGAAAGCTAAGAA |
| Characiformes | *Brycon* sp. 5 | 93.0 | 1,451 | 1 | ACCGCGGTTATACGAGAGACCCTAATTGATAGCTACGGCGTAAAGAGTGGTTTAGGGTAAAAAATGAATAAAGCCAAAGATCCTCCTGGCTGTAGCACGCACTCCGAGGACACGAAGCCCCACTACGAAGGTAGCTTTAAAATATCCCCTGACCCCACGAAAGCTAAGAA |
| Characiformes | *Bryconops* sp. 1 | 96.0 | 5,379 | 5 | ACCGCGGTTATACGAGAGACTCCAGTTGATAGCCACGGCGTAAAGAGTGGTTAAGAAAATATATTAAATAAAGCCAAAGGCCCCCTAAGCCGTCTCACGCACTCCGAGGGTACGAAGCCCAAATACGAAAGTAGCTTTACCTCTTACTATTCTGACCCCACGAAAGCTAAGAA |
| Characiformes | *Bryconops* sp. 2 | 96.5 | 200 | 5 | ACCGCGGTTATACGAGAGACTCCAGTTGATAGCTACGGCGTAAAGAGTGGTTAAGAAAACATATTAAATAAAGCCAAAGGTCCCCTAAGCCGTCTCACGCACTCCGGGGATACGAAACCCAAATACGAAAGTAGCTTTACCCCCTGCTATTCTGACCCCACGAAAGCTAAGAA |
| Characiformes | *Charax* sp. 2 | 95.3 | 9,541 | 3 | ACCGCGGTTATACGAGAGACCCTAATTAATAGCTACGGCGTAAAGAGTGGTTAAGGATAAAAATTAATAAAGCCGAAGATTCTCCTGGCTGTTGCACGCACTCTGAGAACATGAAGCCCCATTACGAAAGTAGCTTTACCAATAATCTCCTGACCCCACGAAAGCTAAGAA |
| Characiformes | *Charax* sp. 3 | 89.2 | 5,298 | 2 | ACCGCGGTTATACGAGAGACCCTAATTAATAGCTACGGCGTAAAGAGTGGTTAAGGGTAAAATTAATAAAGCCGAACATTCTCCTAGCTGTTGAACGCATCTGAGAACATGAAGCCCTACAACGAAAGTAGCTTTACTTCACTTTCCTGACTCCACGAAAGCTAAGAA |
| Characiformes | *Charax* sp. 4 | 95.9 | 142 | 2 | ACCGCGGTTATACGAGAGACCCTAATTAATAGCTACGGCGTAAAGAGTGGTTTAGGGTAAAAATTAATAAAGCCGAAAATTCTCCTGGCTGTTGTACGCACTCTGAGAATATGAAGCCCCATTACGAAAGTAGCTTTACCAATATCTCCTGACCCCACGAAAGCTAAGAA |
| Characiformes | *Charax* sp. 5 | 98.2 | 38 | 1 | ACCGCGGTTATACGAGAGACCCAAATTAATAGCTACAGCGTAAAGAGTGGTTTGGGGTAAAATTAATAAAGCCGAATACTCTCCTGGCCGTCGCACGCATTTTGAGAGCATGAAGCCCTATAACGAAAGTAGCTTTACCAACATTTTCCTGACCCCACGAAAGCTAAGAA |
| Characiformes | *Colossoma* sp. 1 | 93.6 | 224,686 | 6 | ACCGCGGTTATACGAGAGACCCTAGTTGATGACTACGGCGTAAAGAGTGGTTTGGGACCCTCACTAAAATAAAGCCAAAGACCTCTCAAGCTGTCAAACGCACCCCGGAGGCACGAAGCCCTAACACGAAAGTAGCTTTACCTTCTCCCGACGCCACGAAAGCTAAGAC |
| Characiformes | *Colossoma* sp. 2 | 89.3 | 2,473 | 2 | ACCGCGGTTATACGAGAGACCCTAGTTGATGGCTACGGCGTAAAGAGTGGTTTGGGGCCCCCCACTAAAATAAAGCCAAAGACCTCTCAAGCTGTCAAACGCACCCCAGAGGCACGAAGCCCTCGTACGAAAGTAGCTTTATCTTCTCCCGACACCACGAAAGCTAAGAT |
| Characiformes | *Corynopoma* sp. | 92.4 | 1,580 | 2 | ACCGCGGTTATACGAGAGACCCTAATTGATAGCTACGGCGTAAAGAGTGGTTTAGGATAAAAAAGAAATAAAGCCAAAGATCCTCCCAGCTGTCGTACGCATTTTGAGGACACGAAGCCCCTCCACGAAAGTAGCTTTAATAAAATTTTCTTGACCCCACGAAAGCTAAGAA |
| Characiformes | *Cyanocharax* sp. 1 | 91.9 | 41 | 1 | ACCGCGGTTATACGAGAGACCCTAATTGATAGCTACGGCGTAAAGAGTGGTTAAGGATAAAAATAAATAAAGCCAAAGATCCTCCTGGCTGTAGCACGCACTCTGAGGACATGAAGCCCCACTACGAAAGTAACTTTAATTTACACCCTGATGCCACGAAAGCTAAGAA |
| Characiformes | *Cyanocharax* sp. 2 | 90.8 | 2,606 | 2 | ACCGCGGTTATACGAGAGACCCCAATTGATAGCTACGGCGTAAAGAGTGGTTTAAGATAAAAAAGAAATAAAGCCAAAGATCCTCCCAGCTGTCGTACGCATTTTGAGGACACGAAGCCCCCTAACGAAAGTAGCTTTACTAAAACTTTTCTTGACCCCACGAAAGCTAAGAA |
| Characiformes | *Hasemania* sp. | 85.8 | 39 | 3 | ACCGCGGTTAAACGAGAGGGCCCTAGTTGATAGATACGGCGTAAAGGGTGGTTAGGGATATAGAACAAATAAAGTCAAATATTCTCAAAGCTGTTATAAGCCTCCGAGAGCATGAAGCCCCATCACGAAAGTAACTTTATCACACCCCGACCCCACGAAAGCTAAGAA |
| Characiformes | *Hemigrammus* sp. 1 | 91.7 | 212 | 3 | ACCGCGGTTATACGAGAGACCCTAGTTGATAGATACGGCGTAAAGAGTGGTTAAGGGTAAAACTAAATAAAGTCAAACAATCTCTGAGCCGTTGTACGCGATATGAGAGCATGAGGCCCCCCCACGAAAGTAACTTTAATAATTCCTGACCCCACGAAAGCTAAGAA |
| Characiformes | *Hyphessobrycon* sp. 1 | 92.7 | 17 | 1 | ACCGCGGTTATACGAGAGACCCTAGTCGATAGATACGGCGTAAAGCGTGGTTAAGGGATTAATAAATAAAGTCAAAGAGTCTTTTAGCTGTTATAAGCCCACAAGAACACGAAGTCCAATCACGAAAGTAACTTTACCTCTTCCTGACCCCACGAAAGCTAAGGA |
| Characiformes | *Hyphessobrycon* sp. 2 | 86.6 | 264 | 4 | ACCGCGGTTACACGAGTAGACCCTAGTTGATAGATTACGGCGTAAAGAGTGGTTAGGGGTACAAAACCTAATAAAGTTAAACTAACTCCAAGCTGTTGATAAGCACTACGAGACTGTGAAACCCAATGACGAAAGTAACTTTATTTCTCCTGAATCCACGAAAGCTAAGAA |
| Characiformes | *Iguanodectes* sp. | 94.2 | 145 | 3 | ACCGCGGTTATACGAGAGACCCCAGTTGATGGCTACGGCGTAAAGAGTGGTTATGGATTTTAAATTTAACTAAAGCCAAAGTACCCCTAGGTCGTTTTACACATTACGGACGCACGAAGCCCAAAAACGAAGGTAGCTTTACTCCTATCCTGACCCCACGAAAGCTAAGAA |
| Characiformes | *Jupiaba* sp. 1 | 82.1 | 400 | 1 | ACCGCGGTTATACGAGAGACCCCAATTAATTATGTACGGCGTAAAGAGTGGTTAAGAGACCCAATAAATAAAGCTGAGGACCCTCCAAGCTGTTATACGCACCCGAGAACACTAAACCCCATTACGAAAGTAGCTTTAATAACCCCTGACCCCACGAAAGCTAAGAC |
| Characiformes | *Jupiaba* sp. 2 | 94.6 | 3,339 | 7 | ACCGCGGTTATACGAGAGACCCCAGTTGATATATACGGCGTAAAGAGTGGTTATGGGAATAAAACAAATAAAGTCAAACAACCTCTTAGCTGTTATACGCATTATGAGAACATGAAGCCCCCTTACGAAAGTAACTTTACCACCCCCTGACCCCACGAAAGCTAAGAA |
| Characiformes | *Moenkhausia* sp. 1 | 97.6 | 22 | 1 | ACCGCGGTTATACGAGTGGCCCTAGTTGATAGATACGGCGTAAAGAGTGGTTATGGGTAAAAACGAAATAAAGTTAAACAATCTCTTGGCTGTTATACGCGCTGTTAGATCATGAAACCCCTTAACGAAAGTGACTTTAGTATATCCTGACCCCACGAAAGCTAAGAA |
| Characiformes | *Moenkhausia* sp. 2 | 93.6 | 684 | 5 | ACCGCGGTTATACGAGAGACCCAAGTTGATAGGTGCGGCGTAAAGAGTGGTTAAGGGTAAAAAAATAAATAAAGTTAAACAACCTCTATAAGCTGTTATACGCGTCACGAGGACATGAAGCCCATTCACGAAAGTAACTTTACTACCTCCTGACCCCACGAAAGCTAAGAA |
| Characiformes | *Moenkhausia* sp. 3 | 97.1 | 5 | 1 | ACCGCGGTTATACGAGAGACCCAAGTTGATAGACACGGCGTAAAGAGTGGTTAAGGATGAAGAATAAATAAAGTCAAACGACCTCTATGAGCTGTTATACGCATTGCGAGGACATGAAGCCCACTCACGAAAGTAACTTTACCACCTCCTGACCCCACGAAAGCTAAGAAA |
| Characiformes | *Moenkhausia* sp. 4 | 91.1 | 688 | 5 | ACCGCGGTTACACGAATAGACCCTAGTTGATAGATACGGCGTAAAGAGTGGTTAGGGAAATAAATTAATAAAGTTAAACATTCTCCAAATTGTTATACATCACTTCGAGCATATGAAACCCACTCACGAAAGTAACTTTAAATACCCCGACTCCACGAAAGCTAAGAA |
| Characiformes | *Moenkhausia* sp. 5 | 89.9 | 14 | 1 | ACCGCGGTTACACGAGTGCACCCTAGTTGATAGACACGGCGTAAAGGGTGGTTAAGGAACAAACCTAATAAAGTTAAACATCCTCCAAATTGTTATACATAACTTCGAGTATATGAAACCCACTCACGAAAGTAACTTTAACAGCCCCGACTCCACGAAAGCTAAGAA |
| Characiformes | *Paracheirodon* sp. | 88.1 | 89 | 1 | ACCGCGGTTATACGAGAGACCCTAGTTGATAAATACGGCGTAAAGAGTGGTTAGGGATTTAGAATAAATAAAGTCAAATAACCTCTAGGCCGTTACAAGCAATACGAGAATACGAAGCCCCATCACGAAAGTAACTTTACTCCTCCTGACCCCACGAAAGCTAAGAA |
| Characiformes | *Poptella* sp. | 95.2 | 4,950 | 5 | ACCGCGGTTATACGAGAGACCCCAGTTGATATATACGGCGTAAAGAGTGGTTATGGGAATAAAACAAATAAAGTCAAACAACCTCTTAGCTGTTATACGCATTATGAGAACATGAAGCCCCCCTACGAAAGTAACTTTACCACCCCCTGACCCCACGAAAGCTAAGAA |
| Characiformes | *Serrasalmus* sp. | 98.2 | 3,335 | 1 | ACCGCGGTTATACGAGAGACTCCAGTTGATAACTACGGCGTAAAGAGTGGTTTGGGGCCCCACCCAAAATAAAGCCAAAGACCTCCCAAGCCGTCAAACGCACCCCGGAGGCACGAAGTCCTAACGCGAAGGCAGCTTTACCTTCCCCGACGCCACGAAAGCTAAGAA |
| Characiformes | *Bryconops affinis* | 100.0 | 9,344 | 5 | ACCGCGGTTATACGAGAGACCCTAATTGATAGACTACGGCGTAAAGAGTGGTTTAGAACAAAAATAAATAAAGCCAAAGATCCTCCTGGCCGTCGCACGCACTTCGAGGATACGAAGCCCCACTACGAAAGTAGCTTTACTATTAACTTTTCTGACCCCACGAAAGCTAAGAA |
| Characiformes | *Tetragonopterus* sp. | 96.5 | 3,300 | 6 | ACCGCGGTTATACGAGAGACCCTAATTGATAGATTACGGCGTAAAGAGTGGTTTAGAACAAAAATAAATAAAGCCAAAGGTCCTCCTGGCCGTCGCACGCATTTCGAGGATATGAAGCCCCACTACGAAGGTAGCTTTACTATAACTTTTCTGACCCCACGAAAGCTAAGAA |
| Characiformes | *Tyttocharax* sp. | 94.8 | 1,572 | 1 | ACCGCGGTTATACGAGAGACCCTAATTGATAGCTACGGCGTAAAGAGTGGTTTAGGATAAAAACGAATAAAGCCAAAGATCCTCCTGGCTGTAGCACGCATTCCGAGGACACGAAGCCCTACTACGAAGGTAGCTTTACTTATGCTCTCCTGACCCCACGAAAGCTAAGAA |
| Characiformes | *Triportheus* sp. 1 | 99.8 | 232 | 3 | ACCGCGGTTATACGAGAGACCCAAGTTGATAAATACGGCGTAAAGAGTGGTTAAGGATAATAAAGAAATAAAGCCAAAGGCCTTCCAAGCTGTCACACGCACCTCTAAGGTACGAAGCCCAAACACGAAAGTAGCTTTACCACCTGCCTGACCCCACGAAAGCTAAGAA |
| Characiformes | Thoracocharax sp. | 93.9 | 1,275 | 1 | ACCGCGGTTATACGAGAGGCCCTAGTTAATAGATACGGCGTAAAGAGTGGTTAGGAGCCAATATAACTAAAGCCAAAGACCCCCTAAGCTGTCATACGCACCCGGAGGCACGAAGCCCTATAACGAAAGTAGCTTTAAAATCTCTGACCCCACGAAAACTAAGAC |
| Characiformes | *Elachocharax* sp. 2 | 84.3 | 84 | 3 | ACCGCGGTTATACGAGAGACCCTAGTTGATATAACACGGCGTAAAGGGTGGTTAGGGATATAATAAAACTAAAGCCAAAGCCCCCTTGAGCTGTTATACGCATTCCAGGGGCACGAAGTCCAAAAACGAAAGTAGCTTCATTAACATCCTGACCCCACGAAAGCTAAGAAA |
| Characiformes | *Triportheus* sp. 2 | 95.7 | 5,745 | 3 | ACCGCGGTTATACGAGAGACCCTAGTTGATAGACACGGCGTAAAGAGTGGTTAGGGGCAATAAACAAATAAAGCCAAAGACCTTCCAAGCTGTCATACGTACCCCGAAGGCACGAAGCCCAAACACGAAAGTAGCTTTACCCCCCCCCCGACCCCACGAAAGCTAAGAA |
| Characiformes | *Triportheus* sp. 3 | 91.3 | 6,537 | 3 | ACCGCGGTTATACGAGAGGCCCTAGTTGATAGATACGGCGTAAAGAGTGGTTAAGGGTAATAAATAAATAAAGCCAAAGGCCTTCAAAGCTGTCACACGCACCCCGAAGGTACGAAGCCCGAACACGAAAGTAGCTTTACTATACTCCTGACCCCACGAAAGCTAAGGA |
| Siluriformes | *Helogenes* sp. | 90.7 | 24 | 1 | ACCGCGGTTATACGAAAGACTCCAGTTGATAGGCACGGTGTAAAGAGTGGTTAAGAAGTATTATTAATAAAGCCAAACACTCTCCAAGCCGTCATACGCACTCTGAGAGCACGAAGCCCAAACACGAAAGTAGCTTTAAATCTATTATACCTGAACCCACGAAAGCCAAGGA |
| Siluriformes | *Corydoras* sp. 1 | 95.8 | 152 | 4 | ACCGCGGTTATACGAAAGACTCTAGTTGATAAACACGGCGTAAAGGGTGGTTAGGGTAACAATAAAATAAAGTTAAAGACTTACCAAGCCGTCATACGCCCATGAAAGAACGAAAACCAAAAACGAAAGTAACTTTATTATACACCCGAACCCACGAAAGCTAAGAA |
| Siluriformes | *Corydoras* sp. 2 | 98.2 | 13 | 2 | ACCGCGGTTATACGAAAGACTCTAGTTGACAAACACGGCGTAAAGGGTGGTTAGGATAACAACAAAATAAAGCTAAAGATTTACCAAGCCGTCATACGCCCATGTAAAAGCGAAAACCACAAACGAAAGTAACTTTATTACACCATCCGAATCCACGAAAGCTAAGAA |
| Siluriformes | *Dekeyseria amazonica* | 100.0 | 86 | 1 | ACCGCGGTTATACGAAAGACCCCAGTTTATATACACGGCGTAAAGGGTGGTTAGGGGACAAACAAAATAAAGCCAAAGACCCTCTAAGCCGTCATACGCTCACGAAGGTACTAAGCCCAAACACGAAGGTAGCTTTACTAAACATACCCGACTCCACGAAAGCTGAGAA |
| Siluriformes | *Hypostomus plecostomus* | 98.8 | 113 | 1 | ACCGCGGTTATACGAAAGACCCTAGTTTATAGGTACGGCGTAAAGGGTGGTTAGGGGACAAACAAAATAAAGCCAAAGACCTCCTAAGCCGTCATACGCTCACGGAGGCACGAAGCCCAAACACGAAAGTAGCTCTACCAAACATGCCCGACTCCACGAAAGCTGGGAA |
| Siluriformes | *Lasiancistrus saetiger* | 98.8 | 1,837 | 1 | ACCGCGGTTATACGAAAGACCCTAATTTATAGACACGGCGTAAAGGGTGGTTAGGGGACAAACAAAATAAGGCCAAAGACCTCCTAAGCCGTCATACGCTCACGGAGGCACGAAGCCCAAACACGAAAGTAGCTTTACCAAACATGCCCGACTCCACGAAAGCTAGGAA |
| Siluriformes | *Ancistrus* sp. | 97.0 | 938 | 5 | ACCGCGGTTATACGAAAGACCCTAGTTTATAAACACGGCGTAAAGGGTGGTTAGGGGACAAACAAAATAAAGCCAAAGACTCCCTAAGCCGTCATACGCTCACGGAAATACGAAGTCCAAACACGAAAGTAGCTTTACCAAACATGCCCGACTCCACGAAAGCTGGGGG |
| Siluriformes | *Hypoptopoma* sp. 1 | 87.7 | 116 | 1 | ACCGCGGTTATACGAAAGACCCCAGTTTATAGATACGGCGTAAAGGGTGGTTATGGGACAAAATAAAATAGGGCCAAAAACCCACTAGGCTGTTATAAGCATACGAGGACACGAAGCCCCTATACGAAAGTAGCCCTACCAAAAAATGCCCGACGCCACGAAAGCGGGGGG |
| Siluriformes | *Hypoptopoma* sp. 2 | 98.1 | 314 | 3 | ACCGCGGTTATACGAAAGACCCTAGTTTATAGACACGGCGTAAAGGGTGGTTATGGGATAGAACAAAATAGAGCCAAAGACCCCCTAAGCCGTCATACGCCCACGGGGGCACGAAGCCCCCTCACGAAAGTAGCTCTACTAAAAATATACCCGACCCCACGAAAGCTGGGAA |
| Siluriformes | *Hypoptopoma* sp. 3 | 91.9 | 1,139 | 1 | ACCGCGGTTATACGAAAGACCCTAGTTTATAGGCACGGCGTAAAGGGTGGTTATGGGATTTATTAAACTAGAGCCAAAGACCCCCTAAGCCGTCATACGCCCACGGAGGCACGAAGCCCCCATACGAAAGTAGCTCTACCAGAAAATGCCCGACCCCACGAAAGCTGGGAA |
| Siluriformes | *Hypostomus* sp. 1 | 98.2 | 190 | 1 | ACCGCGGTTATACGAAAGACCCTAGTTTATAGTTACGGCGTAAAGGGTGGTTAGGGGACGAACAAAATAAAGCCAAAGACCTCCTAAGCCGTCATACGCTCACGGAGGCACGAAGCCCAGACACGAAAGTAGCTTTACCAAACATGCCCGACTCCACGAAAGCTGGGAA |
| Siluriformes | *Hypostomus* sp. 2 | 98.2 | 111 | 1 | ACCGCGGTTATACGAAAGACCCTAGTTTATAGGTACGGCGTAAAGGGTGGTTAGGGGACAAACAAAATAAAGCCAAAGACCTCCTAAGCCGTCATACGCTCACGGAGGCACGAAGCCCAAACACGAAAGTAGCTCTACCAAACATGCCCGATTCCACGAAAGCTGGGAA |
| Siluriformes | *Pseudacanthicus* sp. | 97.0 | 746 | 1 | ACCGCGGTTAAACGAAAGACCCCAGTTTATAGACACGGCGTAAAGGGTGGTTAGGGGTTAAACAGAATAAAGCCGAAGATTTCTTAAGCCGTCATACGCTCATAAAAACACGAAGCCCGAACACGAAAGTAGCTTTACTAAACATACCCGACTCCACGAAAGCTGGAAA |
| Siluriformes | *Spatuloricaria* sp. | 98.3 | 482 | 1 | ACCGCGGTTATACGAAAGACCCCAGTTTATAGATACGGCGTAAAGGGTGGTTAAGGAAAACCCAAAATAAGGCTAAAGACCTCCTAAGCAGTCATACGCTCCACGGAAACACGAAATCCAAATACGAAAGTTGCCCTAATTAAACTTACCTGACCCCACGAAAGCTGAAAA |
| Siluriformes | *Hemidoras morrisi* | 99.4 | 14 | 1 | ACCGCGGTTATACGAAAGACCCTAGTTGATAGATCACGGCGTAAAGGGTGGTTAAGGAGAACAAAAATAAAGCTAAAGATCCTCTAAGCTGTCATACGCTTTCCGAAGACATGAGATCCAACCACGAAAGTAGCTTTAAACTGTCCTGACGCCACGAAAGCCAAGAA |
| Siluriformes | *Liosomadoras morrowi* | 99.4 | 25 | 1 | ACCGCGGTTATACGAAAGACCCAAGTTGATTAATTACGGCGTAAAGGGTGGTTAAGGGTAAATTGAAAATAAGGCTAAAGACTCTCCAGGCTGTCATACGCTCTCCGAGATAACGAGACCCTCACACGAAAGTATCCTTAAAATTTAAACCCTGACGCCACGAAAGCCAAGAA |
| Siluriformes | *Megalodoras uranoscopus* | 100.0 | 9 | 1 | ACCGCGGTTATACGAAAGACCCTAGTTGATAGACTACGGCGTAAAGGGTGGTTAAGGATTACAAAAATAAAGCTAAAGATCCTCTAGGCTGTCATACGCTTTCCGAGGAGATGAAACCCAGCCACGAAAGTAGCTTTAAACCCCTCCTGACACCACGAAAGCCAAGAA |
| Siluriformes | *Platydoras costatus* | 100.0 | 40 | 2 | ACCGCGGTTATACGAAAGACCCTAGTTGATAGACTACGGCGTAAAGGGTGGTTAAGGAATACAAAAATAAAGCTAAAGATCCTCTAGGCTGTCATACGCTTTCCGAGGAGATGAAGCCCCACCACGAAAGTAGCTTTAAGCATCTCCTGACACCACGAAAGCCAAGAA |
| Siluriformes | *Pterodoras granulosus* | 98.8 | 94 | 1 | ACCGCGGTTATACGAAAGACCCTAGTTGATAGCCCACGGCGTAAAGGGTGGTTAAGGGGCCACAAAAATAAAGCTAAAGATCCTCTAGGCTGTCATACGCTCTCCGAGGATATGAGACCCCACCACGAAAGTAGCTTTAAGCATCTCCTGAAACCACGAAAGCCAAGAA |
| Siluriformes | *Trachydoras nattereri* | 99.4 | 34 | 1 | ACCGCGGTTATACGAAAGACCCTAGTTGATGGATCACGGCGTAAAGGGTGGTTAAGGAGAACAAAAATAAAGCTAAAAATCCTCTAAGCTGTCATACGCTTTCTGAAGACATGAGACCCAACCACGAAAGTAGCTTTAAACTTTCCTGACACCACGAAAGCCAAAGA |
| Siluriformes | *Nemadoras* sp. | 96.4 | 346 | 1 | ACCGCGGTTATACGAAAGACCCTAGTTGATAGATCACGGCGTAAAGGGTGGTTAAGGAGAACAAAAATAAAGCTAAAGATCATCTAAGCTGTCATACGCTTTCCGAAGACATGAAACCCAGTCACGAAAGTAGCTTTAAACCGTCCTGACACCACGAAAGCCAAGAA |
| Siluriformes | *Opsodoras* sp. | 96.4 | 19 | 1 | ACCGCGGTTATACGAAAGACCCTAGTTGATAGATCACGGCGTAAAGGGTGGTTAAGGAATACAAAAATAAAGCTAAAGATCCTCTAAGCTGTCATACGCTTTCCGAAGACATGAGATCCAACTACGAAAGTAGCTTTAAATTACCCTGACGCCACGAAAGCCAAGAA |
| Siluriformes | *Ageneiosus* sp. | 95.3 | 40 | 1 | ACCGCGGTTATACGAAAGACCCTAGTTGATGGGCACGGCGTAAAGGGTGGTTAGGGACTACAAAAAATAGAGCTAAAGATCCTCTAGGTCGTCATACACATTACGAGGAAACGAGACCCAAATACGAAAGTAACTCTAAACAAATCCTGATGCCACGAAAGCCAAGAA |
| Siluriformes | *Auchenipterus* sp. | 97.1 | 13 | 1 | ACCGCGGTTATACGAAAGACCCTAGTTGATAGCCACGGCGTAAAGGGTGGTTAAGGAGCCTAAAAAATAAAGCTAAAGACCCTCTAGGCCGTCATACGCATTACGAGGCAAACGAAGCCCCAACACGAAAGTAGCTTTAGACAAACATCCTGATGCCACGAAAGCCAAGAAA |
| Siluriformes | *Centromochlus* sp. | 89.5 | 835 | 1 | ACCGCGGTTATACGAAAGACCCTAGTTGATTGACACGGCGTAAAGGGTGGTTAAGGCTAAATCACTAAGAATAAGGCTAAAGGCCCACTAAGCTGTCATACGCTTCCCGAGGGAACGAAACCCCATAACGAAAGTAGCCTCAAAATAAACCTGACACCACGAAAGCCAAGGA |
| Siluriformes | *Chasmocranus* sp. | 90.4 | 132 | 4 | ACCGCGGCTATACGAAAGACCCCAGTTGACAGGAACGGCGTAAAGGGTGGTTAAGGACAACCACAAATAAAGTTAAAGAGCCCCCAGGCTGTCGCACGCACTCTGGAGACTCGAGGCCCACACACGAAGGTAACTTTAGAACACTGCCCGACCCCACGAAAGCCAAGAA |
| Siluriformes | *Imparfinis* sp. | 81.5 | 26 | 1 | ACCGCGGTTATACGAGAGACCCTAGTTGAAAAAGACGGCGTAAAGAGTGGTTAAAAATGAATAAAATTAAAGTTAAAGACTCTCTAAGTTGTCGCACACACCCAGAGAACACGAAGCCCAAATACGAAAGTGACTTTATTAATTATTTGACCCCACGAAAGCTAAGAA |
| Siluriformes | *Pimelodella* sp. 1 | 98.3 | 427 | 1 | ACCGCGGTTATACGAAAGACCCTAGTTGATAGGCACGGCGTAAAGGGTGGTTAGGGATATATCAAAAATAAAGTTAAAGAGCCTCTAAGCTGTAGCACGCATCCCGAGAGCTCGAGACCCAAACACGAAAGTAACTTTAAAATACTTCACCTGACCCCACGAAAGCTAAGAA |
| Siluriformes | *Pimelodella* sp. 2 | 97.7 | 35 | 1 | ACCGCGGTTATACGAAAGACCCTAGTTGATAGACACGGCGTAAAGGGTGGTTAGGGATTTCTAAAAATAAAGTTAAAGAGCCTCTAAGCTGTCGCACGCATCCCGAGAGCTCGAAACCCAAACACGAAAGTAACTTTAAAATACTTCACCTGACCCCACGAAAGCTAAGAA |
| Siluriformes | *Rhamdia* sp. | 97.7 | 22 | 1 | ACCGCGGTTATACGAAAGACCCCAGTTGATAGAAACGGCGTAAAGGGTGGTTAGGGAAGACTCAAAATAAAGTTAAAGAGCCTCTAAGCTGTCGCACGCACCCCGAGAGCTCGAGAACCGAACACGAAAGTAACTTTAAAAACACCTCACCTGACCCCACGAAAGCTAAGAAA |
| Siluriformes | *Hypophthalmus edentatus* | 98.8 | 472 | 2 | ACCGCGGTTATACGAAAGACCCTAGTTGATAGCCACGGCGTAAAGGGTGGTTAAGGTATCCTACAAATAAAGCTAAAGAGCCTCTAAGCCGTCGCACGCATTCCGAGAGCCCGAAACCCAAACACGAAAGTAGCTTTAAAAACAACACACCTGACTCCACGAAAGCTAAGAA |
| Siluriformes | *Pimelodella cristata* | 98.9 | 1,359 | 7 | ACCGCGGTTATACGAAAGACCCTAGTTGATAGGCACGGCGTAAAGGGTGGTTAGGGATATATCAAAAATAAAGTTAAAGAGCCTCTAAGCTGTCGCACGCATCCCGAGAGCTCGAGACCCAAACACGAAAGTAACTTTAAAATACTTCACCTGACCCCACGAAAGCTAAGAA |
| Siluriformes | *Pseudoplatystoma reticulatum* | 99.1 | 2,064 | 6 | ACCGCGGTTATACGAAAGACCCTAGTTGATAGCCACGGCGTAAAGGGTGGTTAAGGTAACTAATAAATAAAGCTAAAGAGCCTCTAAGCCGTCGCACGCATTCCGAGAGCTCGAAGCCCAAACACGAAAGTAGCTTTAAAATAAAGCACACCTGACCCCACGAAAGCTAAGAA |
| Siluriformes | *Hypophthalmus* sp. | 97.1 | 102 | 1 | ACCGCGGTTATACGAAAGACCCTAGTTGATAGCCACGGCGTAAAGGGTGGTTAAGGTAAATTACAAATAAAGCTAAAGAGCCTCTAAGCCGTCGCACGCATTCCGAGAGCCCGAAACCCAAATACGAAAGTAGCTTTAAAACCAACACACCTGACTCCACGAAAGCTAAGAA |
| Siluriformes | *Pimelodus* sp. 1 | 97.2 | 739 | 5 | ACCGCGGTTATACGAAAGACCCTAGTTGATAGCCACGGCGTAAAGGGTGGTTAAGGTGTATAACAAATAAAGCTAAATAGCCTCTAAGCCGTCGCACGCATTCCGAGAGCTCGAAACCCAAAAACGAAAGTAGCTTTAAAATAAACAAACCTGACCCCACGAAAGCTAAGAA |
| Siluriformes | *Pimelodus* sp. 2 | 98.3 | 619 | 4 | ACCGCGGTTATACGAAAGACCCTAGTTGATAGCCACGGCGTAAAGGGTGGTTAAGGTACATAACAAATAAAGCTAAATAGCCTCTAGGCCGTCGCACGCATTCCGAGAGCTCGAAACCCAAAAACGAAAGTAGCTTTAAAATAAACAGACCTGACCCCACGAAAGCTAAGAA |
| Siluriformes | *Pimelodus* sp. 3 | 97.7 | 58 | 4 | ACCGCGGTTATACGAAAGACCCTAGTTGATAGCCACGGCGTAAAGGGTGGTTAAGGTATATAATAAATAAAGCTAAATAGCCTCTAGGCCGTCGCACGCATTCCGAGAGCTCGAAACCCAAAAACGAAAGTAGCTTTAAAATAAACAAACCTGACCCCACGAAAGCTAAGAA |
| Siluriformes | *Pinirampus pinirampu* | 99.4 | 4 | 1 | ACCTCGGTTATACGAAAGACCCTAGTTGATGGCCACGGCGTAAAGGGTGGTTAAGGTACCTAACAAATAAAGCTAAATAGCCTCTAAGCCGTCGAACGCGCCCTGAGAACTAGAAACTCAAAAACGAAAGTAGCTTTAAAACAACCCCACCTGACCCCACGAAAGCTAAGAA |
| Siluriformes | *Pimelodus* sp. 4 | 97.8 | 2,522 | 5 | ACCGCGGTTATACGAAAGACCCTAGTTGATAGCCACGGCGTAAAGGGTGGTTAAGGTGTATAACAAATAAAGCTAAATAGCCTCTAAGCCGTCGCACGCATTCCGAGAGCTCGAAACCCAAAAACGAAAGTAGCTTTAAAATAAACAGACCTGACCCCACGAAAGCTAAGAA |
| Siluriformes | *Pimelodus* sp. 5 | 94.8 | 140 | 1 | ACCGCGGTTATACGAAAGACCCTAGTTGACAGCCACGGCGTAAAGGGTGGTTAAGGCATATAACAAATAAAGCTAAATAGCCCCTAAGCCGTCGCACGCACTCCGAGAGCTAGAAACCCAAAAACGAAAGTAGCTTTAAAACAAAAAGACCTGACCCCACGAAAGCTAAGAA |
| Siluriformes | *Pimelodus* sp. 6 | 92.5 | 7 | 1 | ACCGCGGTTATACGAAAGACCCTAGTTGATAAACACGGCGTAAAGGGTGGTTAAGGTAAACAACAAATAAAGCTAAATAGCCTCTAAGCCGTCGCACGCACCCCGAGAGCTCGAAGTCCAAAAACGAAAGTAGCTTTAAAATAACCCCACACCTGACCCCACGAAAGCTAAGAA |
| Siluriformes | *Pimelodus* sp. 7 | 98.3 | 10 | 2 | ACCGCGGTTATACGAAAGACCCTAGTTGATAGCCACGGCGTAAAGGGTGGTTAAGGTAAACAAATAAATAAAGCTAAACAGCCCCTAGGCCGTCGTACGCATTACGGGAGCTAGAAACCCAAACACGAAAGTTGCTTTAAATTAAATATACCTGACTCCACGAAAGCTAAGAA |
| Siluriformes | *Pimelodus* sp. 8 | 97.7 | 14 | 1 | ACCGCGGTTATACGAAAGACCCTAGTTGATAGCCACGGCGTAAAGGGTGGTTAAGGTATATAATAAATAAAGCTAAATAGCCTCTAAGCCGTCGCACGCACTCCGAGAGCTCGAAGCCCAAAAACGAAAGTAGCTTTAAAACAAACACACCTGACCCCACGAAAGCTAAGAA |
| Siluriformes | *Pimelodus* sp. 9 | 95.4 | 4 | 1 | ACCGCGGTTATACGAAAGACCCTAGTTGATAGCCACGGCGTAAAGGGTGGTTAAGGTACATAACAAATAAAGCTAAATAGCCTCTAAGCCGTCGCACGCACTCCGAGAGCTCGAAGCCCAAAAACGAAAGTAGCTTTAAAACAAACCCACACCTGACCCCACGAAAGCTAAGAA |
| Siluriformes | *Pseudoplatystoma tigrinum* | 100.0 | 183 | 2 | ACCGCGGTTATACGAAAGACCCTAGTTGATAGCCACGGCGTAAAGGGTGGTTAAGGTAACTAATAAATAAAGCTAAAGAGCCTCTAAGCCGTCGCACGCATTCCGAGAGCTCGAAGCCCAAACACGAAAGTAGCTTTAAAACAAAACACACCTGACCCCACGAAAGCTAAGAA |
| Siluriformes | *Hemisorubim platyrhynchos* | 100.0 | 110 | 1 | ACCGCGGTTATACGAAAGACCCTAGTTGATAGCCACGGCGTAAAGGGTGGTTAAGGTATAAAATAAATAAAGCCAAAGAGCCTCTAAGTCGTCGTACACATTCCGAGTGCTCGAAGCCCAAATACGAAGGTTGCTTTAACATAACATACACCTGACCCCACGAAAGCTAAGAA |
| Siluriformes | *Brachyplatystoma vaillantii* | 100.0 | 25 | 1 | ACCGCGGTTATACGAAAGACCCCAGTTGATAGCCACGGCGTAAAGGGTGGTTAAGGTAAATAATAAATAAAGCTAAAGAGCCTCTAAGTCGTCGCACGCATTCCGGGAGCTCGAAGCCCAGACACGAAAGTAGCTTTAAAATAAAATACACCTGACTCCACGAAAGCTAAGAA |
| Siluriformes | *Sorubimichthys planiceps* | 99.4 | 431 | 1 | ACCGCGGTTATACGAAAGACCCTAGTTGATAGCCACGGCGTAAAGGGTGGTTAAGGTAGACAACAAATAAAGCTAAAGAACCTCTAAGCTGTCGCACGCATTCCGAGAACTCGAAGCCCAAACACGAAAGTAGCTTTAAAACAAAAACACCTGAACCCACGAAAGCTAAGAA |
| Siluriformes | Pseudoplatystoma sp. | 95.4 | 39 | 1 | ACCGCGGTTATACGAAAGACCCTAGTTGATAGCCACGGCGTAAAGGGTGGTTAAGGTAAATAATAAATAAAGCTAAATAGCCTCTAAGTCGTCGTACACATCCCGAGAGCTCGAAGCCCAAACACGAAAGTAGCTTTAAAATCAAGCACACCTGACCCCACGAAAGCTAAGAA |
| Siluriformes | *Sorubim elongatus* | 100.0 | 282 | 1 | ACCGCGGTTATACGAAAGACCCTAGTTGATAGCTGCGGCGTAAAGGGTGGTTAAGGTATATAACAAATAAAGCTAAAGAACCTCCAAGCCGTCGTACGCATTATGAGAGCTCGAAACCCAAGCACGAAAGTAGCTTTAAAAAATCACACCTGACCCCACGAAAGCTAAGAA |
| Siluriformes | Sorubim sp. | 97.1 | 338 | 2 | ACCGCGGTTATACGAAAGACCCTAGTTGATAGCTACGGCGTAAAGGGTGGTTAAGGTATATAACAAATAAAGCTAAAGAGCCTCCAAGCCGTCGTACGCATTCTGAGAGCTCGAAACCCTAACACGAAAGTAGCTTTAAAAAATAACACCTGACCCCACGAAAGCTAAGAA |
| Siluriformes | *Phractocephalus hemioliopterus* | 100.0 | 316 | 4 | ACCGCGGTTATACGAAAGACCCTAGTTGATAGCTACGGCGTAAAGGGTGGCTAAGGCAGACAACAAATAAAGCCAAAGAGCCTCTAAGCCGTCGCACGCACTCCGAGGCCTCGAAATCCAAACACGAAAGTAGCTTTAAAACAAAACACACCTGACCCCACGAAAGCTAAGAA |
| Siluriformes | *Zungaro* sp. | 86.3 | 39 | 2 | ACCGCGGTTATACGAAAGACCCTAATTAATAGCCACGGCGTAAAGGGTGGTTAAGGAAAAATCACTAAATAAAGCTAAAGGGCCCCCAAGCTGTCGCACGCACCCTGGAAGCTCGAGACCCCTACACGAAAGTAGCTTTAAAATTACCCCGCCTGACCCCACGAAAGCTAAGAA |
| Siluriformes | *Zungaro jahu* | 100.0 | 31 | 1 | ACCGCGGTTATACGAAAGACCCTAGTTGATAGCCACGGCGTAAAGGGTGGTTAAGGTAAATAATAAATAAAGCTAAAGGGCCTCTAAGCCGTCGCACGCATTCCGAGAGCTCGAAACCCAAGCACGAAAGTAGCTTTAAAATAAAACACACCTGACCCCACGAAAGCTAAGAA |
| Siluriformes | *Pseudopimelodus pulcher* | 94.2 | 7 | 1 | ACCGCGGTTATACGAAAGACCCTAGTTGATATTTACGGCGTAAAGGGTGGTTAGGGAAAAACATAAATAAAGCTAAAGAGCCTCTAAGCCGTCGTACGCACCACGAAAGCTCGAAACCCAAGAACGAAAGTAGCTTTAAAATATACTACCTGACCCCACGAAAGCTAAGAA |
| Gymnotiformes | *Electrophorus varii* | 100.0 | 1,234 | 5 | ACCGCGGTTATACGAGAGACTCCAGTTGACAGAACTCGGCATAAAGAGTGGTTATAATACACCCAAATAAAGCCAAAAATCTCTAGAGCCGTCATACGCTTTCCAGAGACATGAAGCCCTAACACGAAAGTAGCTTTATGACATTGAACCCACGAAAGCTAAGAA |
| Gymnotiformes | *Gymnotus carapo* | 100.0 | 121 | 1 | ACCGCGGTTATACGAGAGACCCTAGTTGATAATTACGGCGTAAAGAGTGGTTAAGGAACTACACAATTAAAGCCAAACACTTCCCCGGCTGTTATACGCTCCCGGAAATAACGAAACCCAAACGCGAAAGCAGCTTTATATTATAAGCCTGACCCCACGAAAGCTAAGAT |
| Gymnotiformes | *Electrophorus* sp. | 98.2 | 8,275 | 1 | ACCGCGGTTATACGAGAGACTCCAGTTGACAGAACTCGGCGTAAAGAGTGGTTATGGTACACCTAAATAAAGCCAAAAATCTCTAGAGCCGTCATACGCTTTCCAGAGACATGAAGCCCTAACACGAAAGTAACTTTATTACATTAGACCCACGAAAGCTAAGAA |
| Gymnotiformes | *Gymnotus* sp. 1 | 93.5 | 2,152 | 1 | ACCGCGGTTATACGAGAGACCCTAGTTGATAACTACGGCGTAAAGAGTGGTTAAGAAACCATACAATTAAAGCCAAACTCTTCCCCGGCTGTTATACGCTCCCGGAAATAATGAAGCCCAAACGCGAAAGCAGCTTTATTTTATATCCTGACCCCACGAAAGCTAAGAC |
| Gymnotiformes | *Gymnotus* sp. 2 | 88.3 | 892 | 1 | ACCGCGGTTATACGAGAGGCCCTAGTTGATAACCACGGCGTAAAGAGTGGTTAAGGAGTTATACAACTTAAAGCCAAACACTTCCCCAACTGTTACAAGCTAATGAAAGTAAGAAGCCCAAACACGAAAGCAGCTTTATCCTATAAGCCTGACCCCACGAAAGCTAAGAC |
| Gymnotiformes | *Gymnotus* sp. 3 | 88.9 | 1,735 | 4 | ACCGCGGTTATACGAGAGGCCCTAGTTGATAACCACGGCGTAAAGAGTGGTTAAGGAGTTATACAACTTAAAGCCAAACACTTCCCCAACTGTTACAAGCTAATGAAAGTAAGAAACCCAAACACGAAAGCAGCTTTATATTATAAGCCTGACCCCACGAAAGCTAAGAC |
| Gymnotiformes | *Gymnotus* sp. 4 | 92.4 | 9,561 | 2 | ACCGCGGTTATACGAGAGGCCCTAGTTGATAACTACGGCGTAAAGAGTGGTTAAGGACTATCACAACTAAAGCTAAACATTTCTCCGGCTGTTATACGCTACCAGAAATAACGAAACCCGCACGCGAAAGCAGCTTTATTTAATAAACCTGATCCCACGAAAGCTAGGGT |
| Gymnotiformes | *Gymnotus* sp. 5 | 95.9 | 477 | 4 | ACCGCGGTTATACGAGAGGCCCTAGTTGATAGCTACGGCGTAAAGAGTGGTTAAGGGATATCACAACTAAAGCTAAACATTTCCCCGGCTGTCATACGCTACCAGAAATAACGAAACCCACACGCGAAAGCAGCTTTATCTTATAAACCTGACCCCACGAAAGCTAAGGT |
| Characiformes | *Gephyrocharax* sp. 1 | 97.6 | 36,775 | 5 | ACCGCGGTTATACGAGAGACCCTAGTTGATAAACACGGCGTAAAGAGTGGTTAAGGATAAAAACAAATAAAGTCAAACAATCTCTTAGCTGTTATACGCATTATGAGAACATGAAGCCCTTCTACGAAAGTAACTTTACTAATACCTGACCCCACGAAAGCTAAGGA |
| Characiformes | *Gephyrocharax* sp. 2 | 98.2 | 462 | 5 | ACCGCGGTTATACGAGAGACCCTAGTTGATAAACACGGCGTAAAGAGTGGTTAAGGATAAAAACGAATAAAGTCAAACAATCTCTTAGCTGTTATACGCATTATGAGAACATGAAGCCCTTCTACGAAAGTAACTTTACTATTACCTGACCCCACGAAAGCTAAGAAA |
| Gymnotiformes | *Gymnorhamphichthys* sp. | 95.8 | 1,299 | 4 | ACCGCGGTTATACGAGAGACCCCAGTTGATACCAGCGGCGTAAAGGGTGGTTAAGGGCCCTTACAACTAAAGCTAAACACTTCCCCGGCTGTTGTACGCAAACTGGAAGCCTGAAATCCTGACACGAAAGTAGCTTTATTTTCCCCGACCCCACGAAAGCTGAGAA |
| Gymnotiformes | *Eigenmannia limbata* | 98.9 | 10,443 | 7 | ACCGCGGTTATACGAGAGGCCCTAGTTGATAGCCACGGCGCAAAGAGTGGTTAAGGAGCCCCACTAAATAAAGTCGAACACTTCCTAGGCCGTTATACGTTTTCTAGAAGCACGAAACCCAATTACACGAAAGCAACTTTATACTAAATAACCTGACCCCACGAAAGCTAAGAA |
| Gymnotiformes | *Eigenmannia* sp. 1 | 87.8 | 19,687 | 6 | ACCGCGGTTATACGAGAGGCCCTAGCTGATAGGCACGGCGCAAAGAGTGGTTAAGGAGGCCCTCCCATAAAGCCAAACAATTCCTAGGCCGTCATACGTTTTCTAGAAACACGAAGCCCAATCACACGAAAGCAGCTTTACACCCACCAACCTGACCCCACGAAAGCTAAGGC |
| Gymnotiformes | *Eigenmannia* sp. 2 | 96.6 | 2,200 | 2 | ACCGCGGTTATACGAGAGGCCCTAGTTGATAGCCACGGCGCAAAGAGTGGTTAAGGAACCCCATCAAATAAAGTCGAACACTTCCTAGGCCGTTATACGTTTTCTAGAAACACGAAGCCCAATTACACGAAAGCAACTTTATACTAAACAACCTGACCCCACGAAAGCTAAGAA |
| Gymnotiformes | *Eigenmannia* sp. 3 | 90.2 | 224 | 2 | ACCGCGGTTATACGAGAGGCCCTAGCTGATAGGCACGGCGCAAAGAGTGGTTAAGGAGGCCCTTCAATAAAGCCAAACAACTCCTAGGCCGTCATACGTTTTCTAGAAACACGAAGCCCAATCACACGAAAGCAGCTTTATACTAACCAACCTGACCCCACGAAAGCTAAGAC |
| Gymnotiformes | *Eigenmannia* sp. 4 | 89.1 | 73 | 1 | ACCGCGGTTATACGAGAGGCCCTAGCTGATAGGCACGGCGCAAAGAGTGGTTAAGGAGACCCTTCCATAAAGCCAAACAATTCCTAGGCCGTCATACGTTTTCTAGAAACACGAAGCCCAATCACACGAAAGCAGCTTTACACCAACCAACCTGACCCCACGAAAGCTAAGAC |
| Gymnotiformes | *Eigenmannia* sp. 5 | 98.3 | 6,246 | 1 | ACCGCGGTTATACGAGAGGCCCTAGTTGACAGCCACGGCGCAAAGAGTGGTTAAGGAGTCCTACCAAATAAAGTCAAATACTTCCTAGGCCGTTATACGTTTTCTAGAAACATGAAGCCCAATTACACGAAAGCAACTTTATGCTAAACAACCTGACCCCACGAAAGCTAAGAA |
| Gymnotiformes | *Eigenmannia* sp. 6 | 94.8 | 27 | 1 | ACCGCGGTTATACGAGAGGCCCTAGTTGATAGCTACGGCGCAAAGAGTGGTTAAGGAGCCCACCAAAATAAAGTCAAACACTTCCTAGGCCGTTATACGTTTTCTAGAAGCACGAAGCCCAATTACACGAAAGCAGCTTTATTCTACAACCTGACCACACGAAAGCTAAGAA |
| Gymnotiformes | *Eigenmannia* sp. 7 | 90.0 | 7 | 1 | ACCGCGGTTATACGAGAGGCCCTAGTTGATAGCCACGGCGTAAAGAGTGGTTAAGGAACCTTTTTCCCCCCCCCCATAAAGTCAAACACTTCCTAGGCCGTTATACGTTTTCTAGAAGCACGAAGCCCAATCACACGAAAGCAACTTTATACAACACCTGACCCCACGAAAGCTAAGAA |
| Gymnotiformes | *Eigenmannia* sp. 8 | 96.4 | 8,263 | 5 | ACCGCGGTTATACGAGAGGCCCTAGTTGATAGCTACGGCGCAAAGAGTGGTTAAGGAGCCCACCCAAATAAAGTCAAACACTTCCTAGGCCGTTATACGTTTTCTAGAAGCACGAAGCCCAATTACACGAAAGCAACTTTATTCTACAACCTGACCCCACGAAAGCTAAGAA |
| Gymnotiformes | *Eigenmannia* sp. 9 | 94.3 | 8,680 | 6 | ACCGCGGTTATACGAGAGGCCCTAGTTGATAGCCACGGCGCAAAGAGTGGTTAAGGAGTCCTACTAATAAAGTCAAACGCTTCCTAGGCCGTTATACGCTTTCTAGAAACATGAAGCCCAATTACACGAAAGTAACTTTATACCAAACAACCTGACCCCACGAAAGCTAAGAA |
| Gymnotiformes | *Eigenmannia* sp. 10 | 91.5 | 1,038 | 1 | ACCGCGGTTATACGAGAGGCCCTAGTTGATAGCCACGGCGTAAAGAGTGGTTAAGGAACCTTTCCCCCCCCCATAAAGTCAAACACTTCCTAGGCCGTTATACGTTTTCTAGAAGCACGAAGCCCAATCACACGAAAGCAACTTTATACAACACCTGACCCCACGAAAGCTAAGAA |
| Gymnotiformes | *Eigenmannia* sp. 11 | 92.6 | 600 | 1 | ACCGCGGTTATACGAGAGGCCCTAGTTGATAGCCACGGCGTAAAGAGTGGTTAAGGAACCTTTTTCCCCCCATAAAGTCAAACACTTCCTAGGCCGTTATACGTTTTCTAGAAGCACGAAGCCCAATCACACGAAAGCAACTTTATACAACACCTGACCCCACGAAAGCTAAGAA |
| Gymnotiformes | *Eigenmannia* sp. 12 | 95.9 | 1,736 | 5 | ACCGCGGTTATACGAGAGGCCCTAGTTGATAGCCACGGCGCAAAGAGTGGTTAAGGAGCTCAACCAAATAAAGTCAAACACTTCCTAGGCCGTTATACGTTTTCTAGAAATACGAAGCCCAATTACACGAAAGCAACTTTATCCTGTAACCTGACCCCACGAAAGCTAAGGA |
| Gymnotiformes | *Eigenmannia* sp. 13 | 93.7 | 368 | 1 | ACCGCGGTTATACGAGAGGCCCTAGTTGACAGCCACGGCGCAAAGAGTGGTTAAGGAGTCCTACTAATAAAGTCAAACGCTTCCTAGGCCGTTATACGCTTTCTAGAAACATGAAGCCCAATCACACGAAAGTAACTTTATATCAAACAACCTGACCCCACGAAAGCTAAGAA |
| Gymnotiformes | *Eigenmannia* sp. 14 | 91.0 | 107 | 3 | ACCGCGGTTATACGAGAGGCCCTAGTTGATAGCCACGGCGTAAAGAGTGGTTAAGGAACCTTTTTCCCCCCCCATAAAGTCAAACACTTCCTAGGCCGTTATACGTTTTCTAGAAGCACGAAGCCCAATCACACGAAAGCAACTTTATTCAACACCTGACCCCACGAAAGCTAAGAA |
| Gymnotiformes | *Eigenmannia* sp. 15 | 97.7 | 113 | 3 | ACCGCGGTTATACGAGAGGCCCTAGTTGATAGCCACGGCGCAAAGAGTGGTTAAGGAGTCCTACAAATAAAGTCAAATATTTCCTAGGCCGTTATACGTTTTCTAGAGACATGAAGCCCAATTACACGAAAGCAACTTTATGCTAAACGACCTGACCCCACGAAAGCTAAGAA |
| Gymnotiformes | *Sternopygus* sp. | 98.3 | 35,592 | 10 | ACCGCGGTTATACGAGAGACCCTAGTTGATTAATAACGGCGTAAAGAGTGGTTAAGGGTAAATAAAATTAAAGCCAAAGACTCCCCAAGCCGTCAAACGCACCCAGAGAGCACGAAGCCCAAACACGAAAGTAGCTCTACTAATAACCCGACCCCACGAAAGCTAAGAA |
| Gymnotiformes | *Apteronotus albifrons* | 99.4 | 567 | 5 | ACCGCGGTTATACGAAAGACCCAAGTTGATAGTCACGGCGTAAAGAGTGGTTAAGGGAACTAATAAATAAAGCCAAACACTTCCCAGGCCGTTGCACGTTTTCTGGAAACACGAAGCCCAATCACGAAAGTAGCTTTACATAAACCACCTGACCCCACGAAAGCTAAGAA |
| Gymnotiformes | *Apteronotus* sp. 1 | 97.1 | 19,574 | 2 | ACCGCGGTTATACGAAAGACCCAAGTTGATAGTCACGGCGTAAAGAGTGGTTAAGGGAACTAATAAATAAAGCCAAATGCTTCCCAGGCCGTTGCACGTTTTCTGGAACCACGAAGCCCAATCACGAAAGTAGCTTTACAAAAACCACCTGACCCCACGAAAGCTAAGAA |
| Gymnotiformes | *Apteronotus* sp. 2 | 89.4 | 6 | 1 | ACCGCGGTTATACGAAAGGCCCAAGTTGATAACTACGGCGTAAAGAGTGGTTAAGGAGACCTGAAAATAAAGCCAAATACTTCCCAGGCCGTTGCACGTTTTCTGGAAGCACGAAGCCCAATCACGAAAGTAGCTTTACCACCACCACCTGAACCCACGAAAGCTAAGAA |
| Gymnotiformes | *Apteronotus* sp. 3 | 84.7 | 16,538 | 5 | ACCGCGGTTATACGAGAGACTCTAGTTGATATATACGGCGTAAAGAGTGGTTAAGGAACACAACAATTAAAGCCAAACACTTCCCCGGCCGTCGCACGTTTTCCGGAAGCACGAAGCCCAAACACGAAAGTAGCTTTATAAATACCTGACCCCACGAAAGCTGAGGA |
| Gymnotiformes | *Apteronotus* sp. 4 | 97.6 | 10,996 | 7 | ACCGCGGTTATACGAAAGACCCAAGTTGATAGTCACGGCGTAAAGAGTGGTTAAGGGAACTAATAAATAAAGCCAAATACTTCCCAGGCCGTTGCACGTTTTCTGGAAACACGAAGCCCAATTACGAAAGTAGCTTTACAAAAACCACCTGACCCCACGAAAGCTAAGAA |
| Gymnotiformes | *Apteronotus* sp. 5 | 95.3 | 6,412 | 1 | ACCGCGGTTATACGAAAGACCCAAGTTGATAGTCACGGCGTAAAGAGTGGTTAAGGAAAACGATAAATAAAGCCAAACCCTTCCCAGGCCGTTGCACGTTTTCTGGAAACACGAAGCCCAATTACGAAAGTAGCTTTACATAAATCACCTGACCCCACGAAAGCTAAGAA |
| Gymnotiformes | *Apteronotus* sp. 6 | 97.1 | 4,892 | 7 | ACCGCGGTTATACGAAAGACCCAAGTTGATAGTCACGGCGTAAAGAGTGGTTAAGGGTACTAGTAAATAAAGCCAAACACTTCCCAGGCCGTTGCACGTTTTCTGGAAACACGAAGCCCAACCACGAAAGTAGCTTTACAAAAACCACCTGACCCCACGAAAGCTAAGAA |
| Gymnotiformes | *Apteronotus* sp. 7 | 97.7 | 3,381 | 4 | ACCGCGGTTATACGAAAGACCCAAGTTGATAGTCACGGCGTAAAGAGTGGTTAAGGGAACCTATAAATAAAGCCAAATACTTCCCAGGCCGTTGCACGTTTTCTGGAAACACGAAGCCCAATCACGAAAGTAGCTTTATATAAACCACCTGACCCCACGAAAGCTAAGAA |
| Gymnotiformes | *Apteronotus* sp. 8 | 97.6 | 220 | 1 | ACCGCGGTTATACGAAAGACCCAAGTTGATAGTCACTGCGTAAAGAGTGGTTAAGGGAACCAATAAATAAAGCCAAATACTTCCCAGGCCGTTGCACGTTTTCTGGAAACACGAAGCCCAATCACGAAAGTAGCTTTACATAAACCACCTGACCCCACGAAAGCTAAGAA |
| Gymnotiformes | *Apteronotus* sp. 9 | 96.5 | 65 | 2 | ACCGCGGTTATACGAAAGACCCAAGTTGATAGTCACGGCGTAAAGAGTGGTTAAGGGAACCGATGAATAAAGCCAAATACTTCCCAGGCCGTTGCACGTTTTCTGGAAACACGAAGCCCAATCACGAAAGTAGCTTTACATAAACTACCTGACCCCACGAAAGCTAAGAA |
| Gymnotiformes | *Apteronotus* sp. 10 | 95.9 | 19 | 2 | ACCGCGGTTATACGAAAGACCCAAGTTGATAGTCACGGCGTAAAGAGTGGTTAAGGGAGCCGATAAATAAAGCCAAATACTTCCCAGGCCGTTGCACGTTTTCTGGAAACACGAAGCCCAATTACGAAAGTAGCTTTACATAGACCACCTGACCCCACGAAAGCTAAGAA |
| Characiformes | *Boulengerella maculata* | 99.4 | 126 | 1 | ACCGCGGTTAGACGAGTAGGCCCAAGTTGATAGATTACGGCGTAAAGAGTGGTTAAGGATTTTCTCCAAATTAAAGCCAAAGGCCTTCATCGCTGTTATAAGCAGATCCGAAGATCCGAAGCCCATAACGAAAGTAGCTTTATTACCACCTGACCCCACGAAAGCTAAGGA |
| Cichliformes | *Cichla ocellaris* | 100.0 | 52 | 1 | ACCGCGGTTATACGAGAGGCCCAAGTTGACAGACACCGGCGTAAAGAGTGGCTAGGGAAAAATTACTACTAAAGCCGAACACCTTCAGAACTGTCATACGTTCCCGAAGATAAGAAGCCCCACCACGAAAGTGACTTTATATTACCCGACCCCACGAAAGCTGTGAA |
| Cichliformes | *Hypselecara emporalis* | 98.8 | 38 | 1 | ACCGCGGTTATACGAGAGGCCCAAGTTGACAGGTTCCGGCGTAAAGAGTGGTTAAGGAAAAATACTCTACTAAAGCCGAACCCCCTCAGAACTGTTATACGTTTCCGAAGGAATGAAGCCCTACCACGAAAGTGGCTTTACCCTACCCGACTCCACGAAAGCTGCGAAA |
| Cichliformes | *Aequidens* sp. 1 | 96.4 | 2,807 | 5 | ACCGCGGTTATACGAGAGGCCCAAGTTGATAGGCACCGGCGTAAAGAGTGGTTAAGGAAAATATAAACTAAAGCCGAACGCCCTCAGAACTGTTATACGTTTCCGAAGGAAAGAAGCCCCACCACGAAAGTGGCTTTATTCCCCCCGACCCCACGAAAGCTGTGAA |
| Cichliformes | *Aequidens* sp. 2 | 97.8 | 3,031 | 3 | ACCGCGGTTATACGAGAGGCCCAAGTTGACAGGTGCCGGCGTAAAGAGTGGTTAAGGAAAATATAAAATTAAAGCCGAACTCCTTCAGAACTGTCATACGTTTCCGAAGGTAAGAAGCCCTACCACGAAAGTGGCTTTATTCCCCCCGACCCCACGAAAGCTGTGAA |
| Cichliformes | *Apistogramma sp.* | 86.3 | 165 | 3 | ACCGCGGTTATACGAGGGGCCCAAGTTGATAGGTATCCGGCATAAAGGGTGGTTAAGGATATCCCATCACTAAAGCTGAATGCCCCCAAGACTGTCATACGTTTCCGGGGGAAAGAAACCCATAACGAAAGTAGCTTTATCTTACCTGACCCCACGAAAGCTGTGAA |
| Cichliformes | *Astronotus* sp. | 97.6 | 4,444 | 1 | ACCGCGGTTATACGAGAGGCCCAAGTTGACAGATATCGGCGTAAAGCGTGGTTAAGGAAAACTTCTTCACTAAAGCCGAATGCCCTCAGAACCGTCATACGTTCCCGAAGGTAAGAAGCCCCACCACGAAAGTGGCTTTACACTCCCCGACCCCACGAAAGCTGTGAA |
| Cichliformes | *Bujurquina* sp. | 97.0 | 588 | 1 | ACCGCGGTTATACGAGAGGCCCAAGTTGATAGGTACCGGCGTAAAGAGTGGTTAAGGGACATATATAAACTAAAGCCGAACATTTTCAAAGCTGTTATACGCTTCCGAAGATAAGAAGCCCTACCACGAAAGTGGCTTTACTTCCCCCGACTCCACGAAAGCTGCGGA |
| Cichliformes | *Chaetobranchus* sp. | 98.2 | 417 | 1 | ACCGCGGTTATACGAGAGGCCCAAGTTGACAGACACCGGCGTAAAGAGTGGTTAAGGAATCTCAACACTAAAGCCGAATGTTTTCAAGACCGTTATACGTTTTCGAAAATAAGAAGCCCAACCACGAAAGTGGCTTTATTCCACCCGACCCCACGAGAGCTGTGAA |
| Cichliformes | *Crenicichla* sp. 1 | 96.5 | 47 | 2 | ACCGCGGTTATACGAATTGGCCCAAGTTGATAGGTATCGGCGTAAAGCGTGGTTAGGGACTAACCAAAAACTAGAGCCGAACGTCCTCAAAGCTGTTATACGCTTCCGAAAGAACGAAGTCCTCCAACGAAAGTGGCTTTAATACCCAACCTGACCCCACGAAAGCTGTGAAA |
| Cichliformes | *Crenicichla* sp. 2 | 86.0 | 971 | 2 | ACCGCGGTTATACGAATAGGCCCGAGTTGATAAACATCGGCGTAAAGAGTGGTTAGGCATGAGCTAACAACTAAAGCCGAATGTCCTCAAAGCCGTTATACGTTTCCGAAGGAAAGAAGCCCCCTTACGAAAGTAGCTTTAACCTCCAACCTGACCCCACGAAAGCTGAGAA |
| Cichliformes | *Crenicichla* sp. 3 | 88.4 | 312 | 3 | ACCGCGGTTATACGAGCTGGCCCGAGTTGATAAATACCGGCGTAAAGAGTGGTTAAGGGTATACCAAAAACTAAAGCCGAACATCCTCAAGGCCGTTATACGCTTCCGAAGGAGTGAAGACCCCTCACGAAAGTGGCTTTAATATCCAACCTGACCCCACGAAAGCTGTGAA |
| Cichliformes | *Crenicichla* sp. 4 | 87.8 | 6,265 | 8 | ACCGCGGTTATACGAGATGGCCCAAGTTGATAGATGCCGGCGTAAAGGGTGGTTAGGAGTAAATAAAAACTAAAGCCGAACACCCTCAAGACTGTCATACGCTTCCGAAGGAGCGAAGCCCCCCCACGAAAGTGGCTTTAACACCCAACCTGACCCCACGAAAGCTGTGAC |
| Cichliformes | *Crenicichla* sp. 5 | 97.1 | 332 | 5 | ACCGCGGTTATACGAATTGGCCCAAGTTGATAGGTATCGGCGTAAAGCGTGGTTAGGGACTAACCAAAGACTAGAGCCGAACGTCCTCAAAGCTGTTATACGCTTCCGAAAGAACGAAGTCCTCCAACGAAAGTGGCTTTAATACCCAACCTGACCCCACGAAAGCTGTGAA |
| Cichliformes | *Crenicichla* sp. 6 | 98.3 | 148 | 5 | ACCGCGGTTATACGAATTGGCCCAAGTTGATAGGTATCGGCGTAAAGCGTGGTTAAGGATTAACCAAAGACTAAAGCCGAACGTCCTCAAAGCTGTTATACGCTTCCGAAAGAACGAAGTCCTCCAACGAAAGTGGCTTTAATACCCAACCTGACCCCACGAAAGCTGTGAA |
| Cichliformes | *Heros* sp. | 96.5 | 39 | 1 | ACCGCGGTTATACGAGAGGCCCAAGTTGACAGGTGCCGGCGTAAAGAGTGGTTAAGGGAAAACATCCAACTAAAGCCGAACACCTTCAGAGCTGTTATACGTTCCCGAAGGCATGAAGTTCCACCACGAAAGTGGCTTTACCCCCCCCGACCCCACGAAAGCTGTGAA |
| Cichliformes | *Mesonauta* sp. | 97.0 | 1,206 | 1 | ACCGCGGTTATACGAGAGGCCCAAGTTGACAGGTACCGGCGTAAAGAGTGGTTAAGAAAAAATATTAAACTAAAGCCGAACACCTTCAGAACTGTCATACGTTTCCGAAGGCATGAAGCCCCACCACGAAAGTGGCTTTATTTACCCGACCCCACGAAAGCTGTGAA |
| Cichliformes | *Satanoperca* sp. | 97.0 | 14 | 1 | ACCGCGGTTATACGAGAGGCCCAAGTTGATAGACATCGGCGTAAAGAGTGGTTAAGGAAATCCTAAAACTAAAGCCGAATGCCTTCAAGACTGTTATACGTTTCCGAAGGTAAGAAGCCCCACTACGAAAGTGGCTTTACTCCACCTGATTCCACGAAAGCTGTGAAA |
| Synbranchiformes | *Synbranchus* sp. 1 | 94.6 | 340 | 3 | ACCGCGGTTATACAAGAGACTCAATCTGATAACACAACGGCGTAAAGAGTGGTTAAGAAATTCCCAAACTAAGGTCGCACATCTTCATGGTAGTGGTACACATCCGAACATACGAAGCCCATTCACGAAAGTAACCTTACAGATCTGACTCCACGAAAGCTATGGT |
| Synbranchiformes | *Synbranchus* sp. 2 | 94.6 | 24 | 2 | ACCGCGGTTATACAAGAGACTCAAACTGATAACACAACGGCGTAAAGAGTGGTTAAGAAATTCCTAAACTAAGGTCGCACATCTTCATGGTAGTGATACACATCCGAACATACGAAGCCCGTTCACGAAAGTAACCTTACAGATCTGACTCCACGAAAGCTATGGT |
| Pleuronectiformes | *Hypoclinemus mentalis* | 99.4 | 50,324 | 1 | GCCGCGGTTACACGAGAGGTCCAAGTTGATAAACAACGGCGTAAAGGGTGGTTAGGAATAAAAATAAACTAAAGCCGAACGGTTCACAAAGTCATCCTCAAGCTAACGAGAACATGAAGCCCAACCACGAAAGTGGCTTTACATAATTCTGAATCCACAAAAGCTAAGAA |
| Perciformes | *Plagioscion squamosissimus* | 99.1 | 1,220 | 2 | ACCGCGGTTATACGAGAGGCCCAAGTTGATACTCCACGGCGTAAAGAGTGGTTAAAAAAAGACCTATTACTAAAGCCGAACGCCTTCAAAGCTGTTATACGCATCCGAAGGTGAGAAGCCCACCCACGAAAGTGGCTTTACAACCTTGACCCCACGAAAGCTACGAC |
| Perciformes | *Plagioscion* sp. | 96.4 | 1,195 | 1 | ACCGCGGTTATACGAGAGGCCCAAGTTGATACTTCACGGCGTAAAGAGTGGTTAAAAAGACCTATTACTAAAGCCGAACGCCTTCAAAGCTGTTATACGCATCCGAAGGTGAGAAGCCCACCCACGAAAGTGGCTTTACAACCTTGAATCCACGAAAGCTATGAC |
