## Supplementary material for "The critical role of natural history museums in advancing eDNA for biodiversity studies: a case study with Amazonian fishes": Table S5

| **Order** | **Family** | **Genus** | **Species** | **n individuals** |
| --- | --- | --- | --- | --- |
| Characiformes | Acestrorhynchidae | *Acestrorhynchus* | *abbreviatus* | 9 |
| Characiformes | Acestrorhynchidae | *Acestrorhynchus* | *falcirostris* | 1 |
| Characiformes | Acestrorhynchidae | *Acestrorhynchus* | *microlepis* | 1 |
| Pleuronectiformes | Achiridae | *Apionichthys* | *nattereri* | 1 |
| Pleuronectiformes | Achiridae | *Hypoclinemus* | *mentalis* | 1 |
| Characiformes | Chalceidae | *Chalceus* | *erythrurus* | 2 |
| Characiformes | Anostomidae | *Anostomoides* | *laticeps* | 2 |
| Characiformes | Anostomidae | *Leporinus* | *agassizii* | 5 |
| Characiformes | Anostomidae | *Leporinus* | *cylindriformis* | 3 |
| Characiformes | Anostomidae | *Schizodon* | *fasciatus* | 2 |
| Gymnotiformes | Apteronotidae | *Adontosternarchus* | *balaenops* | 1 |
| Gymnotiformes | Apteronotidae | *Adontosternarchus* | *clarkae* | 1 |
| Gymnotiformes | Apteronotidae | *Adontosternarchus* | *sachsi* | 22 |
| Gymnotiformes | Apteronotidae | *Compsaraia* | *compsa* | 1 |
| Gymnotiformes | Apteronotidae | *Porotergus* | *gimbeli* | 1 |
| Gymnotiformes | Apteronotidae | *Sternarchorhynchus* | *cramptoni* | 1 |
| Siluriformes | Auchenipteridae | *Ageneiosus* | *inermis* | 2 |
| Siluriformes | Auchenipteridae | *Ageneiosus* | *lineatus* | 5 |
| Siluriformes | Auchenipteridae | *Ageneiosus* | *ucayalensis* | 1 |
| Siluriformes | Auchenipteridae | *Auchenipterichthys* | *coracoideus* | 1 |
| Siluriformes | Auchenipteridae | *Auchenipterus* | *brachyurus* | 5 |
| Siluriformes | Auchenipteridae | *Auchenipterus* | *nuchalis* | 2 |
| Siluriformes | Auchenipteridae | *Centromochlus* | *existimatus* | 2 |
| Siluriformes | Auchenipteridae | *Epapterus* | *dispilurus* | 1 |
| Siluriformes | Auchenipteridae | *Tatia* | *intermedia* | 9 |
| Siluriformes | Auchenipteridae | *Trachelyopterus* | *galeatus* | 6 |
| Siluriformes | Auchenipteridae | *Tympanopleura* | *brevis* | 21 |
| Siluriformes | Auchenipteridae | *Tympanopleura* | *piperata* | 4 |
| Beloniformes | Belonidae | *Pseudotylosurus* | *microps* | 1 |
| Characiformes | Characidae | *Aphyocharax* | *avary* | 249 |
| Characiformes | Characidae | *Astyanax* | cf. *guianensis* | 7 |
| Characiformes | Bryconidae | *Brycon* | aff. *pesu* | 5 |
| Characiformes | Iguanodectidae | *Bryconops* | *alburnoides* | 1 |
| Characiformes | Characidae | *Ctenobrycon* | *spilurus* | 2 |
| Characiformes | Characidae | *Galeocharax* | *gulo* | 2 |
| Characiformes | Characidae | *Knodus* | *smithi* | 46 |
| Characiformes | Characidae | *Protocheirodon* | *pi* | 1 |
| Characiformes | Characidae | *Moenkhausia* | *cotinho* | 2 |
| Characiformes | Characidae | *Moenkhausia* | *gracilima* | 55 |
| Characiformes | Characidae | *Moenkhausia* | *grandisquamis* | 1 |
| Characiformes | Characidae | *Moenkhausia* | *jamesi* | 8 |
| Characiformes | Characidae | *Moenkhausia* | sp. "red eye" | 1 |
| Characiformes | Characidae | *Paragoniates* | *alburnus* | 6 |
| Characiformes | Characidae | *Prionobrama* | *filigera* | 6 |
| Characiformes | Characidae | *Roeboides* | *affinis* | 3 |
| Characiformes | Characidae | *Roeboides* | *myersii* | 1 |
| Characiformes | Characidae | *Stethaprion* | *erythrops* | 3 |
| Characiformes | Characidae | *Tetragonopterus* | *argenteus* | 6 |
| Characiformes | Triportheidae | *Triportheus* | *albus* | 1 |
| Characiformes | Triportheidae | *Triportheus* | *angulatus* | 4 |
| Characiformes | Triportheidae | *Triportheus* | *auritus* | 1 |
| Characiformes | Chilodontidae | *Caenotropus* | *labyrinthicus* | 1 |
| Cichliformes | Cichlidae | *Biotodoma* | *cupido* | 1 |
| Cichliformes | Cichlidae | *Cichla* | *monoculus* | 1 |
| Cichliformes | Cichlidae | *Crenicichla* | *cincta* | 5 |
| Cichliformes | Cichlidae | *Crenicichla* | *inpa* | 1 |
| Cichliformes | Cichlidae | *Crenicichla* | *reticulata* | 1 |
| Characiformes | Crenuchidae | *Characidium* | aff. *longum* | 2 |
| Characiformes | Crenuchidae | *Melanocharacidium* | *melanopteron* | 1 |
| Characiformes | Ctenoluciidae | *Boulengerella* | *maculata* | 1 |
| Characiformes | Curimatidae | *Curimata* | *roseni* | 1 |
| Characiformes | Curimatidae | *Curimatella* | *dorsalis* | 2 |
| Characiformes | Curimatidae | *Curimatella* | *immaculata* | 2 |
| Characiformes | Curimatidae | *Curimatella* | *meyeri* | 1 |
| Characiformes | Curimatidae | *Steindachnerina* | *bimaculata* | 9 |
| Characiformes | Cynodontidae | *Cynodon* | *gibbus* | 1 |
| Characiformes | Cynodontidae | *Hydrolycus* | *tatauaia* | 1 |
| Characiformes | Cynodontidae | *Rhaphiodon* | *vulpinus* | 2 |
| Siluriformes | Doradidae | *Acanthodoras* | *spinosissimus* | 1 |
| Siluriformes | Doradidae | *Agamyxis* | *pectinifrons* | 5 |
| Siluriformes | Doradidae | *Hemidoras* | *morrisi* | 52 |
| Siluriformes | Doradidae | *Hemidoras* | *stenopeltis* | 12 |
| Siluriformes | Doradidae | *Megalodoras* | *uranoscopus* | 1 |
| Siluriformes | Doradidae | *Nemadoras* | *elongatus* | 3 |
| Siluriformes | Doradidae | *Nemadoras* | *humeralis* | 5 |
| Siluriformes | Doradidae | *Platydoras* | *armatulus* | 2 |
| Siluriformes | Doradidae | *Pterodoras* | *granulosus* | 13 |
| Siluriformes | Doradidae | *Rhynchodoras* | *woodsi* | 9 |
| Siluriformes | Doradidae | *Trachydoras* | sp. | 8 |
| Siluriformes | Doradidae | *Trachydoras* | *steindachneri* | 4 |
| Cupleiformes | Engraulidae | *Anchoviella* | sp. | 5 |
| Characiformes | Erythrinidae | *Hoplias* | *malabaricus* | 1 |
| Characiformes | Gasteropelecidae | *Thoracocharax* | *stellatus* | 7 |
| Characiformes | Hemiodontidae | *Hemiodus* | *microlepis* | 2 |
| Characiformes | Hemiodontidae | *Hemiodus* | *unimaculatus* | 1 |
| Siluriformes | Heptapteridae | *Imparfinis* | *stictonotus* | 1 |
| Siluriformes | Heptapteridae | *Mastiglanis* | sp. | 2 |
| Siluriformes | Heptapteridae | *Pimelodella* | aff. *boliviana* | 1 |
| Siluriformes | Heptapteridae | *Pimelodella* | *boliviana* | 4 |
| Siluriformes | Heptapteridae | *Pimelodella* | *howesi* | 2 |
| Siluriformes | Heptapteridae | *Pimelodella* | *serrata* | 2 |
| Siluriformes | Heptapteridae | *Pimelodella* | sp. | 1 |
| Gymnotiformes | Rhamphichthyidae | *Steatogenys* | *elegans* | 10 |
| Gymnotiformes | Rhamphichthyidae | *Steatogenys* | sp. | 21 |
| Siluriformes | Loricariidae | *Ancistrus* | sp. | 2 |
| Siluriformes | Loricariidae | *Aphanotorulus* | *emarginatus* | 5 |
| Siluriformes | Loricariidae | *Dekeyseria* | *amazonica* | 3 |
| Siluriformes | Loricariidae | *Hypoptopoma* | *gulare* | 6 |
| Siluriformes | Loricariidae | *Hypoptopoma* | *steindachneri* | 3 |
| Siluriformes | Loricariidae | *Hypoptopoma* | *sternoptychum* | 71 |
| Siluriformes | Loricariidae | *Hypoptopoma* | *thoracatum* | 1 |
| Siluriformes | Loricariidae | *Hypostomus* | cf. *plecostomus* | 3 |
| Siluriformes | Loricariidae | *Hypostomus* | *pyrineusi* | 4 |
| Siluriformes | Loricariidae | *Lamontichthys* | *filamentosus* | 23 |
| Siluriformes | Loricariidae | *Lasiancistrus* | aff. schomburgkii | 4 |
| Siluriformes | Loricariidae | *Limatulichthys* | *griseus* | 2 |
| Siluriformes | Loricariidae | *Loricaria* | *cataphracta* | 1 |
| Siluriformes | Loricariidae | *Panaqolus* | *purusiensis* | 13 |
| Siluriformes | Loricariidae | *Panaqolus* | sp. | 1 |
| Siluriformes | Loricariidae | *Peckoltia* | *brevis* | 2 |
| Siluriformes | Loricariidae | *Peckoltichthys* | *bachi* | 4 |
| Siluriformes | Loricariidae | *Pseudacanthicus* | sp. | 3 |
| Siluriformes | Loricariidae | *Pterygoplichthys* | *gibbiceps* | 4 |
| Siluriformes | Loricariidae | *Rineloricaria* | *lanceolata* | 4 |
| Siluriformes | Loricariidae | *Spatuloricaria* | sp. | 19 |
| Siluriformes | Pimelodidae | *Brachyplatystoma* | *juruense* | 1 |
| Siluriformes | Pimelodidae | *Calophysus* | *macropterus* | 1 |
| Siluriformes | Pimelodidae | *Cheirocerus* | *goeldii* | 2 |
| Siluriformes | Pimelodidae | *Hypophthalmus* | *marginatus* | 37 |
| Siluriformes | Pimelodidae | *Leiarius* | *pictus* | 1 |
| Siluriformes | Pimelodidae | *Megalonema* | *amaxanthum* | 1 |
| Siluriformes | Pimelodidae | *Pimelodina* | *flavipinnis* | 1 |
| Siluriformes | Pimelodidae | *Pimelodus* | *blochii* | 2 |
| Siluriformes | Pimelodidae | *Platysilurus* | *mucosus* | 2 |
| Siluriformes | Pimelodidae | *Propimelodus* | *caesius* | 3 |
| Siluriformes | Pimelodidae | *Propimelodus* | sp. | 1 |
| Siluriformes | Pimelodidae | *Sorubim* | *maniradii* | 2 |
| Myliobatiformes | Potamotrygonidae | *Potamotrygon* | *motoro* | 3 |
| Myliobatiformes | Potamotrygonidae | *Potamotrygon* | *scobina* | 2 |
| Characiformes | Prochilodontidae | *Prochilodus* | *nigricans* | 1 |
| Perciformes | Sciaenidae | *Plagioscion* | *squamosissimus* | 28 |
| Characiformes | Serrasalmidae | *Mylossoma* | *aureum* | 1 |
| Characiformes | Serrasalmidae | *Mylossoma* | *duriventre* | 2 |
| Characiformes | Serrasalmidae | *Serrasalmus* | *compressus* | 2 |
| Characiformes | Serrasalmidae | *Serrasalmus* | *rhombeus* | 1 |
| Characiformes | Serrasalmidae | *Serrasalmus* | sp. | 7 |
| Gymnotiformes | Sternopygidae | *Eigenmannia* | *limbata* | 4 |
| Gymnotiformes | Sternopygidae | *Eigenmannia* | *macrops* | 151 |
| Gymnotiformes | Sternopygidae | *Rhabdolichops* | *caviceps* | 11 |
| Gymnotiformes | Sternopygidae | *Rhabdolichops* | *eastwardi* | 24 |
| Gymnotiformes | Sternopygidae | *Rhabdolichops* | *lundbergi* | 10 |
| Gymnotiformes | Sternopygidae | *Rhabdolichops* | *navalha* | 84 |
| Gymnotiformes | Sternopygidae | *Rhabdolichops* | *troscheli* | 1 |
| Gymnotiformes | Sternopygidae | *Sternopygus* | *macrurus* | 3 |
| Siluriformes | Trichomycteridae | *Henonemus* | *punctatus* | 1 |
