## Supplementary material for "The critical role of natural history museums in advancing eDNA for biodiversity studies: a case study with Amazonian fishes": Table S6

| **Order** | **Family** | **Genus_species** | **Frequency** |
| --- | --- | --- | --- |
| Osteoglossiformes | Arapaimidae | *Arapaima gigas* | 4 |
| Cupleiformes | Engraulidae | *Anchoviella* sp. 1 | 4 |
| Cupleiformes | Engraulidae | *Anchoviella* sp. 2 | 1 |
| Cupleiformes | Clupeidae | *Pellona* sp. 1 | 1 |
| Cupleiformes | Clupeidae | *Pellona* sp. 2 | 1 |
| Characiformes | Erythrinidae | *Hoplias* sp. | 2 |
| Characiformes | Cynodontidae | *Cynodon meionactis* | 1 |
| Characiformes | Cynodontidae | *Hydrolycus scomberoides* | 4 |
| Characiformes | Cynodontidae | *Hydrolycus* sp. 1 | 3 |
| Characiformes | Serrasalmidae | *Myleus* sp. | 1 |
| Characiformes | Hemiodontidae | *Anodus* sp. | 3 |
| Characiformes | Anostomidae | *Leporinus apollo* | 1 |
| Characiformes | Anostomidae | *Leporinus* sp. 1 | 1 |
| Characiformes | Anostomidae | *Scheizodon knerii* | 5 |
| Characiformes | Anostomidae | *Leporinus* sp. 4 | 1 |
| Characiformes | Anostomidae | *Leporinus* sp. 5 | 4 |
| Characiformes | Anostomidae | *Leporinus* sp. 6 | 2 |
| Characiformes | Anostomidae | *Leporinus* sp. 11 | 1 |
| Characiformes | Anostomidae | *Megaleporinus* sp. | 1 |
| Characiformes | Curimatidae | *Potamorhina* sp. | 3 |
| Characiformes | Curimatidae | *Cyphocharax* sp. | 1 |
| Characiformes | Prochilodontidae | *Prochilodus harttii* | 5 |
| Characiformes | Prochilodontidae | *Prochilodus* sp. | 2 |
| Characiformes | Prochilodontidae | *Semaprochilodus* sp. 4 | 2 |
| Characiformes | Acestrorhynchidae | *Acestrorhynchus* sp. 1 | 1 |
| Characiformes | Chalceidae | *Chalceus erythrurus* | 3 |
| Characiformes | Chalceidae | *Chalceus macrolepidotus* | 2 |
| Characiformes | Serrasalmidae | *Pygocentrus nattereri* | 4 |
| Characiformes | Bryconidae | *Brycon* sp. 2 | 1 |
| Characiformes | Bryconidae | *Brycon* sp. 5 | 1 |
| Characiformes | Characidae | *Charax* sp. 2 | 2 |
| Characiformes | Characidae | *Charax* sp. 3 | 1 |
| Characiformes | Serrasalmidae | *Colossoma* sp. 1 | 5 |
| Characiformes | Serrasalmidae | *Colossoma* sp. 2 | 1 |
| Characiformes | Characidae | *Corynopoma* sp. | 1 |
| Characiformes | Characidae | *Cyanocharax* sp. 2 | 2 |
| Characiformes | Characidae | *Jupiaba* sp. 1 | 1 |
| Characiformes | Characidae | *Jupiaba* sp. 2 | 2 |
| Characiformes | Characidae | *Moenkhausia* sp. 3 | 1 |
| Characiformes | Serrasalmidae | *Serrasalmus* sp. | 1 |
| Characiformes | Characidae | *Tetragonopterus* sp. | 1 |
| Characiformes | Characidae | *Tyttocharax* sp. | 1 |
| Characiformes | Gasteropelecidae | *Thoracocharax* sp. | 1 |
| Characiformes | Triportheidae | *Triportheus* sp. 2 | 2 |
| Characiformes | Triportheidae | *Triportheus* sp. 3 | 3 |
| Siluriformes | Loricariidae | *Dekeyseria amazonica* | 1 |
| Siluriformes | Loricariidae | *Hypostomus plecostomus* | 1 |
| Siluriformes | Loricariidae | *Lasiancistrus saetiger* | 1 |
| Siluriformes | Loricariidae | *Hypoptopoma* sp. 1 | 1 |
| Siluriformes | Loricariidae | *Hypoptopoma* sp. 2 | 3 |
| Siluriformes | Loricariidae | *Hypoptopoma* sp. 3 | 1 |
| Siluriformes | Loricariidae | *Hypostomus* sp. 1 | 1 |
| Siluriformes | Loricariidae | *Hypostomus* sp. 2 | 1 |
| Siluriformes | Loricariidae | *Pseudacanthicus* sp. | 1 |
| Siluriformes | Loricariidae | *Spatuloricaria* sp. | 1 |
| Siluriformes | Doradidae | *Hemidoras morrisi* | 1 |
| Siluriformes | Doradidae | *Liosomadoras morrowi* | 1 |
| Siluriformes | Doradidae | *Megalodoras uranoscopus* | 1 |
| Siluriformes | Doradidae | *Platydoras costatus* | 2 |
| Siluriformes | Doradidae | *Pterodoras granulosus* | 1 |
| Siluriformes | Auchenipteridae | *Ageneiosus* sp. | 1 |
| Siluriformes | Auchenipteridae | *Auchenipterus* sp. | 1 |
| Siluriformes | Auchenipteridae | *Centromochlus* sp. | 1 |
| Siluriformes | Heptapteridae | *Pimelodella* sp. 1 | 1 |
| Siluriformes | Pimelodidae | *Hypophthalmus edentatus* | 2 |
| Siluriformes | Heptapteridae | *Pimelodella cristata* | 1 |
| Siluriformes | Pimelodidae | *Pseudoplatystoma eticulatum* | 5 |
| Siluriformes | Pimelodidae | *Pimelodus* sp. 1 | 5 |
| Siluriformes | Pimelodidae | *Pimelodus* sp. 2 | 3 |
| Siluriformes | Pimelodidae | *Pinirampus pinirampu* | 1 |
| Siluriformes | Pimelodidae | *Pimelodus* sp. 4 | 5 |
| Siluriformes | Pimelodidae | *Pimelodus* sp. 5 | 1 |
| Siluriformes | Pimelodidae | *Pimelodus* sp. 8 | 1 |
| Siluriformes | Pimelodidae | *Pseudoplatystoma tigrinum* | 2 |
| Siluriformes | Pimelodidae | *Hemisorubim platyrhynchos* | 1 |
| Siluriformes | Pimelodidae | *Brachyplatystoma vaillantii* | 1 |
| Siluriformes | Pimelodidae | *Sorubimichthys planiceps* | 1 |
| Siluriformes | Pimelodidae | *Pseudoplatystoma* sp. | 1 |
| Siluriformes | Pimelodidae | *Sorubim elongatus* | 1 |
| Siluriformes | Pimelodidae | *Sorubim* sp. | 1 |
| Siluriformes | Pimelodidae | *Phractocephalus hemioliopterus* | 4 |
| Siluriformes | Pimelodidae | *Zungaro jahu* | 1 |
| Gymnotiformes | Gymnotidae | *Gymnotus* sp. 1 | 1 |
| Gymnotiformes | Sternopygidae | *Eigenmannia limbata* | 2 |
| Gymnotiformes | Sternopygidae | *Eigenmannia* sp. 1 | 1 |
| Gymnotiformes | Sternopygidae | *Eigenmannia* sp. 6 | 1 |
| Gymnotiformes | Sternopygidae | *Eigenmannia* sp. 8 | 2 |
| Gymnotiformes | Sternopygidae | *Eigenmannia* sp. 9 | 4 |
| Gymnotiformes | Sternopygidae | *Eigenmannia* sp. 13 | 1 |
| Gymnotiformes | Sternopygidae | *Sternopygus* sp. | 4 |
| Gymnotiformes | Apteronotidae | *Apteronotus albifrons* | 1 |
| Gymnotiformes | Apteronotidae | *Apteronotus* sp. 1 | 1 |
| Gymnotiformes | Apteronotidae | *Apteronotus* sp. 3 | 2 |
| Gymnotiformes | Apteronotidae | *Apteronotus* sp. 4 | 3 |
| Gymnotiformes | Apteronotidae | *Apteronotus* sp. 6 | 2 |
| Gymnotiformes | Apteronotidae | *Apteronotus* sp. 7 | 2 |
| Gymnotiformes | Apteronotidae | *Apteronotus* sp. 8 | 1 |
| Characiformes | Ctenoluciidae | *Boulengerella maculata* | 1 |
| Cichliformes | Cichlidae | *Cichla ocellaris* | 1 |
| Cichliformes | Cichlidae | *Aequidens* sp. 2 | 1 |
| Cichliformes | Cichlidae | *Apistogramma sp.* | 1 |
| Cichliformes | Cichlidae | *Crenicichla* sp. 2 | 1 |
| Cichliformes | Cichlidae | *Crenicichla* sp. 3 | 3 |
| Cichliformes | Cichlidae | *Crenicichla* sp. 4 | 3 |
| Cichliformes | Cichlidae | *Mesonauta* sp. | 1 |
| Pleuronectiformes | Achiridae | *Hypoclinemus mentalis* | 1 |
| Perciformes | Sciaenidae | *Plagioscion squamosissimus* | 2 |
