## Supplementary material for "The critical role of natural history museums in advancing eDNA for biodiversity studies: a case study with Amazonian fishes": Table S7

| **Order** | **Family** | **Genus** | **Species** | **n individuals** |
| --- | --- | --- | --- | --- |
| Characiformes | Acestrorhynchidae | *Acestrorhynchus* | *falcatus* | 2 |
| Characiformes | Anostomidae | *Leporinus* | *agassizii* | 10 |
| Characiformes | Anostomidae | *Leporinus* | *cylindriformis* | 2 |
| Characiformes | Anostomidae | *Pseudanos* | *trimaculatus* | 3 |
| Beloniformes | Belonidae | *Potamorrhaphis* | *guianensis* | 6 |
| Siluriformes | Callichthyidae | *Corydoras* | *ambiacus* | 1 |
| Siluriformes | Callichthyidae | *Corydoras* | *arcuatus* | 38 |
| Siluriformes | Callichthyidae | *Corydoras* | *melanistius* | 5 |
| Siluriformes | Callichthyidae | *Corydoras* | *reticulatus* | 2 |
| Characiformes | Characidae | *Astyanax* | aff. *elachylepis* | 3 |
| Characiformes | Characidae | *Astyanax* | *bimaculatus* | 3 |
| Characiformes | Characidae | *Brachychalcinus* | *copei* | 11 |
| Characiformes | Bryconidae | *Brycon* | *amazonicus* | 2 |
| Characiformes | Bryconidae | *Brycon* | *melanopterus* | 14 |
| Characiformes | Iguanodectidae | *Bryconops* | *caudomaculatus* | 6 |
| Characiformes | Iguanodectidae | *Bryconops* | *inpa* | 25 |
| Characiformes | Characidae | *Creagrutus* | *cochui* | 4 |
| Characiformes | Characidae | *Gymnocorymbus* | *thayeri* | 2 |
| Characiformes | Characidae | *Hyphessobrycon* | aff. *heterorhabdus* | 3 |
| Characiformes | Characidae | *Hyphessobrycon* | *agulha* | 2 |
| Characiformes | Characidae | *Hyphessobrycon* | *diancistrus* | 1 |
| Characiformes | Iguanodectidae | *Iguanodectes* | *spilurus* | 13 |
| Characiformes | Characidae | *Moenkhausia* | *chrysargyrea* | 4 |
| Characiformes | Characidae | *Moenkhausia* | *collettii* | 17 |
| Characiformes | Characidae | *Moenkhausia* | *grandisquamis* | 8 |
| Characiformes | Characidae | *Moenkhausia* | *oligolepis* | 4 |
| Characiformes | Characidae | *Moenkhausia* | sp. | 37 |
| Characiformes | Characidae | *Phenacogaster* | aff. *beni* | 2 |
| Characiformes | Characidae | *Phenacogaster* | aff. *pectinatus* | 3 |
| Characiformes | Triportheidae | *Triportheus* | *pictus* | 1 |
| Characiformes | Characidae | *Tyttocharax* | *cochui* | 10 |
| Characiformes | Chilodontidae | *Chilodus* | *punctatus* | 3 |
| Cichliformes | Cichlidae | *Aequidens* | *palidus* | 1 |
| Cichliformes | Cichlidae | *Crenicichla* | *inpa* | 8 |
| Cichliformes | Cichlidae | *Crenicichla* | *regani* | 1 |
| Characiformes | Crenuchidae | *Characidium* | aff. *longum* | 15 |
| Characiformes | Crenuchidae | *Characidium* | aff. *pteroides* | 18 |
| Characiformes | Crenuchidae | *Characidium* | *etheostoma* | 12 |
| Characiformes | Crenuchidae | *Melanocharacidium* | *dispilomma* | 1 |
| Cyprinodontiformes | Rivulidae | *Anablepsoides* | sp. | 18 |
| Cyprinodontiformes | Rivulidae | *Laimosemion* | sp. | 1 |
| Characiformes | Erythrinidae | *Erythrinus* | *erithrynus* | 1 |
| Characiformes | Erythrinidae | *Hoplias* | *malabaricus* | 2 |
| Characiformes | Gasteropelecidae | *Carnegiella* | *strigata* | 1 |
| Gymnotiformes | Gymnotidae | *Gymnotus* | *coropinae* | 1 |
| Gymnotiformes | Gymnotidae | *Gymnotus* | *curupira* | 1 |
| Siluriformes | Heptapteridae | *Brachyrhamdia* | *meesi* | 1 |
| Siluriformes | Heptapteridae | *Chasmocranus* | sp. | 2 |
| Siluriformes | Heptapteridae | *Myoglanis* | *koepckei* | 27 |
| Siluriformes | Heptapteridae | *Pimelodella* | *boliviana* | 8 |
| Gymnotiformes | Rhamphichthyidae | *Hypopygus* | *lepturus* | 2 |
| Characiformes | Lebiasinidae | *Copeina* | *guttata* | 8 |
| Siluriformes | Loricariidae | *Ancistrus* | sp. | 22 |
| Siluriformes | Pimelodidae | *Pimelodus* | *albofasciatus* | 4 |
| Characiformes | Prochilodontidae | *Semaprochilodus* | *insignis* | 4 |
| Gymnotiformes | Rhamphichthyidae | *Gymnorhamphichthys* | *rondoni* | 19 |
