## Supplementary material for "The critical role of natural history museums in advancing eDNA for biodiversity studies: a case study with Amazonian fishes": Table S8

| **Order** | **Family** | **Genus_species** | **Frequencies** |
| --- | --- | --- | --- |
| Characiformes | Crenuchidae | *Characidium* sp. 1 | 2 |
| Characiformes | Crenuchidae | *Characidium* sp. 2 | 5 |
| Characiformes | Crenuchidae | *Characidium* sp. 3 | 2 |
| Characiformes | Crenuchidae | *Characidium* sp. 4 | 2 |
| Characiformes | Crenuchidae | *Elachocharax* sp. 1 | 3 |
| Characiformes | Erythrinidae | *Hoplerythrinus unitaeniatus* | 5 |
| Characiformes | Erythrinidae | *Hoplias malabaricus* | 2 |
| Characiformes | Erythrinidae | *Hoplerythrinus* sp. | 5 |
| Characiformes | Erythrinidae | *Hoplias* sp. | 5 |
| Characiformes | Cynodontidae | *Hydrolycus* sp. 2 | 1 |
| Characiformes | Serrasalmidae | *Serrasalmus eigenmanni* | 1 |
| Characiformes | Anostomidae | *Leporinus apollo* | 2 |
| Characiformes | Anostomidae | *Leporinus fasciatus* | 1 |
| Characiformes | Anostomidae | *Anostomus* sp. | 5 |
| Characiformes | Anostomidae | *Leporinus* sp. 2 | 1 |
| Characiformes | Anostomidae | *Leporinus* sp. 3 | 1 |
| Characiformes | Anostomidae | *Leporinus* sp. 4 | 5 |
| Characiformes | Anostomidae | *Leporinus* sp. 5 | 5 |
| Characiformes | Anostomidae | *Leporinus* sp. 6 | 5 |
| Characiformes | Anostomidae | *Leporinus* sp. 7 | 5 |
| Characiformes | Anostomidae | *Leporinus* sp. 8 | 4 |
| Characiformes | Anostomidae | *Leporinus* sp. 9 | 4 |
| Characiformes | Anostomidae | *Leporinus* sp. 10 | 2 |
| Characiformes | Chilodontidae | *Caenotropus labyrinthicus* | 1 |
| Characiformes | Prochilodontidae | *Semaprochilodus* sp. 1 | 5 |
| Characiformes | Prochilodontidae | *Chilodus* sp. | 2 |
| Characiformes | Prochilodontidae | *Semaprochilodus* sp. 2 | 2 |
| Characiformes | Prochilodontidae | *Semaprochilodus* sp. 3 | 2 |
| Characiformes | Prochilodontidae | *Semaprochilodus* sp. 5 | 2 |
| Characiformes | Acestrorhynchidae | *Acestrorhynchus falcatus* | 5 |
| Characiformes | Acestrorhynchidae | *Acestrorhynchus* sp. 2 | 1 |
| Characiformes | Chalceidae | *Chalceus erythrurus* | 2 |
| Characiformes | Characidae | *Charax pauciradiatus* | 5 |
| Characiformes | Characidae | *Charax* sp. 1 | 1 |
| Characiformes | Characidae | *Ctenobrycon hauxwellianus* | 3 |
| Characiformes | Characidae | *Jupiaba ocellata* | 4 |
| Characiformes | Characidae | *Moenkhausia lepidura* | 3 |
| Characiformes | Serrasalmidae | *Myloplus rubripinnis* | 4 |
| Characiformes | Serrasalmidae | *Pygocentrus nattereri* | 5 |
| Characiformes | Characidae | *Astyanax* sp. 1 | 5 |
| Characiformes | Characidae | *Astyanax* sp. 2 | 5 |
| Characiformes | Characidae | *Astyanax* sp. 3 | 5 |
| Characiformes | Characidae | *Astyanax* sp. 4 | 1 |
| Characiformes | Characidae | *Astyanax* sp. 5 | 1 |
| Characiformes | Bryconidae | *Brycon* sp. 1 | 5 |
| Characiformes | Bryconidae | *Brycon* sp. 2 | 5 |
| Characiformes | Bryconidae | *Brycon* sp. 3 | 5 |
| Characiformes | Bryconidae | *Brycon* sp. 4 | 5 |
| Characiformes | Iguanodectidae | *Bryconops* sp. 1 | 5 |
| Characiformes | Iguanodectidae | *Bryconops* sp. 2 | 5 |
| Characiformes | Characidae | *Charax* sp. 2 | 1 |
| Characiformes | Characidae | *Charax* sp. 4 | 2 |
| Characiformes | Characidae | *Charax* sp. 5 | 1 |
| Characiformes | Serrasalmidae | *Colossoma* sp. 1 | 1 |
| Characiformes | Serrasalmidae | *Colossoma* sp. 2 | 1 |
| Characiformes | Characidae | *Corynopoma* sp. | 1 |
| Characiformes | Characidae | *Cyanocharax* sp. 1 | 1 |
| Characiformes | Characidae | *Hasemania* sp. | 3 |
| Characiformes | Characidae | *Hemigrammus* sp. 1 | 3 |
| Characiformes | Characidae | *Hyphessobrycon* sp. 1 | 1 |
| Characiformes | Characidae | *Hyphessobrycon* sp. 2 | 4 |
| Characiformes | Iguanodectidae | *Iguanodectes* sp. | 3 |
| Characiformes | Characidae | *Jupiaba* sp. 2 | 5 |
| Characiformes | Characidae | *Moenkhausia* sp. 1 | 1 |
| Characiformes | Characidae | *Moenkhausia* sp. 2 | 5 |
| Characiformes | Characidae | *Moenkhausia* sp. 4 | 5 |
| Characiformes | Characidae | *Moenkhausia* sp. 5 | 1 |
| Characiformes | Characidae | *Paracheirodon* sp. | 1 |
| Characiformes | Characidae | *Poptella* sp. | 5 |
| Characiformes | Iguanodectidae | *Bryconops affinis* | 5 |
| Characiformes | Characidae | *Tetragonopterus* sp. | 5 |
| Characiformes | Triportheidae | *Triportheus* sp. 1 | 2 |
| Characiformes | Crenuchidae | *Elachocharax*_sp. 2 | 3 |
| Siluriformes | Cetopsidae | *Helogenes* sp. | 1 |
| Siluriformes | Callichthyidae | *Corydoras* sp. 1 | 4 |
| Siluriformes | Callichthyidae | *Corydoras* sp. 2 | 2 |
| Siluriformes | Loricariidae | *Ancistrus* sp. | 5 |
| Siluriformes | Heptapteridae | *Chasmocranus* sp. | 4 |
| Siluriformes | Heptapteridae | *Imparfinis* sp. | 1 |
| Siluriformes | Heptapteridae | *Pimelodella* sp. 2 | 1 |
| Siluriformes | Heptapteridae | *Rhamdia* sp. | 5 |
| Siluriformes | Heptapteridae | *Pimelodella cristata* | 4 |
| Siluriformes | Pimelodidae | *Hypophthalmus* sp. | 2 |
| Siluriformes | Pimelodidae | *Pimelodus* sp. 3 | 2 |
| Siluriformes | Pimelodidae | *Pimelodus* sp. 7 | 1 |
| Siluriformes | Pimelodidae | *Zungaro* sp. | 5 |
| Siluriformes | Pseudopimelodidae | *Pseudopimelodus pulcher* | 1 |
| Gymnotiformes | Gymnotidae | *Electrophorus varii* | 4 |
| Gymnotiformes | Gymnotidae | *Gymnotus carapo* | 1 |
| Gymnotiformes | Gymnotidae | *Gymnotus* sp. 3 | 4 |
| Gymnotiformes | Gymnotidae | *Gymnotus* sp. 4 | 5 |
| Gymnotiformes | Gymnotidae | *Gymnotus* sp. 5 | 5 |
| Characiformes | Characidae | *Gephyrocharax* sp. 1 | 4 |
| Characiformes | Characidae | *Gephyrocharax* sp. 2 | 4 |
| Gymnotiformes | Rhamphichthyidae | *Gymnorhamphichthys* sp. | 4 |
| Gymnotiformes | Sternopygidae | *Eigenmannia limbata* | 2 |
| Gymnotiformes | Sternopygidae | *Eigenmannia* sp. 1 | 2 |
| Gymnotiformes | Sternopygidae | *Eigenmannia* sp. 2 | 1 |
| Gymnotiformes | Sternopygidae | *Eigenmannia* sp. 3 | 1 |
| Gymnotiformes | Sternopygidae | *Eigenmannia* sp. 4 | 3 |
| Gymnotiformes | Sternopygidae | *Eigenmannia* sp. 7 | 2 |
| Gymnotiformes | Sternopygidae | *Eigenmannia* sp. 8 | 5 |
| Gymnotiformes | Sternopygidae | *Eigenmannia* sp. 9 | 3 |
| Gymnotiformes | Sternopygidae | *Eigenmannia* sp. 12 | 3 |
| Gymnotiformes | Sternopygidae | *Eigenmannia* sp. 14 | 5 |
| Gymnotiformes | Sternopygidae | *Eigenmannia* sp. 15 | 4 |
| Gymnotiformes | Sternopygidae | *Sternopygus* sp. | 1 |
| Gymnotiformes | Apteronotidae | *Apteronotus albifrons* | 2 |
| Gymnotiformes | Apteronotidae | *Apteronotus* sp. 3 | 3 |
| Gymnotiformes | Apteronotidae | *Apteronotus* sp. 4 | 4 |
| Gymnotiformes | Apteronotidae | *Apteronotus* sp. 6 | 2 |
| Gymnotiformes | Apteronotidae | *Apteronotus* sp. 7 | 2 |
| Gymnotiformes | Apteronotidae | *Apteronotus* sp. 9 | 2 |
| Gymnotiformes | Apteronotidae | *Apteronotus* sp. 10 | 1 |
| Cichliformes | Cichlidae | *Hypselecara emporalis* | 5 |
| Cichliformes | Cichlidae | *Aequidens* sp. 1 | 2 |
| Cichliformes | Cichlidae | *Aequidens* sp. 2 | 3 |
| Cichliformes | Cichlidae | *Apistogramma sp.* | 2 |
| Cichliformes | Cichlidae | *Crenicichla* sp. 1 | 1 |
| Cichliformes | Cichlidae | *Crenicichla* sp. 4 | 5 |
| Cichliformes | Cichlidae | *Crenicichla* sp. 5 | 5 |
| Cichliformes | Cichlidae | *Crenicichla* sp. 6 | 5 |
| Cichliformes | Cichlidae | *Heros* sp. | 1 |
| Cichliformes | Cichlidae | *Satanoperca* sp. | 1 |
| Synbranchiformes | Synbranchidae | *Synbranchus* sp. 1 | 3 |
| Synbranchiformes | Synbranchidae | *Synbranchus* sp. 2 | 2 |
