## Supplementary material for "The critical role of natural history museums in advancing eDNA for biodiversity studies: a case study with Amazonian fishes": Table S9

| Family | Species | Ident(%) | LOD | Read | Freq |
| --- | --- | --- | --- | --- | --- |
| Petromyzontidae | *Petromyzon marinus* | 100.0 | HIGH | 65 | 1 |
| Rajidae | *Rajella bigelowi* | 100.0 | MOD | 7 | 1 |
| Cyprinidae | *Acheilognathus cyanostigma* | 100.0 | MOD | 23 | 1 |
| Cobitidae | *Cobitis matsubarai* | 100.0 | MOD | 561 | 1 |
| Macrouridae | *Coelorinchus labiatus* | 100.0 | HIGH | 41 | 1 |
| Gadidae | U98.5 *Microglanis leptostriatus* | 94.8 | LOW | 22 | 1 |
| Gadidae | U98.5 *Molva dipterygia* | 96.4 | LOW | 154 | 1 |
| Odontobutidae | *Odontobutis obscura* | 100.0 | HIGH | 6,806 | 2 |
| Gobiidae | *Chaenogobius gulosus* | 100.0 | HIGH | 1,039 | 1 |
| Gobiidae | *Favonigobius gymnauchen* | 100.0 | HIGH | 81 | 1 |
| Gobiidae | *Gobiopsis macrostoma* | 100.0 | HIGH | 56 | 1 |
| Gobiidae | *Gymnogobius opperiens* | 100.0 | HIGH | 138 | 1 |
| Gobiidae | *Gymnogobius* sp. | 99.4 | LOW | 49 | 1 |
| Gobiidae | *Tridentiger kuroiwae* | 100.0 | LOW | 30 | 1 |
| Mugilidae | *Mugil cephalus* | 100.0 | HIGH | 70 | 1 |
| Atherinidae | U98.5 *Atherina boyeri* gotu1 | 82.0 | HIGH | 257 | 3 |
| Syngnathidae | U98.5 *Syngnathidae* sp. CBM ZF 17997 | 83.7 | LOW | 15 | 1 |
| Mullidae | *Parupeneus barberinus* | 100.0 | HIGH | 1,690 | 1 |
| Siganidae | *Siganus fuscescens* | 100.0 | LOW | 763 | 1 |
| Scorpaenidae | *Paracentropogon rubripinnis* | 99.4 | HIGH | 34 | 1 |
| Platycephalidae | *Platycephalus indicus* | 100.0 | HIGH | 57 | 1 |
| Gasterosteidae | *Pungitius pungitius* | 100.0 | LOW | 57 | 1 |
| Gasterosteidae | *Pungitius sp Brackish* | 99.4 | LOW | 76 | 1 |
| Cottidae | *Cottus amblystomopsis* | 100.0 | HIGH | 106 | 1 |
| Cottidae | *Pseudoblennius marmoratus* | 100.0 | HIGH | 8 | 1 |
